# Optimizing Peptide Crosslinks for Cell-Responsive Hydrogels

**DOI:** 10.1101/2024.05.16.594348

**Authors:** Yingjie Wu, Samuel J. Rozans, Abolfazl Salehi Moghaddam, E. Thomas Pashuck

## Abstract

Cells dynamically modify their local extracellular matrix by expressing proteases that degrade matrix proteins. This enables cells to spread and migrate within tissues, and this process is often mimicked in hydrogels through the incorporation of peptide crosslinks that can be degraded by cell-secreted proteases. However, the cleavage of hydrogel crosslinks will also reduce the local matrix mechanical properties, and most crosslinking peptides, such as the widely used GPQGIWGQ “PanMMP” sequence, lead to bulk degradation of the hydrogel. A subset of proteases are localized to the cell membrane and are only active in the pericellular region in the immediate vicinity of the cell surface. These membrane-type proteases have important physiological roles and enable cells to migrate within tissues. In this work we developed an approach to identify and optimize peptide sequences that are specifically degraded by membrane-type proteases. We utilized a proteomic screen to identify peptide targets, and coupled this with a functional assay that both quantifies peptide degradation by individual cell types and can elucidate whether the peptides are primarily cleaved by soluble proteases or membrane-type proteases. We then used a split-and-pool synthesis approach to generate more than 300 variants of the target peptide to improve the degradation behavior. We identified an optimized peptide sequence, KLVADLMASAE, which is primarily degraded by membrane-type proteases, but enables both endothelial cells and stem cells grown in KLVADLMASAE-crosslinked hydrogels to spread and have viabilities similar to the gels crosslinked by the PanMMP peptide. Notably, the biological performance of the KLVADLMASAE peptide-cross linked gels was significantly improved from the initial peptide target found in the proteomic screen. This work introduces a functional approach to identifying and refining protease-substrate peptides as a way to enhance the properties of hydrogel matrices.

## Introduction

Synthetic hydrogel scaffolds are widely used to model tissues *in vitro* and regenerate them *in vivo*.^1,2^ These gels are often made of crosslinked polymers networks, such as poly(ethylene glycol) (PEG), that offer a high degree of tailorability to the matrix mechanical properties and signaling moieties present in the hydrogels.^3^ Because many polymers do not degrade on the relevant time scales for cell studies, they are frequently crosslinked with peptides that are substrates for cell-secreted proteases, such as the GPQGIWGQ “PanMMP” peptide that is cleaved by many members of the matrix metalloproteinase (MMP) family of proteases.^4,5^ Utilizing protease-cleavable peptides to crosslink hydrogels has the benefit of enabling cell spreading and migration within synthetic matrices (**Fig. 1A**), however they also introduce variability into hydrogel properties since the matrix is actively degraded by cells.

**Fig. 1.**
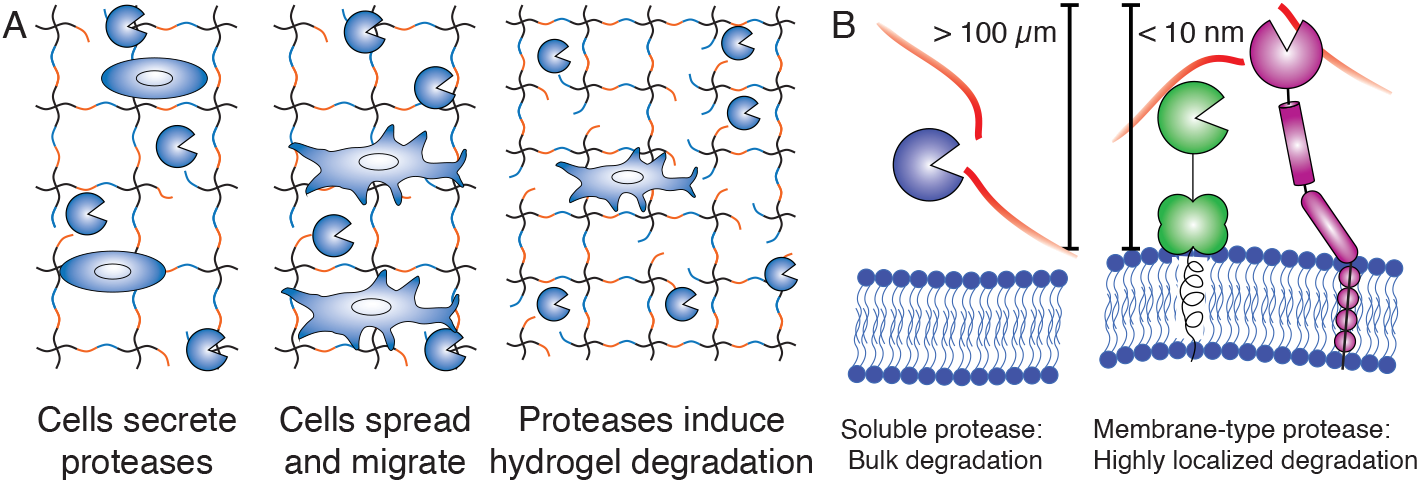
(**A**) Soluble proteases enable cells to spread and migrate within peptide cross linked hydrogels, but can also induce bulk degradation. (**B**) Soluble cell-secreted proteases can diffuse more than 100 μm from the cell surface and induce bulk hydrogel degradation, while membrane-type proteases typically have enzymatic activity located to the cell surface.

Most hydrogels are crosslinked by peptides that are cleaved by soluble proteases that induce bulk hydrogel degradation.^6-8^ The mechanical properties of hydrogels are important for cell behavior^9^ and these properties are dependent on the extent of crosslinking between polymer chains.^6,10^ While synthetic matrices are often treated as relatively inert scaffolds that can be modified with specific ligands or properties, the extent to which the encapsulated cells degrade the hydrogel network will alter the local network structure.^11^ The rate of bulk hydrogel degradation can vary by cell type,^12^ donor age,^13^ and cell seeding density,^14^ which introduces experimental variability in the time-dependent properties of synthetic matrices. A variety of crosslinking peptides have been used to tune how cells degrade the gels, such as increasing or decreasing the rate of bulk hydrogel degradation,^5,15,16^ control cell migration by cell type,^12,17^ or mimic the combine several peptides to mimic the complex environment found in tissues.^18^

Most peptide-cross linked hydrogels are degraded by soluble MMPs which can diffuse over 100 μm from the surface of cells and induce bulk degradation of matrices (**Fig. 1B**).^6,7^ In human tissues protease activity is highly regulated to prevent uncontrolled matrix degradation,^19^ and one mechanism by which cells spatially control degradation is by sequestering proteases to the cell membrane.^20^ These “membrane-type” proteases are involved in numerous physiological processes, and members of this family of proteases, such as MMP-14 (also called MT1-MMP) play important roles in cell migration *in vivo*,^21^ while only degrading the pericellular matrix in the immediate vicinity of the cell surface.^22^ Identifying peptides that are primarily cleaved on the cell surface, rather than by soluble proteases, is a promising method for maintaining hydrogel properties consistent across different cell types, culture periods, and seeding densities, while enabling cell spreading and migration within the matrix. Hydrogels have been crosslinked with peptides that are substrates for membrane-type proteases.^12,15^ However, cell express a complex mixture of proteases^19^ and most MMP peptides, including the PanMMP peptide, are cleaved by numerous members of the MMP family.^5,23^ As a result of this non-specificity, most peptides that are cleaved by membrane-type proteases are also cleaved by soluble proteases, limiting their ability to reduce bulk hydrogel degradation.

In this work we used a proteomics-based approach to identify novel protease-substrate peptides. We then used a functional cell-based degradation assay to select a peptide which is primarily degraded in cell culture only when cells are present, but not in cell-conditioned media that contains soluble proteases. These peptides were candidates for membrane-specific degradation. We then utilized as split-and-pool synthesis scheme to make 361 variants of the most promising membrane-specific peptide to improve the degradation kinetics while maintaining specificity for degradation when cells are present. The optimized peptides were used as crosslinks for hydrogels, and we determined that a KLVADLMASAE peptide enables both human mesenchymal stem cells (hMSCs) and human umbilical vein endothelial cells (hUVECs) to spread in the gels with average cell area and viability equivalent to the PanMMP peptide, but significantly improved over the initial peptide identified in the proteomics screen.

## Results and Discussion

### Proteomics-based approach to peptide discovery

We utilized a proteomics-based technique to identify protease substrate peptides and then developed a novel functional assay to quantify how quickly these peptides are degraded. Terminal amine isotopic labeling of substrates (TAILS) is proteomics technique in which the sites of proteolytic degradation of proteins are labelled through dimethylation of N-terminal amines, which is a chemical modification that is not found in nature.^24,25^ These dimethylated N-termini can then be identified in liquid chromatography mass spectrometry (LCMS) and are candidates for sites of proteolytic cleavage. Candidate peptides found in the TAILS screen were then synthesized with the 4-5 amino acids flanking the N- and C-terminal sides of the likely site of proteolytic cleavage, and modified with an acetylated β-alanine on the N-terminus and an amidated β-alanine on the C-terminus to prevent non-specific degradation from exopeptidases.^26^

Most *in vitro* protease activity assays incubate the peptides or proteins of interest with a single protease in a buffered solution. However, the kinetics of protease activity in tissues are complex and highly regulated, and both the activity^27^ and substrate specificity^28^ of proteases can be modulated through association with other proteins.^29^ Most MMP-substrate peptides are also cleaved by multiple proteases,^5^ and within the context of biomaterials the most important factor is typically how the combined activity of all proteases modifies the local microenvironment versus the activity due to any individual enzyme. We synthesized peptides identified in the TAILS proteomic screen, pooled them, and added them either to cell culture media on top of cells, the “Cells” condition, or conditioned media that does not have cells, the “Media” condition (**Fig. 2A**). The fraction of peptides remaining after 24 hours in each condition was then quantified using LCMS. Soluble proteases should be present in both conditions, however membrane-type proteases should only be found in the samples containing cells and not the conditioned media samples. A key benefit of this approach is that is surveys functional proteolytic activity of all proteases, including the effects of inhibitors and co-factors, and is able to quantify how quickly soluble peptides are degraded by all of the proteases secreted by different types.

**Fig. 2.**
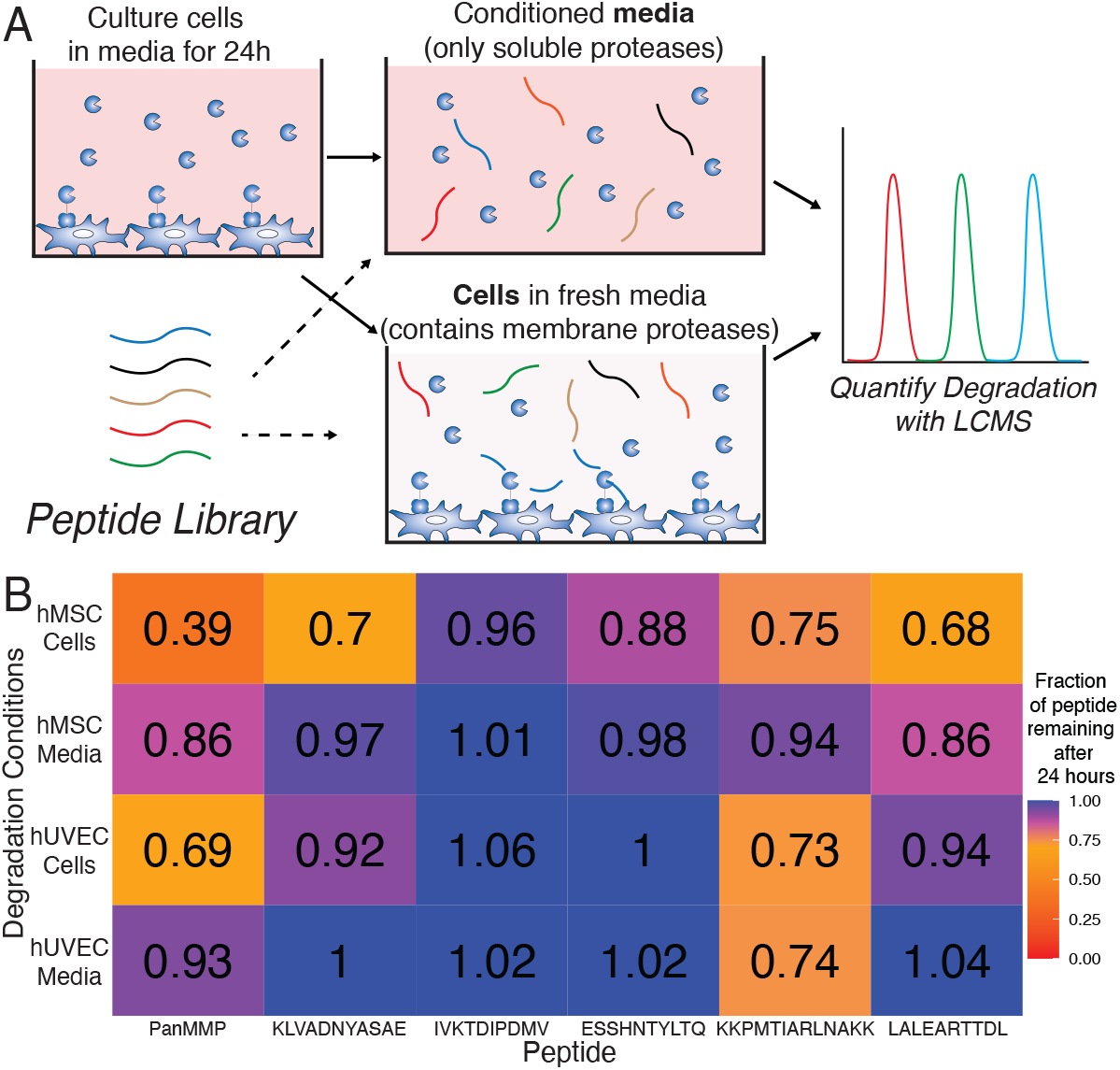
(**A**) We used a functional approach to quantify how peptides are degraded by both cells in media (Cells) and conditioned media (Media). Since membrane-type proteases should only be found on cells, peptides which are primarily degraded in the Cells condition are candidates for relativity specificity to membrane-type proteases. (**B**) We found that the commonly-used PanMMP showed degradation in both Cells and Media conditions, while the KLVADNYASAE peptide showed minimal degradation in the Media condition.

We tested our initial peptide library on both hMSCs and hUVECs and found that peptides had varying degradation rates by both cell type and cell/media condition. In **Fig. 2B** our data shows that some peptides, such as the IVKTDIPDMV sequence, had minimal degradation under any conditions, while both the KKPMTIARLNAKK peptide and PanMMP peptide were significantly degraded by both hMSCs and hUVECs in both the Cells and Media conditions. The KLVADNYASAE peptide was identified as a peptide sequences which was cleaved by both hMSCs and hUVECs, but was primarily cleaved only when cells are present and not in conditioned media, having less than 3% degradation in media condition for either cell type.

While the KLVADNYASAE peptide showed more than 5% degradation in the Cells condition for both hMSCs (30% degradation) and hUVECs (8% degradation), the amount of degradation in the Cell condition was much lower than the PanMMP peptide for both hMSCs (61% for the PanMMP peptide versus 30% for the KLVADNYASASE) and hUVECs (31% for the PanMMP peptide versus 8% for the KLVADNYASAE peptide). Since the final goal of these peptides is to crosslink hydrogels and enable cells to spread and migrate, it is desirable to have peptides that are cleaved under the Cells condition as rapidly as possible to ensure that the spreading and migration processes are not hindered by the inability of cells to degrade the local matrix. The rate at which a specific peptide is degraded by aproteaseis highly sensitive to the amino acids near the cleavage site,^30^ and mutating amino acids near this site can significantly change the rate of protease activity.^23^

### Split-and-pool synthesis of peptide libraries to improve protease kinetics

Our proteomics-based method for identifying peptide sequences is limited to the protease-accessible parts of proteins secreted by cells, which contains just a minuscule fraction of the ∼10^13^ possible 10 amino acid sequences. We used a “split-and-pool” approach^31^ to synthesize more than 300 peptide variants of the KLVADNYASAE peptide first identified in our proteomics screen (**Fig. 3A**). Previous work has shown that enzyme kinetics of membrane-type MMPs, including MMP-14, are highly sensitive to the amino acids near the site of cleavage.^32,33^ We mutated the P1’ and P2’ locations on the peptide, which are the two amino acids on the C-terminal site of cleavage, having the format KLVADX_1_X_2_ASAE. Peptides were synthesized on solid-phase resin consisting of 10^5^-10^6^ particles per gram of resin and upon addition of the ASAE amino acids the resin was split 19 ways. Each of the canonical amino acid except cysteine was added to one of the 19 fractions. These were then pooled together into a single batch containing all 19 variants, and then split 19 ways again to create 361 (=19^2^) variants having the sequence KLVADX_1_X_2_ASAE, where X_1_ and X_2_ can be each of the 19 different amino acids. The second split of 19 different variants was then pooled into 10 different libraries and the remaining KLVAD amino acids were added to each of the libraries. The ten peptide libraries containing either 19 or 38 different peptides that were incubated with hUVEC and hMSC in the Cells and Media conditions and the degradation of each peptide in the libraries was quantified using LCMS.

**Fig. 3.**
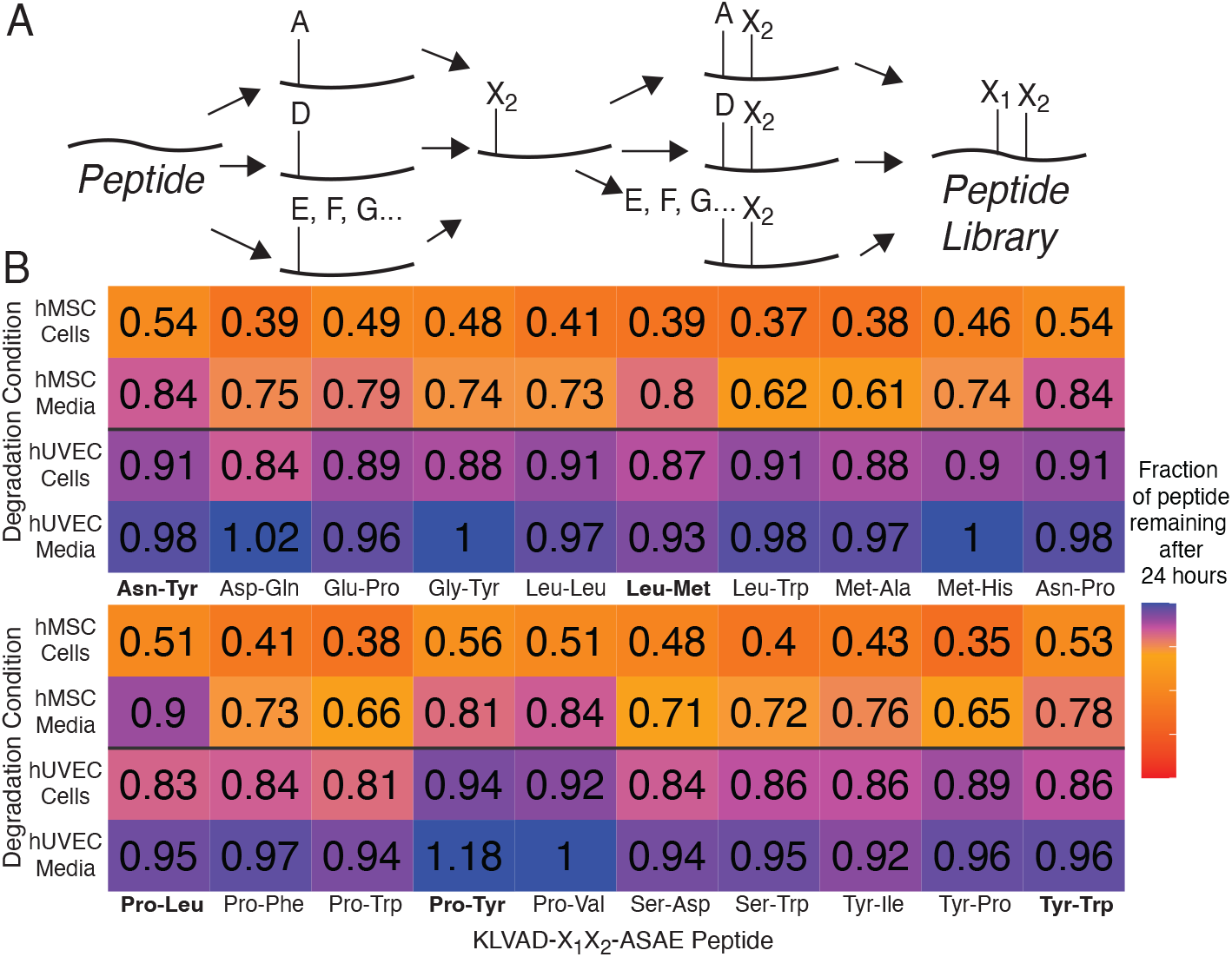
(**A**) A split-and-pool approach was used to tune peptide degradation kinetics by making 361 variants of the KLVADNYASAE peptide having the structure KLVADX_1_X_2_ASAE. (**B**) Modifying the amino acids near the site of proteolytic cleavage tuned the degradation kinetics for both hMSCs and hUVECs (N = 2).

The peptides in the libraries were evaluated with two primary considerations: having maximal degradation in the “Cells” condition that contains membrane-proteases, and minimal degradation in the “Media” condition which should not have membrane-type proteases. An ideal crosslinking peptide should also be degraded by both hMSCs and hUVECs, like the PanMMP peptide so that it can be used across different cell types. Minimizing degradation in the Media condition is important to limit unwanted bulk hydrogel degradation. **Fig. 3B** shows the subset of 361 peptide variants which had minimal degradation in hUVEC Media condition (>93% remaining after 24 hours), but had significant degradation in the hMSC Cells condition (<54% of peptide remaining after 24 hours), and also included the Asn-Tyr peptide from KLVAD-**NY** -ASAE, the peptide sequence from which the split-and-pool library was derived. Peptides were further refined by selecting for those which had minimal degradation in hMSC media (>77% of peptide remaining). From this smaller pool the following peptides were selected, KLVAD-**LM**-ASAE, KLVAD- **PL** -ASAE, PLVAD- **PY** - ASAE, and KLVAD-**YW**-ASAE, in addition to the parent peptide, KLVAD-**NY**-ASAE for further characterization.

The most promising peptides identified in the split-and-pool study were then synthesized using standard peptide synthesis protocols and cultured with hMSCs and hUVECs at a higher concentration of 50 μM in both the Cells and Media condition to validate their degradation kinetics. As can be seen in **Fig. 4**, the KLVAD-derived peptides had less than 10% degradation in the cell condition, and in most cases had less than 5% degradation. Utilizing the split-and-pool approach was successful in improving the degradation kinetics of the peptide identified in the proteomic screen. For instance, the KLVAD-**LM** peptide had increased degradation compared to the KLVAD-**NY** peptide in the cells condition for both hMSCs (54% degradation versus 22% degradation and hUVECs (16% degradation versus 4% degradation) while also having reduced degradation in the Media conditions. The PanMMP peptide show significant cleavage in the Media condition which contains soluble proteases for both hMSCs and hUVECs. These results indicate that the an initial proteomic peptide screen was able to identify a peptide with the desired degradation specificity and that the subsequent split-and-pool screen improved peptide degradation under desired conditions while minimizing degradation in undesirable conditions.

**Fig. 4.**
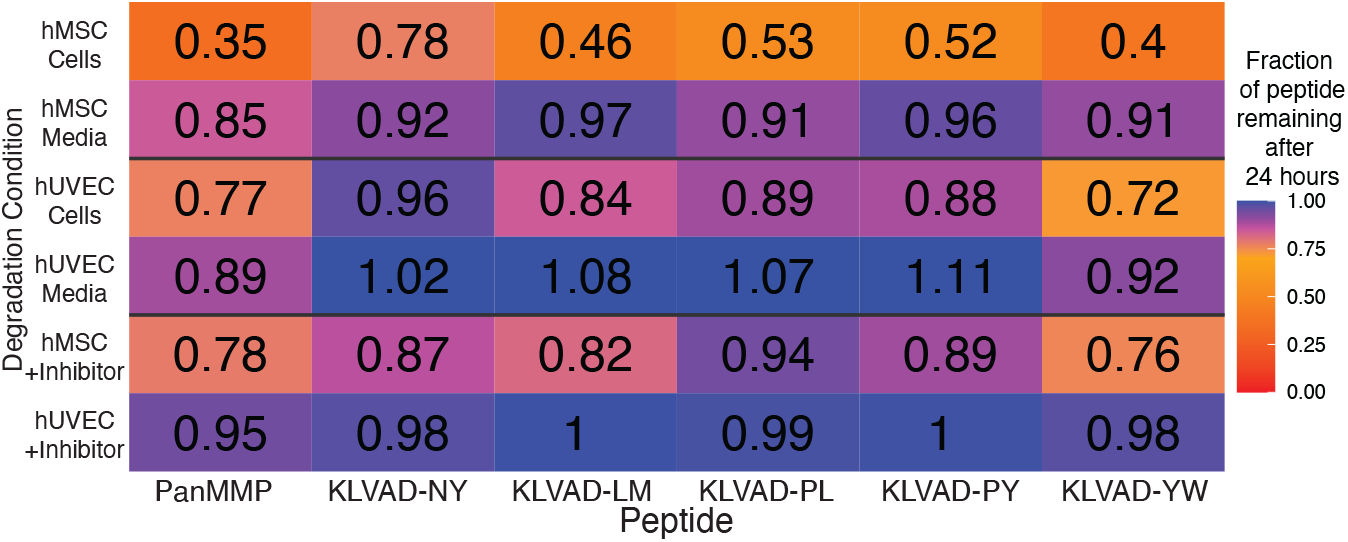
The most promising peptides from the split-and-pool studies were cultured hMSCs and hUVECs and the amount of peptide degradation in both the Cells and Media was quantified using LCMS. Peptides were also added to the Cells condition that contained an MMP-14 inhibitor. KLVAD peptides showed significant specificity for degradation in the Cells condition, and the addition of an MMP-14 inhibitor greatly reduced the peptide degradation rate.

A key benefit of a functional approach to quantifying degradation during culture is that the kinetics of degradation can be tuned without any prior knowledge of protease expression profiles. A consequence of this approach is that the identity of the specific protease(s) that are cleaving the peptides is completely unknown. The membrane protease MMP-14 is a widely-expressed across cell types, including both hMSCs^34,35^ and hUVECS,^36,37^ where it has roles in migration, differentiation,^34,35^ and angiogenesis.^36,37^ This makes MMP-14 a likely candidate for catalyzing peptide degradation in the presence of cells. We added the MMP-14 inhibitor NSC 405020^38^ to the Cells condition and saw a reduction in protease degradation of all peptides across both cell types (**Fig. 4**). Interestingly, while most peptides still had greater than 10% degradation in the Cells + Inhibitor condition for hMSCs, only the PanMMP had greater than 2% degradation after 24 hours when cultured with hUVECs. These results indicate that MMP-14 is possibly responsible for almost all peptide degradation in the hUVEC Cells condition, while the partial reduction of proteolytic degradation in the hMSC Cells condition suggests that both MMP-14 and other proteases are responsible for peptide degradation. It also notable that the MMP-14 inhibitor reduced PanMMP degradation in hMSCs from 65% degradation to 22%, and in hUVECs from 15% degradation to 5%, indicating that MMP-14 plays a major role in cell-based degradation of the PanMMP sequence, even though it is known to be cleaved by many other MMPs. This indicates that while both peptides are cleaved by more than one protease, a larger fraction of protease activity against the KLVAD peptides is due the membrane-type MMP-14 protease. However, it should be noted that MMP-14 has been shown to be involved in the proteolytic activation of MMP-2^39^ and other MMPs.^40,41^ So while these results suggest that inhibiting MMP-14 reduces proteolytic cleavage, it is possible that other proteases are ultimately responsible for cleaving the peptides.

### Cell growth and viability within peptide-cross linked hydrogels

Each of the peptides was then used to crosslink hydrogels containing hMSCs or hUVECs at 20 million cells/mL and cultured for 1, 3, or 7 days. We found that both the PanMMP peptide and KLVAD-**LM** supported cell spreading and growth within the hydrogels (**Fig. 5**). Interestingly, hydrogels crosslinked by the original KLVAD-**NY**-ASAE peptide was found to be cytotoxic, and by Day 7 there were no detectable cells within the hMSC-seeded gels, and hUVECs showed only minimal cell spreading. The KLVAD-**PY** peptide also had less cell spreading and viability compared to the PanMMP and KLVAD-**LM** peptides for both hMSCs and hUVECs (**Fig. 5 I-J**). Interestingly, the addition of an MMP-14 inhibitor completely reduced the ability of hUVECs to increase their spreading in from Day 1 to Day 7 in a KLVAD-**LM** crosslinked hydrogel, but hUVECs in a PanMMP-crosslinked hydrogel were still able to spread (**Fig. S1-2**). This supports the inhibitor studies for the soluble peptides in which the MMP-14 inhibitor completely abolished degradation of the KLVAD-**LM** peptide by hUVECs, but not the PanMMP peptide. (**Fig. 4**).

**Fig. 5.**
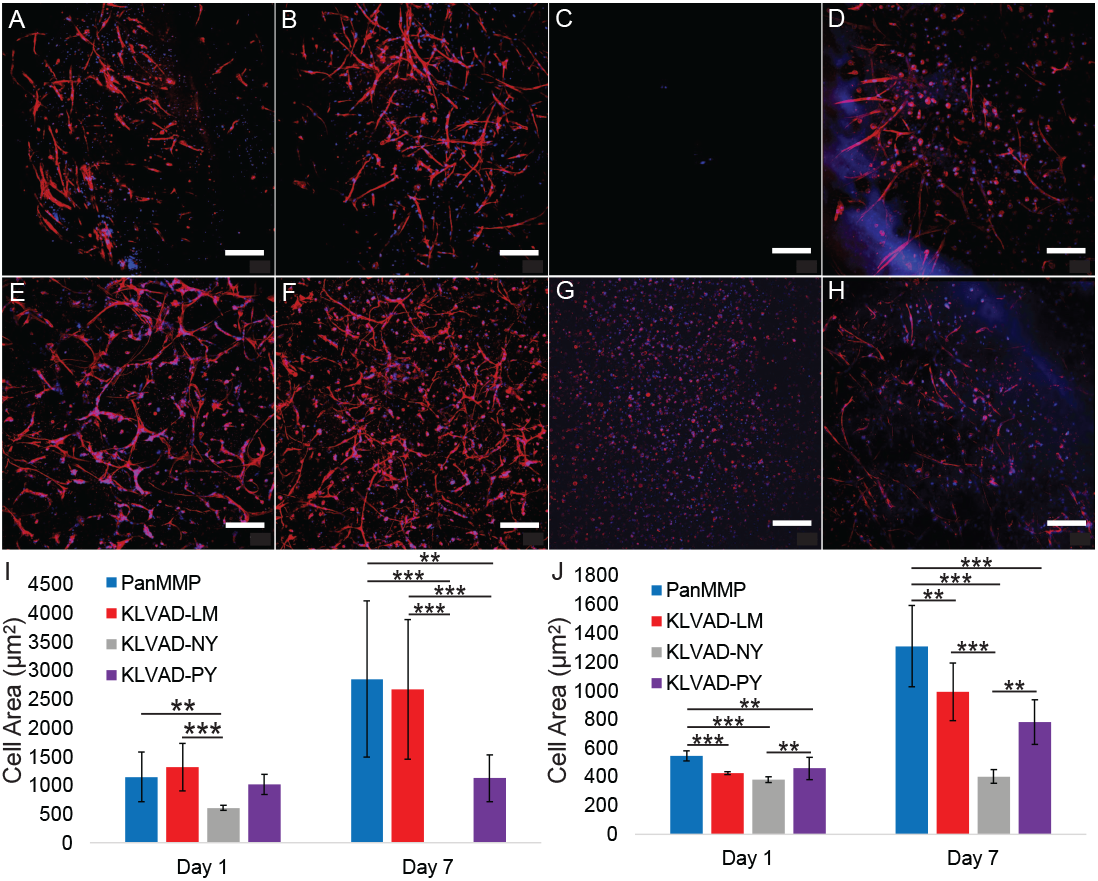
Cells were grown for seven days in PEG hydrogels with different crosslinking peptides. hMSCs in hydrogels crosslinked by the (**A**) PanMMP, (**B**) KLVAD-LM, (**C**) KLVAD-NY, and (**D**) KLVAD-PY peptides, and hUVECs cultured in hydrogels crosslinked by the (**E**) PanMMP, (**F**) KLVAD-LM, (**G**) KLVAD-NY, and (**H**) KLVAD-PY peptides. The average cell area for was quantified at 1 and 3 days for (**I**) hMSCs and (**J**) hUVECs. Scale bar is 100 μm. * indicates p < 0.05, ** indicates p < 0.01, and *** indicates that p < 0.001 by Tukey’s host-hoc test.

Cell viability was also found to be highly dependent on the sequence of the peptide crosslinking the PEG hydrogels. hMSCs cultured in hydrogels crosslinked by either the either the PanMMP or KLVAD-**LM** peptide had significantly higher viability after seven days of culture than the KLVAD-**NY** or KLVAD-**PY** peptide by either metabolic activity assays (**Fig. 6A**) or cell number (**Fig. 6b**). Notably, the KLVAD-**LM**-ASAE peptide is not found within the human proteome and thus cannot be identified through a proteomics-only approach, and this highlights the importance of utilizing a split-and-pool methodology for improving the biological performance of crosslinking peptides. hUVECs cultured in gels crosslinked by the PanMMP peptide had higher metabolic activity after seven days than the KLVAD peptides (**Fig. 6C**), although the difference in cell number was not significant (**Fig. 6D**). hUVEC growth is optimized in very weak highly degraded hydrogels,^42^ so the propensity of the PanMMP peptide to undergo bulk degradation could be advantageous for this cell type.

**Fig. 6.**
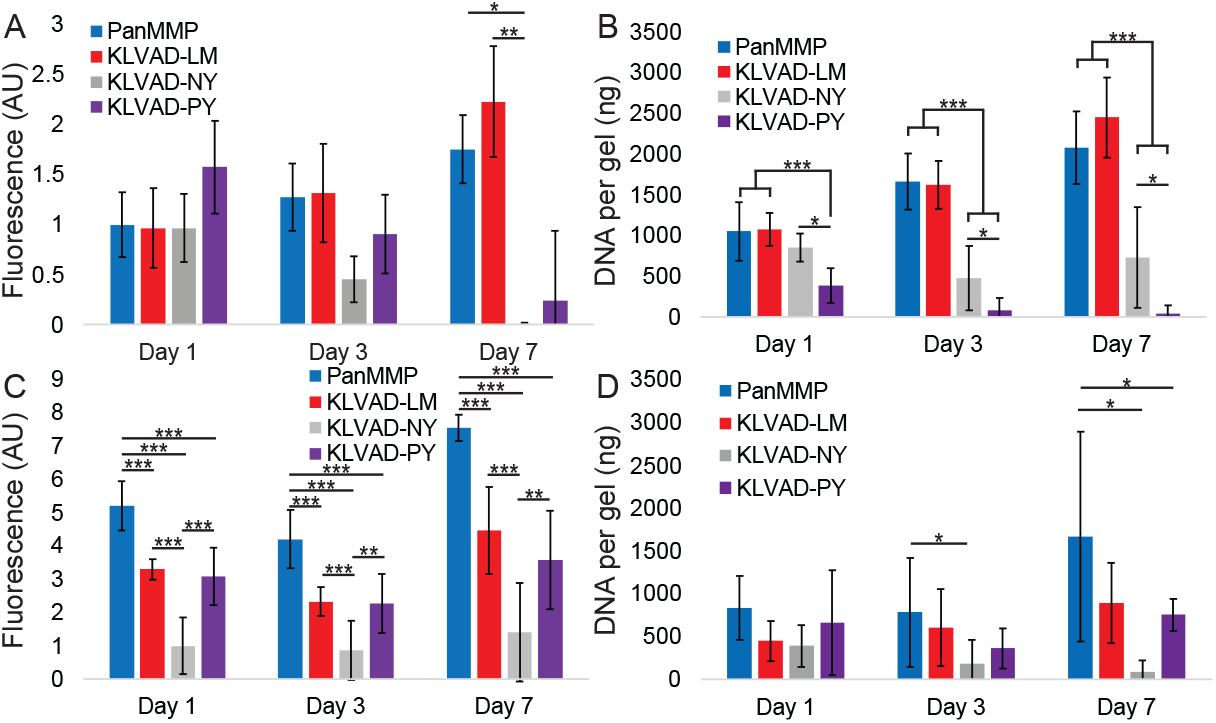
The sequence of crosslinking peptides influenced cell viability within hydrogels. The Pan-MMP and KLVAD-LM peptides had higher hMSCs viability as determined by both (**A**) alamarBlue viability and (**B**) DNA quantification, while the viability for hUVECs had less dependence on peptide for both alamarBlue (**C**) and DNA quantification (**D**). * indicates p < 0.05, ** indicates p < 0.01, and *** indicates that p < 0.001 by Tukey’s host-hoc test.

## Conclusions

In conclusion, we utilized an approach combining proteomics, functional degradation screens, and split-and-pool synthesis to identify and refine peptides to control cell-matrix interactions within synthetic hydrogels. Through the use of a split-and-pool approach used to generate larger peptide libraries we identified a KLVAD-**LM**-ASAE peptide sequence that is less degraded by soluble proteases than the commonly used GPQGIWGQ PanMMP peptide, but still shows significant degradation in the presence of cells. Compared to the KLVAD-**NY**-ASAE peptide which was initially identified in the proteomics screen, the KLVAD-**NY**-ASAE peptide had increased cell spreading and viability for hMSCs and hUVECs. Since protease activity is altered in a range of developmental, regenerative, and degenerative processes, this approach has the potential to improve our ability to harness cell-matrix interactions for a therapeutic benefit.

## Acknowledgements

We would like to acknowledge our funding sources, the NIH (1R21GM143593-01) and NSF (award 2138723). We are grateful to the lab of Lesley Chow for the use of their preparative high performance liquid chromatography.

## Materials

All peptide synthesis reagents were purchased from Chemscene or Ambeed. N,N-Dimethylformamide and dichloromethane (both from VWR BDH Chemicals), piperidine (Millipore Sigma), trifluoroacetic acid (Millipore Sigma), diethyl ether (Fisher Scientific) were used as purchased. 20 kDa 8-arm poly(ethylene glycol) dibenzocyclooctyne (PEG-DBCO) was purchased from Biopharm PEG. Batches which dissolved in water in under 30 seconds were used as received. Otherwise they were quickly purified by dissolving them in isopropanol and removing impurities using a 10 kDa Amicon centrifugal filter. Approximately 200 mg of PEG was dissolved in 20 mL of isopropanol and it was centrifuged at 4,500 RPM until there was less than one milliliter of PEG-DBCO/isopropanol left in the filter. This was repeated twice, and then the filtrate was lyophilized and used.

## Methods

### Peptide Synthesis Procedure

Peptides were synthesized using standard solid phase peptide synthesis protocols using either manual synthesis or an automated peptide synthesizer (CEM Liberty Blue) using standard Fmoc-protected amino acids (Chemscene) on a Rink amide resin (Supra Sciences) unless otherwise noted. All amide couplings were done using O-(6-chlorobenzotriazol-1-yl)-N,N,N′,N′-tetramethyluronium hexafluorophosphate (HCTU) unless otherwise noted. For each coupling the amino acid, HCTU, and DIPEA were added in a 4:4:6 ratio to the peptide. During peptide synthesis a ninhydrin test was performed after every addition to test for the presence of free amines. Upon a positive test, the coupling was replicated until the test was negative. A capping step was then performed with acetic anhydride (Sigma-Aldrich) in a 10:5:100 acetic anhydride:DIPEA:DMF solution twice for 5 min, and then a ninhydrin test was performed to check for complete capping of the free amines. After successful coupling, the Fmoc group was removed, washing the resin with 20% piperidine in DMF twice for 5 min. A ninhydrin test was performed to check for a positive result.

For split-and-pool steps the resin was washed 3x with DMF and then the entire amount of resin was weighed on a scale. This total mass of resin was divided by 22 and this amount was added into each of into 19 separate tubes. Twenty two was used to account for resin loss during transport and weighing. The reactions were performed in 15 mL tubes and upon successful coupling of all 19 amino acids all 19 fractions of resin were re-combined and then split 19 ways again using the same protocol. To reduce the complexity of the individual libraries, these 19 fractions were combined into 10 different libraries.

Peptides libraries and all peptides containing tryptophan were cleaved using 92.5% trifluoroacetic acid (TFA), 2.5% H_2_O, 2.5% triisopropylsilane (TIPS), and 2.5% dithiothreitol (DTT). Peptides which lacked a tryptophan were cleaved without DTT. Peptides were typically cleaved for 2-3 hours at room temperature using approximately 2 mL of cleavage solution per mM of peptide. however peptides containing azides were cleaved for 30 minutes to prevent degradation of the azide group. The mass was checked using electrospray ionization and if protecting groups remained the peptide was re-cleaved for 30 minutes. At the end of the cleavage the peptides were precipitated in diethyl ether. These were then centrifuged for 5 minutes at 4,000 rpm, and the supernatant was discarded. The peptide pellet was washed with diethyl ether, and centrifuged, and this was repeated twice. The peptide pellet was allowed to dry, and then dissolved in water and neutralized with ammonium hydroxide prior to purification.

The cyclic RGD peptide was synthesized on a 2-chlorotrityl chloride resin. The first amino acid (0.3 mmol of amino acid per gram of resin) was dissolved in dichloromethane (DCM) and added to the resin in a shaker vessel. Five equivalents of DIPEA was then added, and after 5 min of shaking another 1.5 equiv of DIPEA was added. After 1 h the unreacted 2-chlorotrityl chloride resin was capped with an excess of methanol for 30 min. The rest of the amino acids were then coupled using standard solid phase Fmoc-synthesis protocols. After the last Fmoc group was deprotected the resin was washed 3× in DMF and 3× in DCM. The peptide was then cleaved under mild acidic conditions consisting of 5% trifluouroacetic acid (TFA) and 2.5% triisopropylsilane (TIS) in DCM. The mild cleavage solution was added to the resin for 5 minutes and then collected into a round bottom flask and this was repeated until the resin turned dark red or black. The collected liquid was then precipitated in diethyl ether. This was then dissolved in 50:50 acetonitrile:water, neutralized with 1M NH_4_OH, and lyophilized. The protected linear GRGDSK(N_3_) peptide was then cyclized. This was done by dissolving the peptide into DMF at 1 mg/mL and adding 3 equivalents of (O-(7-azabenzotriazol-1-yl)-N,N,N’,N’-tetramethyluronium hexafluorophosphate) (HATU). After six hours the DMF was removed using rotary evaporation at 75°C.

All peptides were purified using HPLC using a Phenomenex Gemini 5 μm NX-C18 110 Å LC Column 150×21.2 mm. Gradients were run from 95% Mobile Phase A (water with 0.1% TFA) and 5% Mobile Phase B (acetonitrile with 0.1% TFA) to 100% Mobile Phase B. A typical HPLC run featured a two minute equilibration step, followed by a 10 minute ramp from 95% Mobile Phase A to 100% Mobile Phase B, and then two minutes of equilibration at 100% Mobile Phase B, before ramping back down to the starting conditions. Notably, the split-and-pool libraries were ramped up to 100% Mobile Phase B over two minutes, since these libraries consisted of approximately 38 different peptides which weren’t intended to be separated from each other. The protected cyclic RGD peptides was purified using a gradient that ramped from 30% Mobile Phase B to 100% Mobile Phase B. After purification all peptides were lyophilized and were ready to use, except for the protected cyclic RGD peptide which was cleaved using 95% TFA, 2.5% TIPS, and 2.5% H_2_O for one hour.

### General Cell Culture Procedure

Human mesenchymal stem cells (hMSCs) (Rooster Bio, Donor 310305, a 20 year old African American female) and human umbilical vein endothelial cells (hUVECs) (Lifeline Cell Technology, Donor 04608, an African American male) were cultured according to the manufacturers instructions using RoosterNourish hMSC media or Lifeline VascuLife EnGS hUVEC media.

### Peptide Degradation Study Procedure

hUVECs (75,000 per well) or hMSCs (36,000 per well) were seeded into a 48 well plate and 500 μL media was added to the wells. The cells were cultured for 24 hours, at which point the conditioned media was transferred into a new plate that does not contain cells and 500 μL fresh media was added into the wells with cells. The peptide libraries were dissolved at a stock solution of 10 mM total peptide content. 95 μL of 38 peptide library and 47.5 μL for 19 peptide library peptide solution was added to each well and media was taken at 0 and 24 hours. Add 4 μL of acetic acid to all collected media in the LC-MS plates and store them in a -80 °C freezer to prevent further proteolytic degradation. From each sample, 10 μL of crude solution was introduced by the LC-MS through an Thermo Scientific™ Vanquish™ LC System (Thermo Fisher Scientific) which outputted to a Thermo Scientific™ LTQ XL™ Linear Ion Trap Mass Spectrometer (Thermo Fisher Scientific). The sampled mixture was trapped on a column (ProntoSIL C18 AQ, 120 Å, 3 μm, 2.0 × 50 mm HPLC Column, PN 0502F184PS030, MAC-MOD

Analytical Inc.). The samples were loaded onto the column with a solvent containing acetonitrile/ water, 5:95 (v/v) containing 1% Acetic Acid at a flow rate of 300 μL/min and held for one minute. The sample was then eluted from the column with a linear gradient of 5-40% Solvent B (1% Acetic acid in Acetonitrile) at the same flow rate for five minutes. This was followed by a 1 min ramp up to 100% Solvent B, where it was re-equilibrated with Solvent A (1% Acetic Acid) to 5% solvent B over the course of 1 min and held there for 2 min. The column temperature was a constant 29 °C. The Mass Spectrometer was operated in positive ion mode. Using a Heated ESI, the Source Voltage was set to 4.1 kV, and the capillary temperature was 350 °C. For non-library degradation studies, 2.5 μL of 10 mM peptide stock solution for each peptide was added. For all degradation studies using soluble peptides, a non-proteolytically degradable NH_2_-βFβAβAβAβAβAβA-amide peptide, where βF is β-phenylalanine and βA is β-alanine, was added at a 50 μM concentration as used as an internal standard. Peptides degradation ratios were measured by LCMS and analyzed by Xcalibur. The relative concentrations of peptide were calculated by the ratio of peak area of peptide and peak area of βFβA_6_.

### Hydrogel Fabrication

8-arm 20 kDa poly(ethylene glycol) (PEG) functionalized with dibenzocyclooctyne (DBCO) was purchased from BiopharmPEG and used as received. PEG batches which did not quickly go into solution were further purified. This was done by dissolving the 200 mg of PEG-DBCO in 20 mLs of isopropanol and using a 10 kDa centrifugal filter to remove impurities by centrifuging the PEG-DBCO at 4,500 RPM until less than 1 mL was remaining. This was repeated two more times, and then PEG-DBCO left in the upper part of the filter was dried under vacuum and then used.

For all hydrogel studies, total volume of hydrogel was 20 μL, made with 3.5% (wt/wt) PEG, 40% of the arms were functionalized crosslinking peptides and 10% were functionalized with cyclic-RGD with a final cell concentration of 20 million cells per mL. Briefly, the PEG was dissolved at 140 mg/mL in PBS, the crosslinking peptides were dissolved at 22.4 mM in PBS and the cyclic RGD was dissolved at 5.6 mM in PBS. hMSCs were harvested from T-75 flasks. After centrifugation and pouring the supernatant liquid, the cells were resuspended at 80,000,000 cells/mL or 4,000,000 cells/mL. The cyclic RGD and PEG PBS solutions were mixed at the ratio of 1:1. Add 10 μL of the PEG - cyclic RGD mixer in the wells and incubate for 10 mins. The crosslinking peptides and cell solutions were mixed at a ratio of 1:1. 10 μL of the peptide - cell mixture was pipetted into the PEG - cyclic RGD mixer previously dropped in the wells. The solutions were pipetted until well mixed and after 15 minutes in the incubator 1 mL of media was added into each well and store in the incubator. For confocal microscope imaging gels, place 12 mm round slides in each well of 24 wells plates. For cell viability test, sylgard layers were prepared in 24 well plates to prevent unexpected cell growth on the well bottom wall. Generally, Sylgard kit Part A/Part B were mixed in a 10:1 ratio by volume and the mixture was added to the 24 well plates such that the bottom of each well was completely covered by Sylgard. The plates were placed in a biosafety hood overnight with UV light on. For all studies three technical replicates were done per condition, and the studies were repeated three times.

### Phalloidin/DAPI staining

The hydrogels were washed with PBS three times and incubated n 4% formaldehyde in PBS for 20 minutes. The hydrogels were again washed with PBS three times, followed by an incubation in 0.25% triton-x100 in PBS solution for 20 mins. The hydrogels were washed with PBS three times, followed by an 20 minute incubation of in 1% BSA in PBS. The gels were washed 3x in PBS and 700 μL of Phalloidin/DAPI PBS staining solution (0.1% BSA, 1 μg/mL DAPI and 1 μg/ mL phalloidin-iFluor 555) was added to each well and it was incubated for 1 hour in the dark at room temperature. The hydrogels were washed 3x and 1 mL PBS was added into in each well and they were stored in a 4 °C fridge until they were ready for imaging.

### Quantification of Cell Spreading

Hydrogels were imaged by confocal microscope (LSM 800, AxioObserver, EC Olan-Neofluar 10x/0.3 objective). The 12 mm round slides with hydrogels were taken out of 24 well plates and flipped gel side down on round bottom slides. Confocal microscopy was carried out in the Z-stack mode. Phalloidin was excited at 488 nm and recorded between 493 and 630 nm. DAPI was excited at 405 nm and recorded between 410 and 501 nm. Cells in each hydrogel were measured with ImageJ software according to the particle analysis results and then averaged to obtain the average cell area in the hydrogels.

### Viability Testing

#### alamarBlue metabolic activity assay

On Days 1, 3, and 7, the media from the wells was removed add 500 μL of a 10x diluted Alamar Blue stock solution in media was added to each well with hydrogels. Three wells without any cells were used for background subtraction. The gels were incubated for four hours, at which time the absorbance was measured with a spectrophotometer at wavelength of 579 nm (600 nm as reference). The cell viability percentage was assessed according to the manufacturer’s protocol.

#### DNA quantification assay

Hydrogels seeded in Sylgard coated plates were homogenized in 300 mL Papain digestion solution (125 μg/mL Papain, 2 mM L-cysteine and 0.333 M EDTA) over night at 60 °C to dissolve the samples and completely release their Double Stranded DNA (dsDNA) into solution. dsDNA was quantified using the Quantiflour® dsDNA System (E2670, Promega) according to the manufacturer’s protocol. a plate reader (SpectraMax® iD3, Molecular Devices) was used to measure the fluorescence (504nm Ex /531nm Em).

### Statistical analysis

Statistical analysis was done using a Student’s t-test for comparisons with two conditions, and ANOVAs with a Tukey post-hoc test for comparisons with more than two conditions. All statistical analysis was done in R, and samples were considered statistically significant if they had a p-value of less than 0.05.

Description of the TAILS procedure and characterization of the peptides and peptide libraries can be found in the supporting information.

