## Supplemental Information for "Optimizing Peptide Crosslinks for Cell-Responsive Hydrogels"

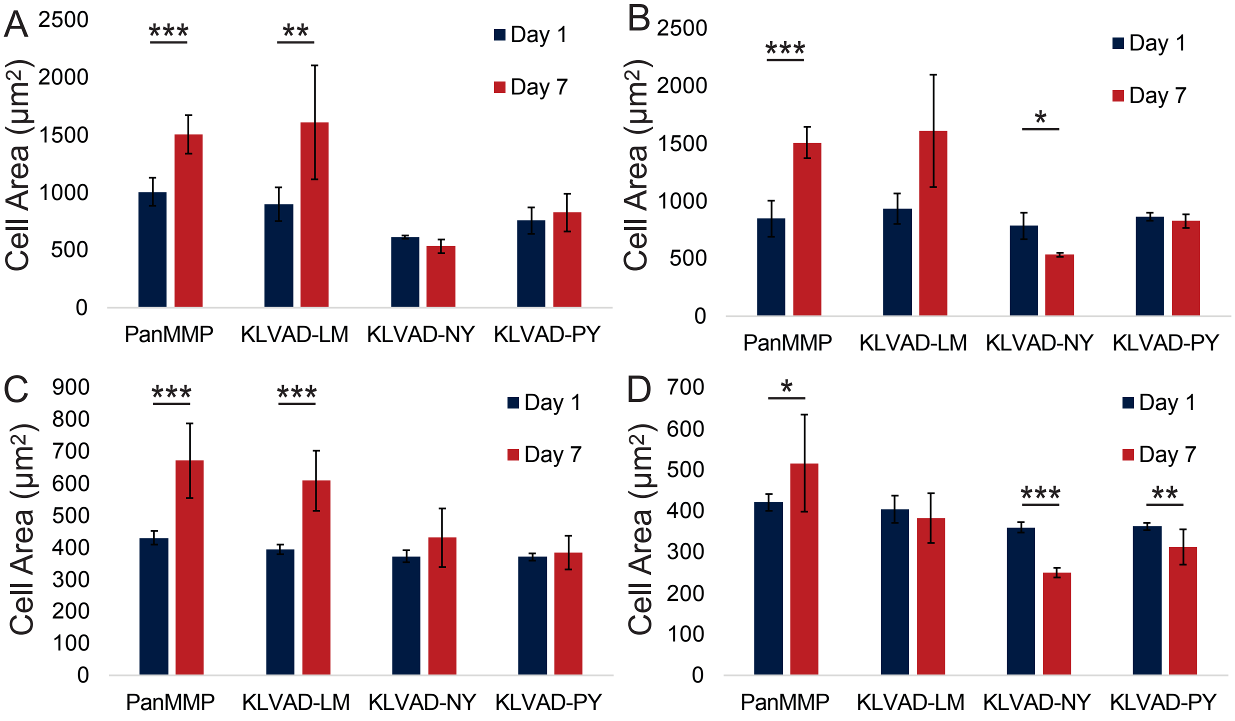


**Fig. S1.** Cells were cultured in PEG hydrogels at 1 million cells/mL with and without an MMP-14 inhibitor and their area was measured using confocal microscopy. (**A**) hMSCs without the inhibitor, (**B**) hMSCs with the inhibitor (**C**) hUVECs without the inhibitor and (**D**) hUVECs with the inhibitor. Notably, the ability of hUVECs to spread in KLVAD-LM gels after 7 days was diminished when the MMP-14 inhibitor was added. * indicates p < 0.05, ** indicates p < 0.01, and *** indicates that p < 0.001 by Tukey's host-hoc test.

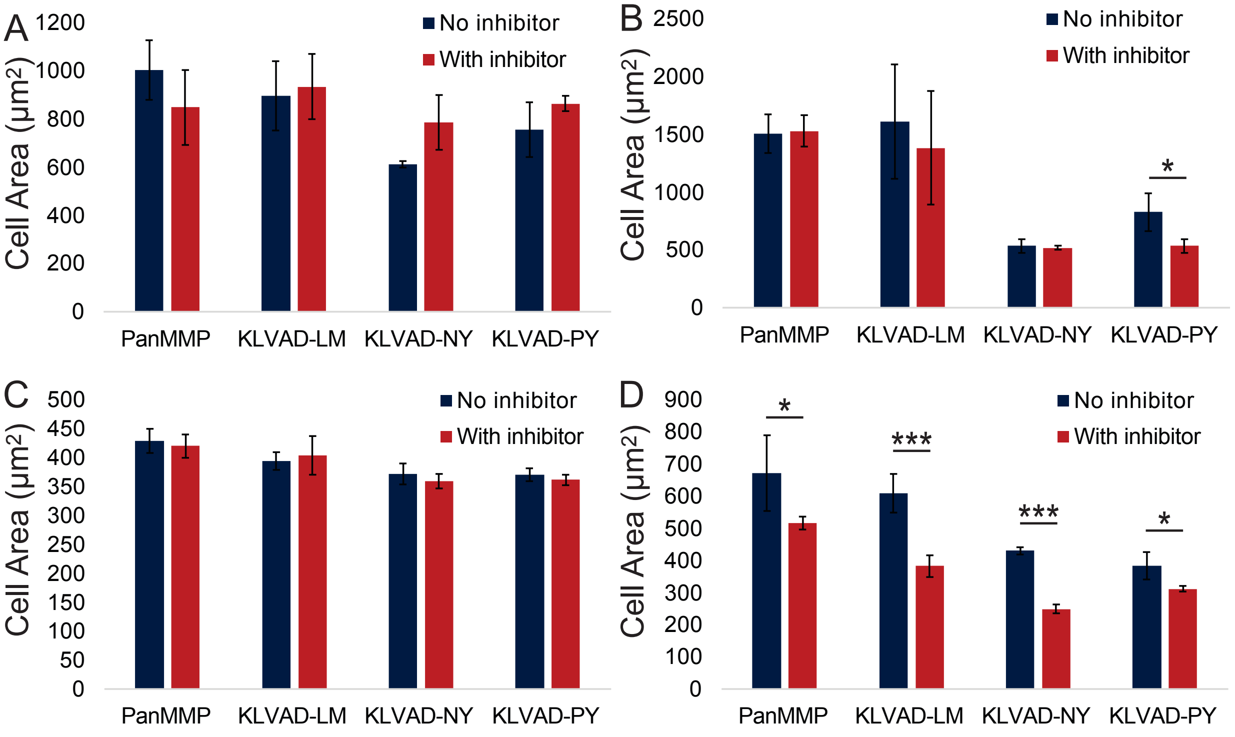

**Fig. S2.** Cells were cultured in PEG hydrogels 1 million cells/mL with and without an MMP-14 inhibitor and their area was measured using confocal microscopy. (**A**) hMSCs at Day 1, (**B**) hMSCs at Day 7, (**C**) hUVECs at Day 1, and (**D**) hUVECs at Day 7. * indicates $p < 0.05$, ** indicates $p < 0.01$, and *** indicates that $p < 0.001$ by Tukey's host-hoc test.

### TAILS Proteomics Procedure

Cells were grown in their respective media until they reached 70% confluence, and four T-75 flasks were used per cell type. The media was removed and the cells were washed 3x to remove serum proteins and dead cells. Serum-free media was added for 30 minutes, and then replaced with fresh serum-free media. Cells were grown for 24 hours at which time the media was collected, treated with the HALT protease inhibitor cocktail, and centrifuged at 2,200g to remove dead cells. The centrifuged media was filtered using a 0.2 µm syringe filter and 15% (v/v) of 100% trichloroacetic acid (TCA) was added to precipitate the proteins. This was put on ice for 3-4 hours, and then centrifuged at 9000g for 15min at 4°C and the supernatant was decanted. The pellet was washed with freezer-cold (-20 °C) acetone and it was centrifuged again at 9000g for 15min at 4 °C. Three acetone washes per performed and then the pellet was resuspended at 8M guanidinium hydrochloride was added to make a protein concentration of 2 mg/mL, which was determined using a BCA protein quantification assay.

*Disulfide reduction an alkylation*

The pellet was sonicated until it went into solution, and then this was mixed 1:1 with 200 mM HEPES at pH 7 to make a final protein concentration of ~1-2 mg/mL. A fresh solution of 1M dithiothreitol (DTT) was made and added to make a final DTT concentration in the sample of 5 mM. This was incubated with the proteins for 1 hour at 65 °C to reduce the disulfide bonds on the proteins. The samples were cooled to room temperature, and then a solution of 0.1M iodoacetamide was added to a final concentration of 15 mM to alkylate the cysteines on the proteins. The sample was vortexed and incubated in the dark for 30 minutes at room temperature. 20 µL of 1M DTT was added per mL and vortexed.

The proteins were then isolated using a water:methanol:chloroform protocol. 3 mLs of ice-cold water were added per mL of protein solution. This was then mixed with 4 mLs of ice-cold methanol per mL of protein solution. Then 1 mL of ice-cold chloroform was added per mL of protein solution, and it was vortexed to mix the solution. This was centrifuged at 3,000 x g at 4 °C for 7 min and the white protein interphase was isolated using a spatula and transferred to a 1.5 mL microcentrifuge tube. This pellet was washed 3x with 1 mL of methanol.

*Dimethylation of primary amines*

The pellet was resuspended/dissolved in 100 µL of 0.1 M NaOH was added to the pellet, followed by 50 µL of 1M HEPES, pH 7.5, and 100 µL of water. The pellet was sonicated until the protein went into solution. This was mixed with 250 µL of 6 M guanidinium hydrochloride and the pH was adjusted to 7.5 using either NaOH or HCl. A fresh 2M working stock of formaldehyde and a fresh 1M working stock of $NaBH_3CN$ (typically 6.3 mg/100 µL of water) were made. 10 µL of 2M formaldehyde was added to the 500 µL protein solution make a 40 mM final concentration, immediately followed by 10 µL of of the 1M $NaBH_3CN$. The pH was adjusted to 6-7 and this was incubated overnight at 37°C to dimethylate all primary amines on the proteins, including both lysines and N-terminal amines. The following morning an additional 20 mM of fresh formaldehyde and 10 mM of fresh $NaBH_3CN$ was added and incubated at 37°C for one hour to ensure complete dimethylation and the reaction was quenched by adding 1 M Tris, pH 6.8 to a 100 mM final concentration. The samples were precipitated using the water:methanol:chloroform protocol to precipitate the proteins and the pellet was washed with methanol 3x.

*Trypsin digestion of proteins*
The pellet was resuspended in ~100 µL of 6 M GuCl and then diluted ten-fold with 100 mM HEPES, pH 8.0 for trypsin digestion. Proteomics-grade trypsin was added to the solution at a 1:100 trypsin:sample ratio and mixed with pipetting and vortexing. The sample was incubated overnight at 37°C.

*Negative selection of dimethylated peptides using HPG-ALD polymer*
The HPG-ALD polymer was used to remove tryptic peptides. N-terminal amines present in the sample during the dimethylation step, such as those which were generated from proteolytic cleavage of the protein, were dimethylated and will not be reactive towards the aldehyde groups in the high molecular weight HPG-ALD polymer. N-terminal amines generated in the trypsinization step will react with the aldehyde moieties on the high molecular weight HPG-ALD polymers and can be separated from the peptides with dimethylated N-termini. The frozen HPG-ALD polymer solution was thawed at room temperature and added at a polymer:peptide ratio of 5:1 and then immediately after freshly prepared $NaBH_3CN$ was added to a 20 mM final concentration and the pH was adjusted to 6-7 using either NaOH or HCl. The samples were then incubated overnight at 37°C.

*N-terminome sample recovery*
1M Tris buffer pH 6.8 was added to a 100 mM final concentration. The pH was checked to be between 6-7 and it was incubated at 37°C to quench the aldehydes on the HPG-ALD polymer. An 10-kDa MWCO 0.5 mL Amicon centrifugal filter was pre-washed with 400 µL 100 mM NaOH, centrifuged at 12,000 rcf for 5 min and the flow-through was discarded. A second wash with deionized water was done at 12,000 rcf for 5 min and the flow-through was again discarded. 300 µL of the HPG-ALD/peptide mixture was laded into the centrifugal filter and it was centrifuged at 12,000 rcf for 10 minutes and the flow through which contains the desired dimethylated peptides was collected and transferred to a fresh 15 mL tube. The remaining HPG-ALD/peptide mixture was added to the centrifugal filter and was spun at 12,000 rcf for 10 minutes and the flow through was collected. The remaining HPG-ALD in the filter was washed with 50:50 methanol:water to solubilize hydrophobic peptides and the centrifugation step was repeated and the flow through was collected into the 15 mL tube. The combined flow throughs were lyophilized, and then re-dissolved in 1-2 mL of water with 5% acetonitrile and 0.1% trifluoroacetic acid. This was then injected onto a prep-HPLC for desalting. Briefly, after the sample was injected the gradient was kept at 95% water and 5% acetonitrile with 0.1% TFA for five minutes to allow the salts to elute. The gradient was then ramped up to 100% acetonitrile (with 0.1% TFA) over two minutes and held there for three minutes, at which point it was ramped back down to 95% water and 5% acetonitrile with 0.1% TFA. Fractions which eluted peptides after the beginning of the ramp were collected, combined, and lyophilized.

*Cation exchange chromatography*
Dimethylated peptides were then fractionated using strong cation exchange (SCX) chromatography using a 100 mm × 4.6 mm, 5 µm, 300 Å PolySULFOETHYL SCX column from PolyLC. One mg was loaded onto the column per injection, and a gradient from 100% Buffer A (30% acetonitrile, 5 mM KH2PO4, pH 2.7) to 100% Buffer B (30% acetonitrile, 5 mM KH2PO4, 0.5 M NaCl, pH 2.7) over 30 minutes and was run at a flow rate of 1 mL per minute. The collected fractions were combined into approximately 10 fractions and lyophilized. The samples were then dissolved in 1 mL of water, and desalted using C18 Sep-Pak Vac 6cc cartridges. Briefly the columns were conditioned with 6 ml of isopropanol and equilibrated with 6 ml of 0.1% formic acid. Once the 0.1% formic acid had been flushed through, the samples were loaded and it was washed with water containing 4.25% acetonitrile and 0.1% formic acid to wash the salts through. The peptides were then eluted with 50:50 acetonitrile:water and 0.1% formic acid. The samples were then lyophilized and ready for liquid chromatography mass spectrometry.

*Proteomics Using Liquid Chromatography Mass Spectrometry*

Samples were either sent to the proteomics core facility at the University of Pennsylvania or analyzing in the lab using a Thermo Fisher Vanquish ultra high pressure liquid chromatography (UPLC) and an LTQ-XL mass spectrometer using a HALO Peptide ES-C18 1.0x250mm 2.7 µm UHPLC Column 1.0x250mm (Mac-Mod) at 55°C under micro-flow 50 µL/min conditions.1 The samples were dissolved into 40 µL and 10 µL was injected onto the UPLC column. The mobile phase was ramped between 95% Phase A (water with 1% acetic acid) and 5% Phase B (acetonitrile with 1% acetic acid) over 60 minutes and peptides were identified using a Top 5 MS/MS method.

*Proteomics Analysis*

The data was processed using the trans-proteomics pipeline (TPP) (http://www.tppms.org/). The LTQ RAW data was converted to mzXML format in profile mode (not centroid) using msconvert tools in the TPP. Peptides were identified using both the X!Tandem and Mascot databases and peptide validation was done using the Peptide Prophet. For X!Tandem searches the following parameters were used: fragment mass tolerance 0.8 kDa and fixed modifications including: cysteine carbamidomethylation (+57.021464) and peptide N-terminal and lysine dimethylation (+28.031300). Variable modifications: methionine oxidation (+15.994915), asparagine deamidation (+0.984016) and glutamine deamidation (+0.984016) For Mascot searches the following peptide modifications were used: cysteine carbamidomethylation (+57.021464), peptide N-terminal and lysine dimethylation (+28.031300); variable modifications: methionine oxidation (+15.994915), asparagine deamidation (+0.984016) and glutamine deamidation (+0.984016).

The results from the X!Tandem and Mascot searches were analyzed using the Xinteract, PeptideProphet and XPRESS tools of the TPP. The following PeptideProphet options were used: ‘Use accurate mass binning’, ‘Do not use the NTT model’ and ‘Use decoy hits to pin down the negative distribution’ and to provide the correct identifier for decoy proteins. The output of was an interact pepXML file that included all the peptides and their related information including the peptide sequence, protein, and scoring.

Peptides which had greater than 95% likelihood of an accurate match were selected and peptides with a cysteine were removed. Peptides in this list which came from proteins found either on the cell membrane or extracellularly were then selected. The pepXML file gives the sequence of the peptides found in the fragment identified in LCMS. Since this only the C-terminal side of the site of proteolytic cleavage, the Uniprot database (https://www.uniprot.org/) was used to identify the amino acids on the N-terminal side of the cleavage. Typically peptides of approximately ten amino acids were used, roughly five amino acids on either side of the cleavage site, however this was occasionally shifted to include more hydrophilic/charged amino acids within the sequence to help ensure that the peptides are soluble in cell culture media.

### Peptide Characterization

A

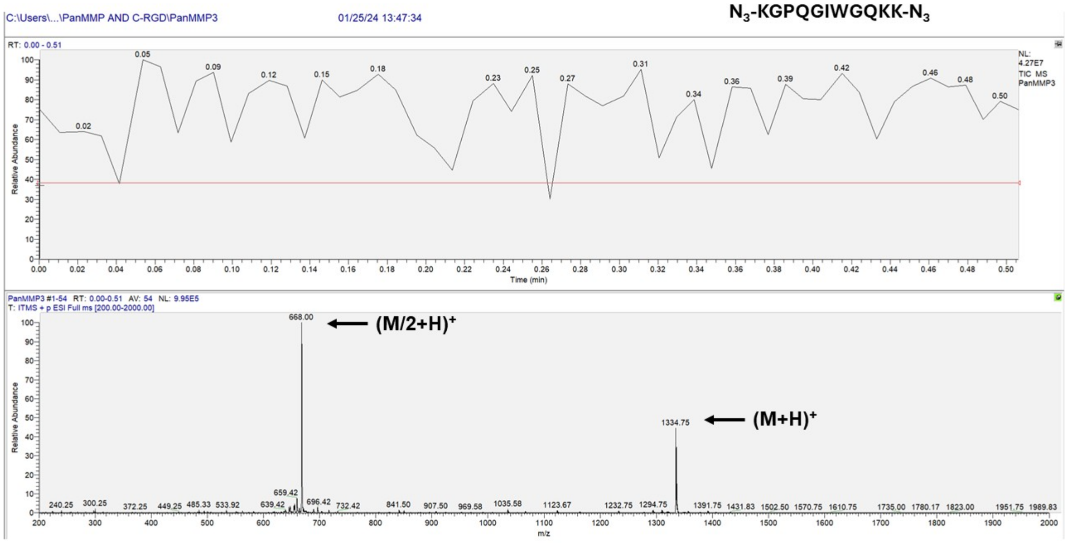


B

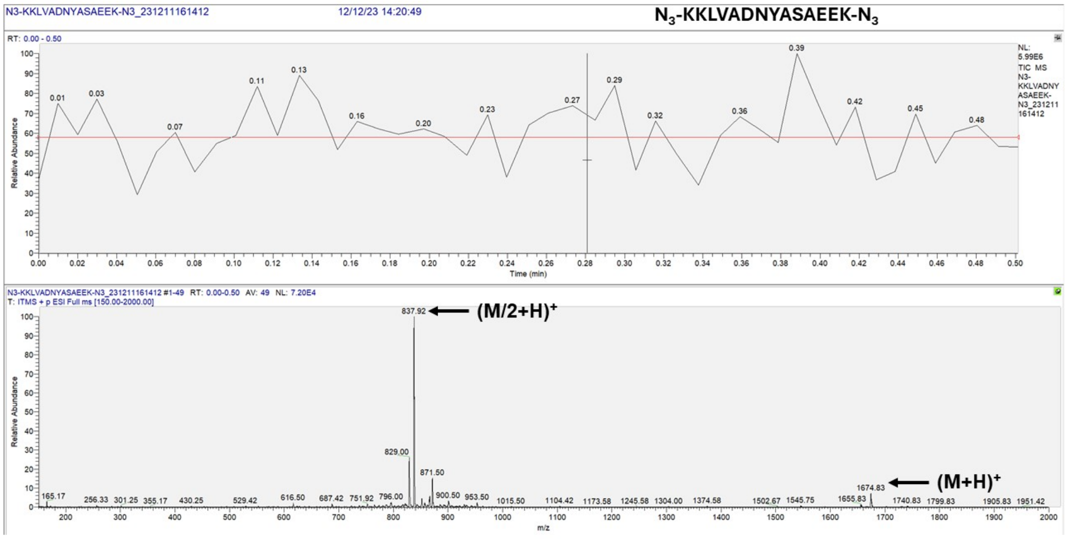

C

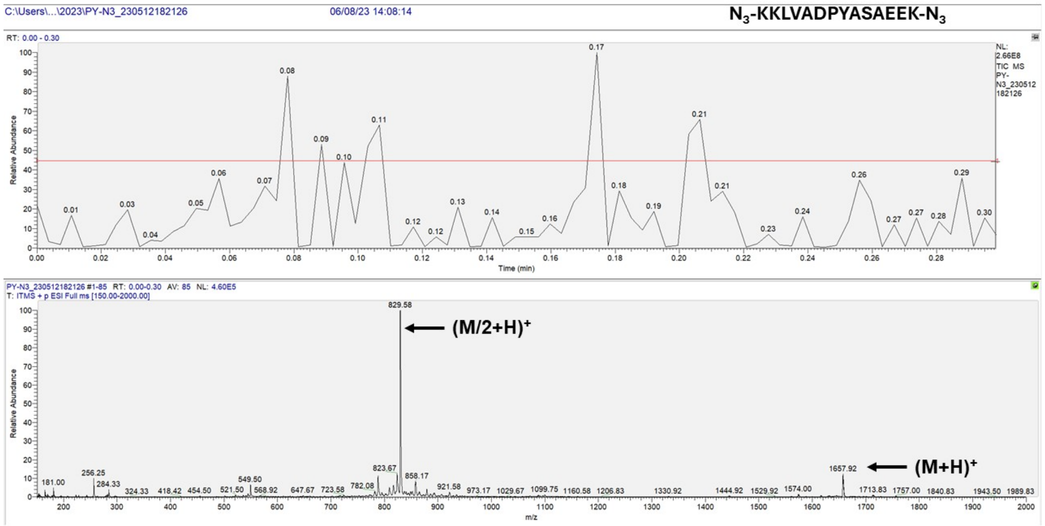
C:\Users\...\2023\PY-N3_230512182126
06/08/23 14:08:14
$N_3$-KKLVADPYASAEEK-$N_3$
RT: 0.00 - 0.30
Relative Abundance
Time (min)
NL: 2.66E8 TIC MS PY-N3_230512182126
PY-N3_230512182126 #1-85 RT: 0.00-0.30 AV: 85 NL: 4.60E5
T: ITMS + p ESI Full ms [150.00-2000.00]
829.58
(M/2+H)+
1657.92
(M+H)+
m/z

D

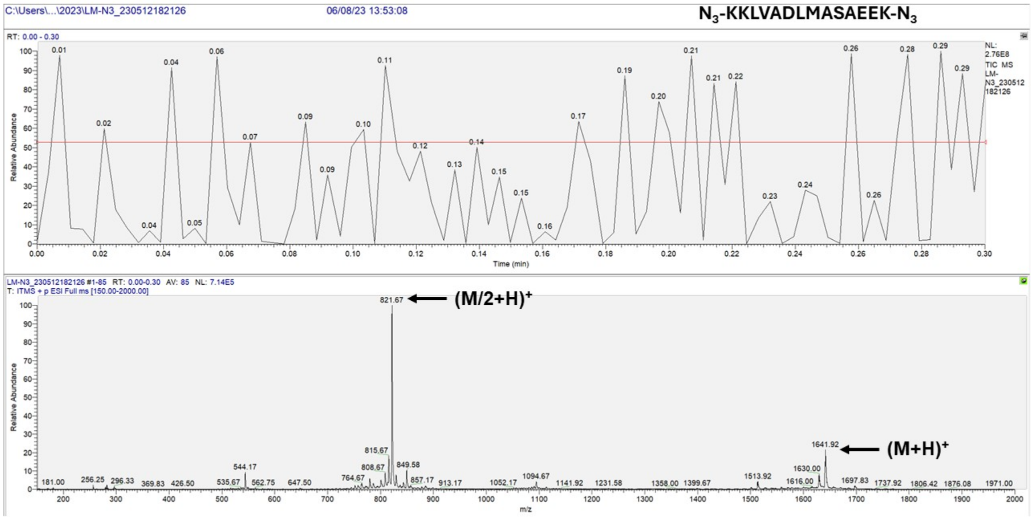
C:\Users\...\2023\LM-N3_230512182126
06/08/23 13:53:08
$N_3$-KKLVADLMASAEEK-$N_3$
RT: 0.00 - 0.30
Relative Abundance
Time (min)
NL: 2.76E8 TIC MS LM-N3_230512182126
LM-N3_230512182126 #1-85 RT: 0.00-0.30 AV: 85 NL: 7.14E5
T: ITMS + p ESI Full ms [150.00-2000.00]
821.67
(M/2+H)+
1641.92
(M+H)+
m/z

E

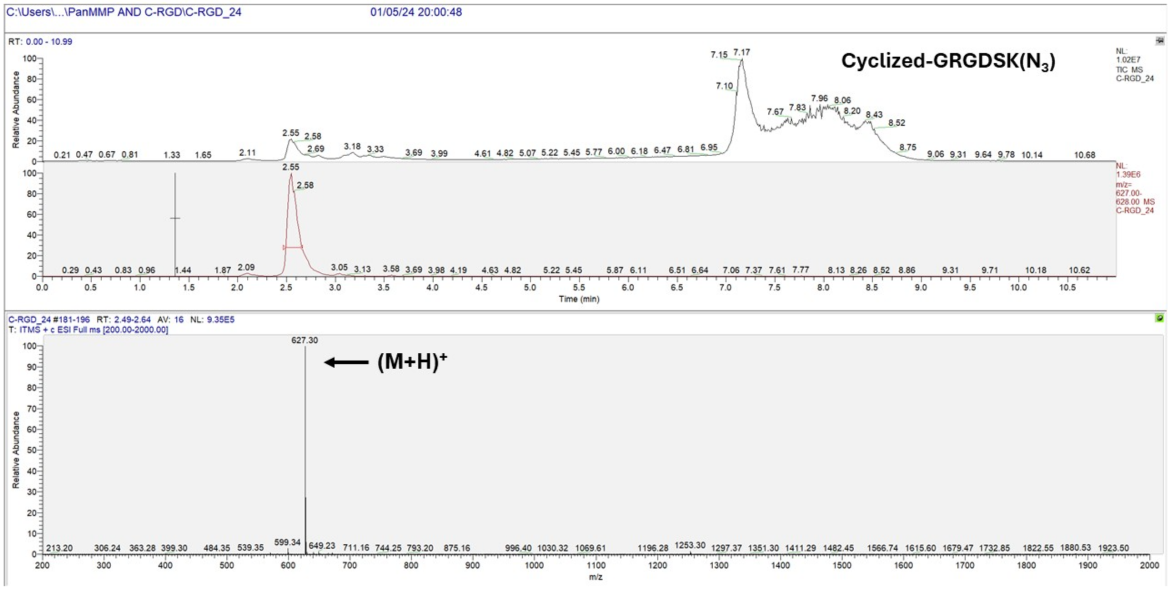


F

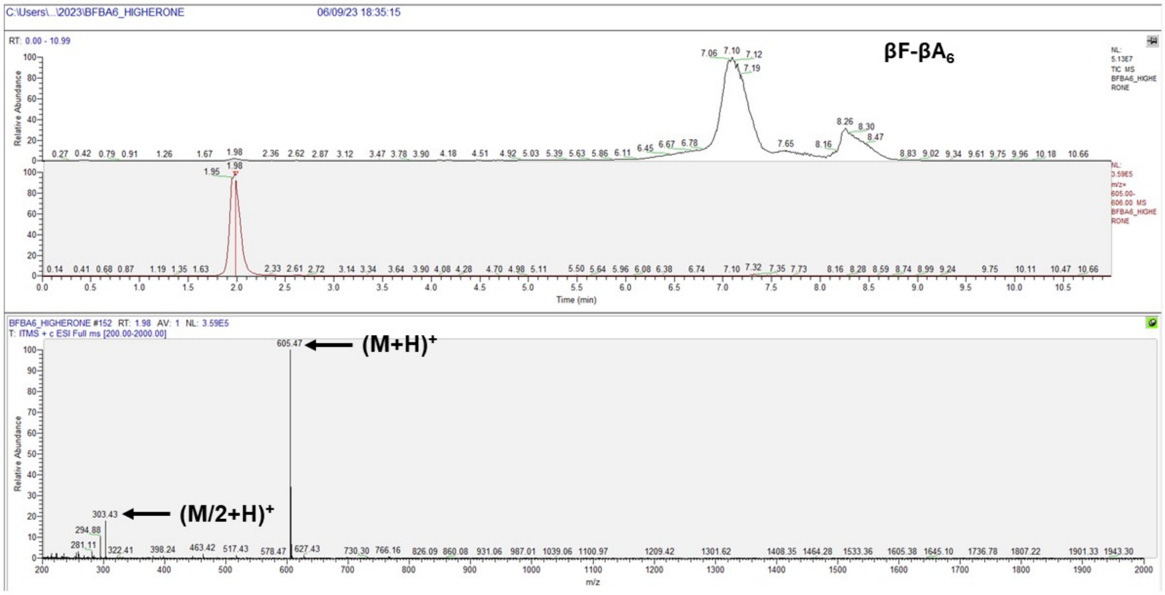


**Fig. S3.** Mass spec characterization of peptides. Crosslinking peptides (**A**) $N_3$-KGPQGIWGQK($N_3$), (**B**) $N_3$-KKLVADNYASAEEK($N_3$) (**C**) $N_3$-KKLVADPYASAEEK($N_3$), (**D**) $N_3$-KKLVADLMASAEEK($N_3$), (**E**) cyclic GRGDSK($N_3$), and (**F**) the βF-β$A_6$ internal standard for degradation studies.

### Characterization of KLVAD-XX-ASAE split-and-pool peptide libraries

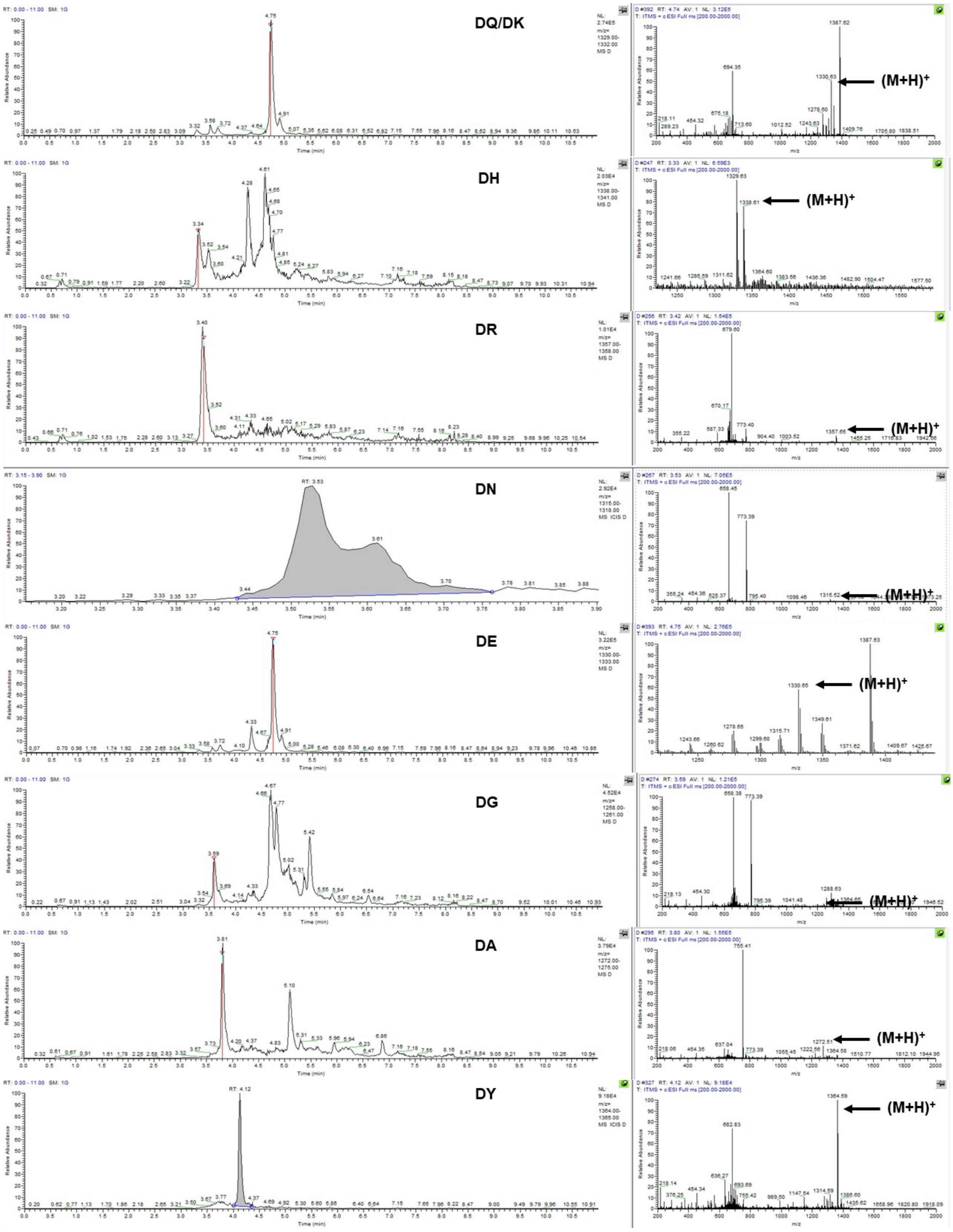

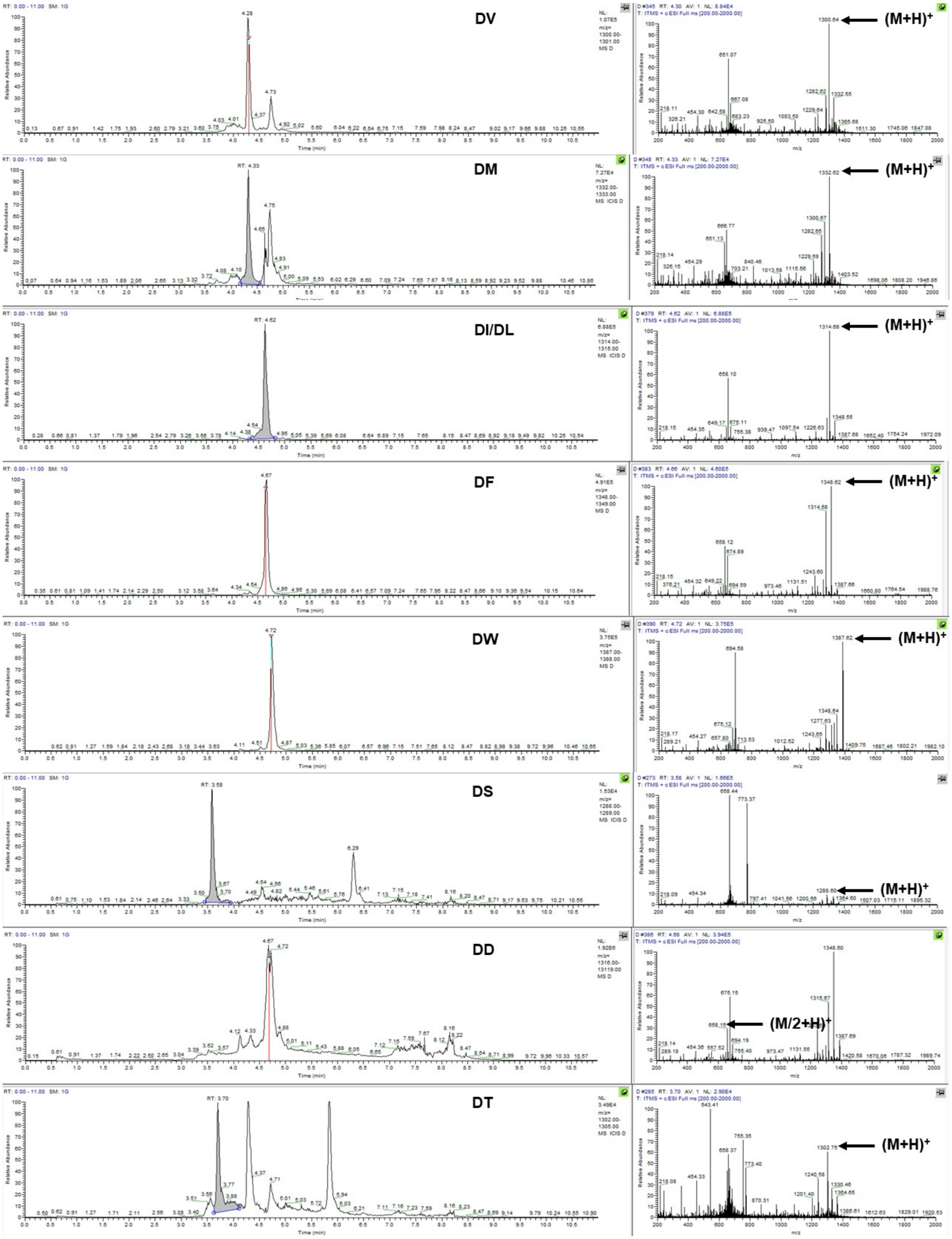
DV
(M+H)+
DM
(M+H)+
DI/DL
(M+H)+
DF
(M+H)+
DW
(M+H)+
DS
(M+H)+
DD
(M/2+H)+
DT
(M+H)+

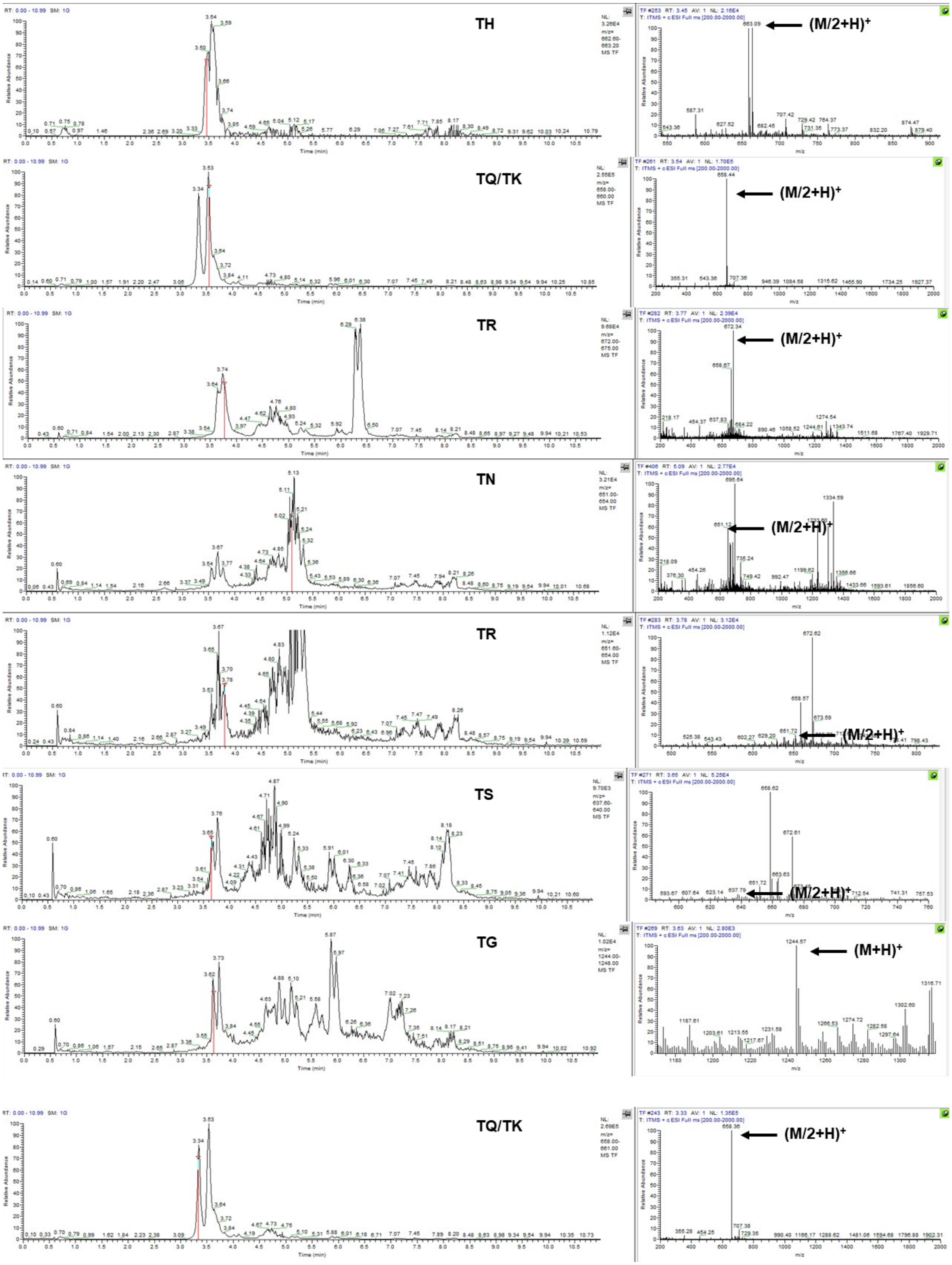
TH
(M/2+H)+
TQ/TK
(M/2+H)+
TR
(M/2+H)+
TN
(M/2+H)+
TR
(M/2+H)+
TS
(M/2+H)+
TG
(M+H)+
TQ/TK
(M/2+H)+

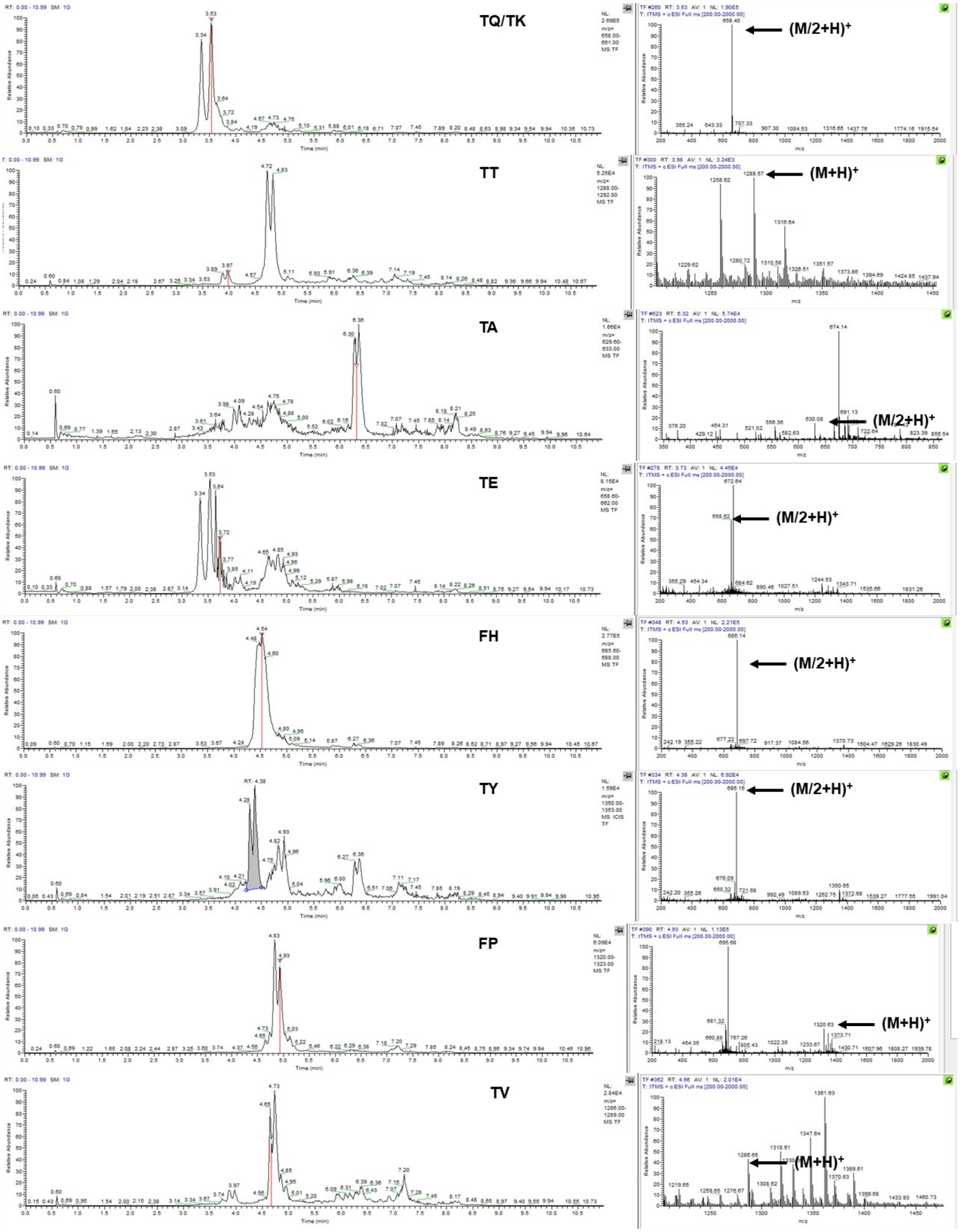
TQ/TK
(M/2+H)+
TT
(M+H)+
TA
(M/2+H)+
TE
(M/2+H)+
FH
(M/2+H)+
TY
(M/2+H)+
FP
(M+H)+
TV
(M+H)+

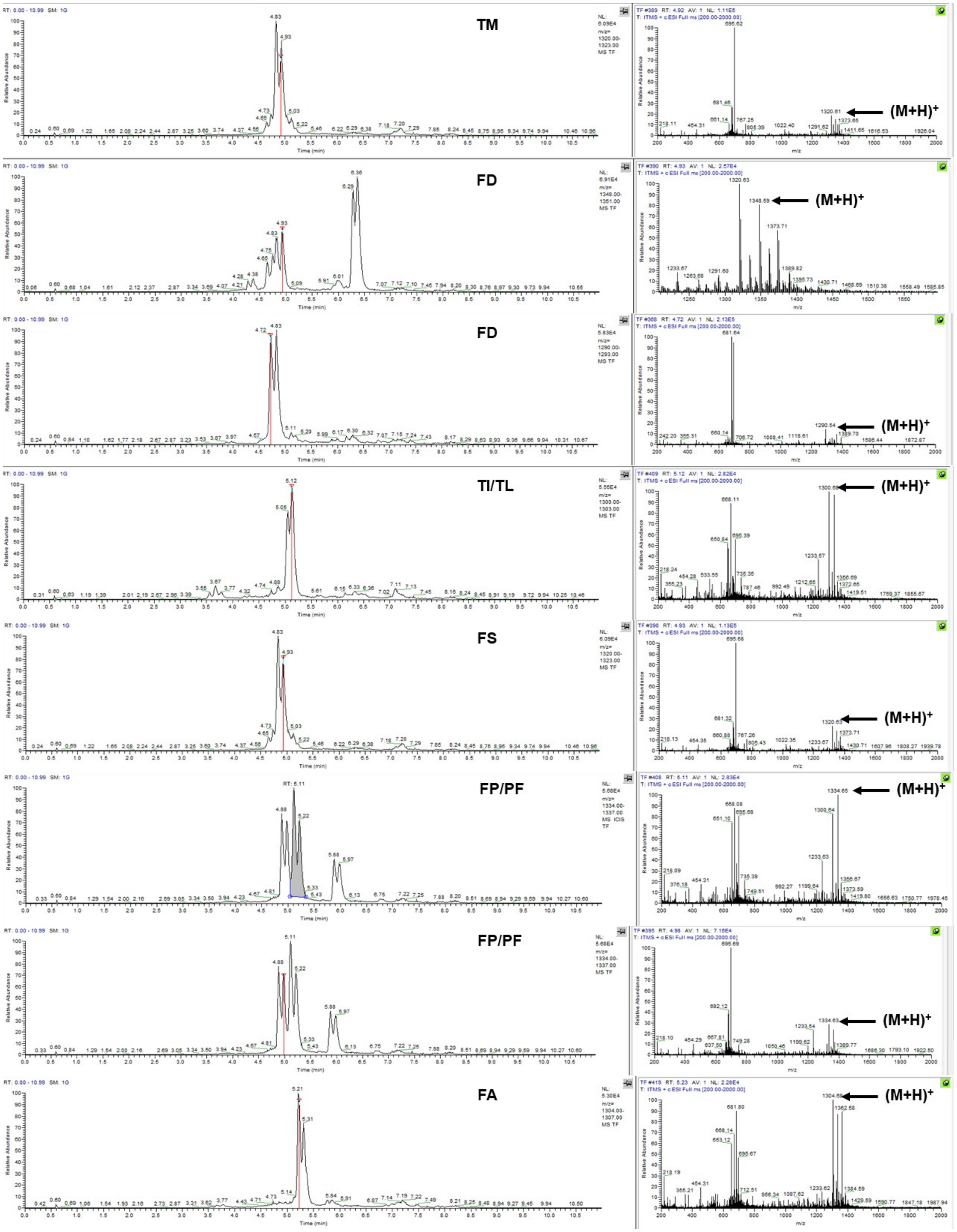
TM
(M+H)+
FD
(M+H)+
FD
(M+H)+
TI/TL
(M+H)+
FS
(M+H)+
FP/PF
(M+H)+
FP/PF
(M+H)+
FA
(M+H)+
Time (min)
Relative Abundance
m/z

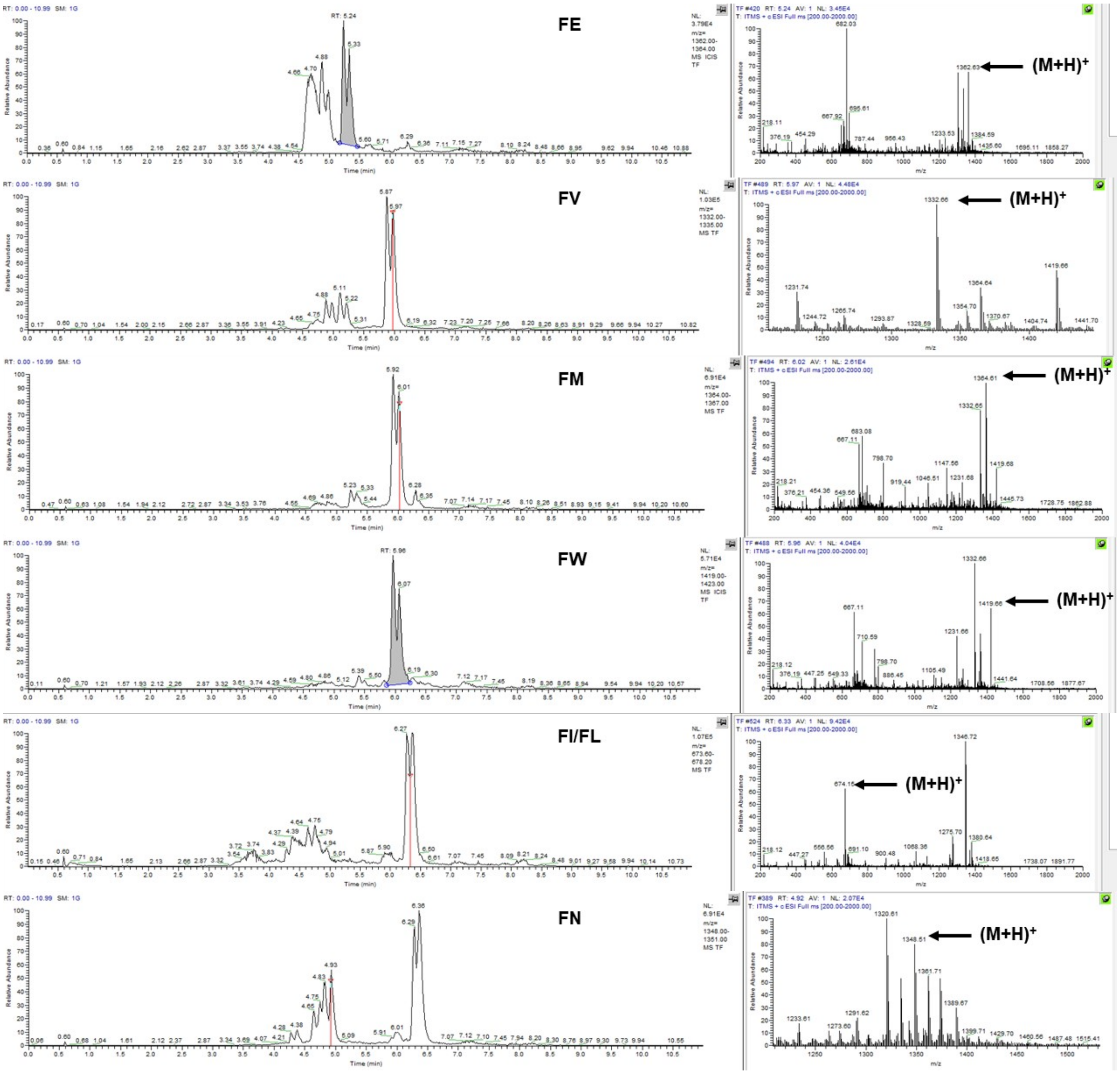
FE
(M+H)+
FV
(M+H)+
FM
(M+H)+
FW
(M+H)+
FI/FL
(M+H)+
FN
(M+H)+
Relative Abundance
Time (min)
m/z

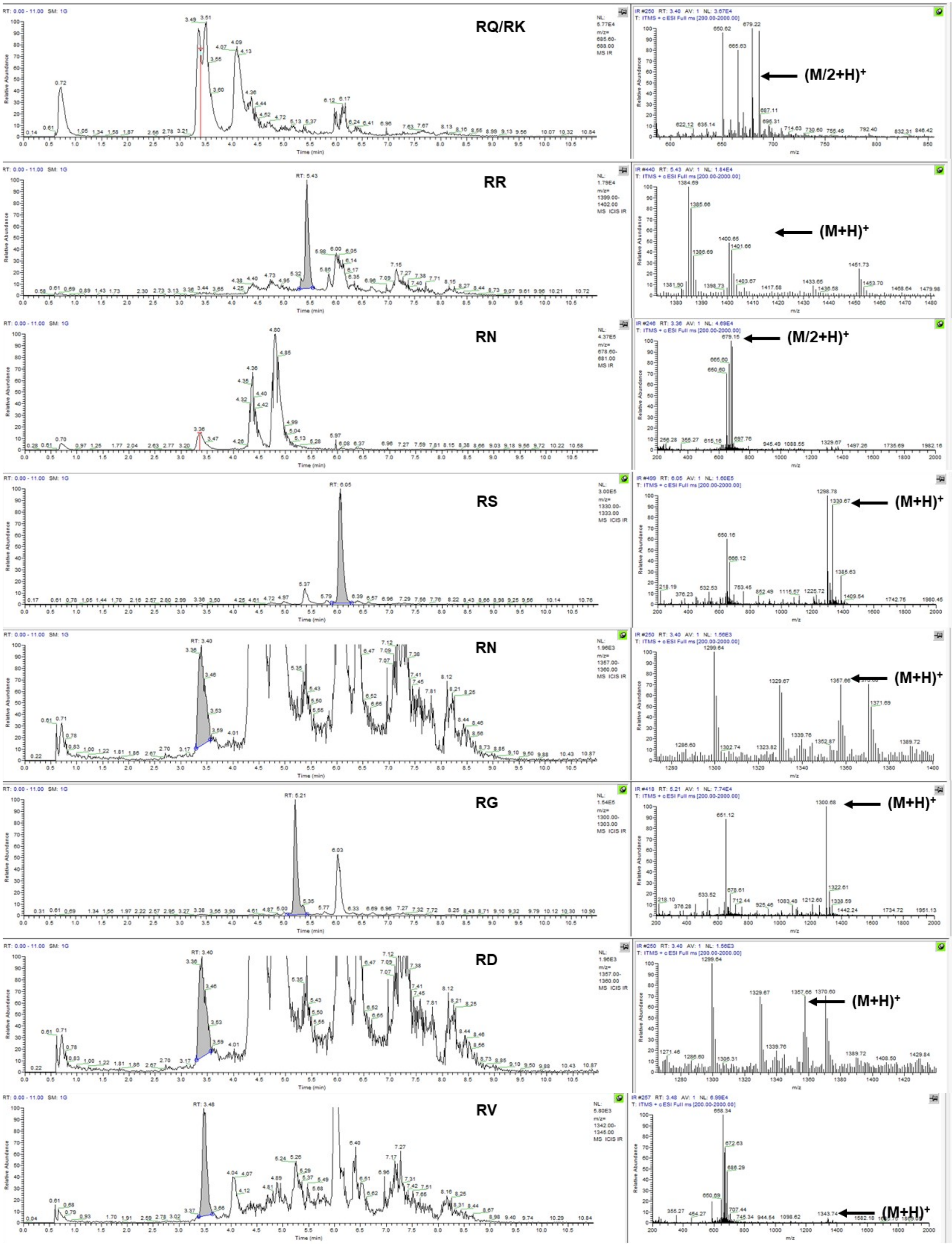
RQ/RK
(M/2+H)+
RR
(M+H)+
RN
(M/2+H)+
RS
(M+H)+
RN
(M+H)+
RG
(M+H)+
RD
(M+H)+
RV
(M+H)+
Relative Abundance
Time (min)
m/z

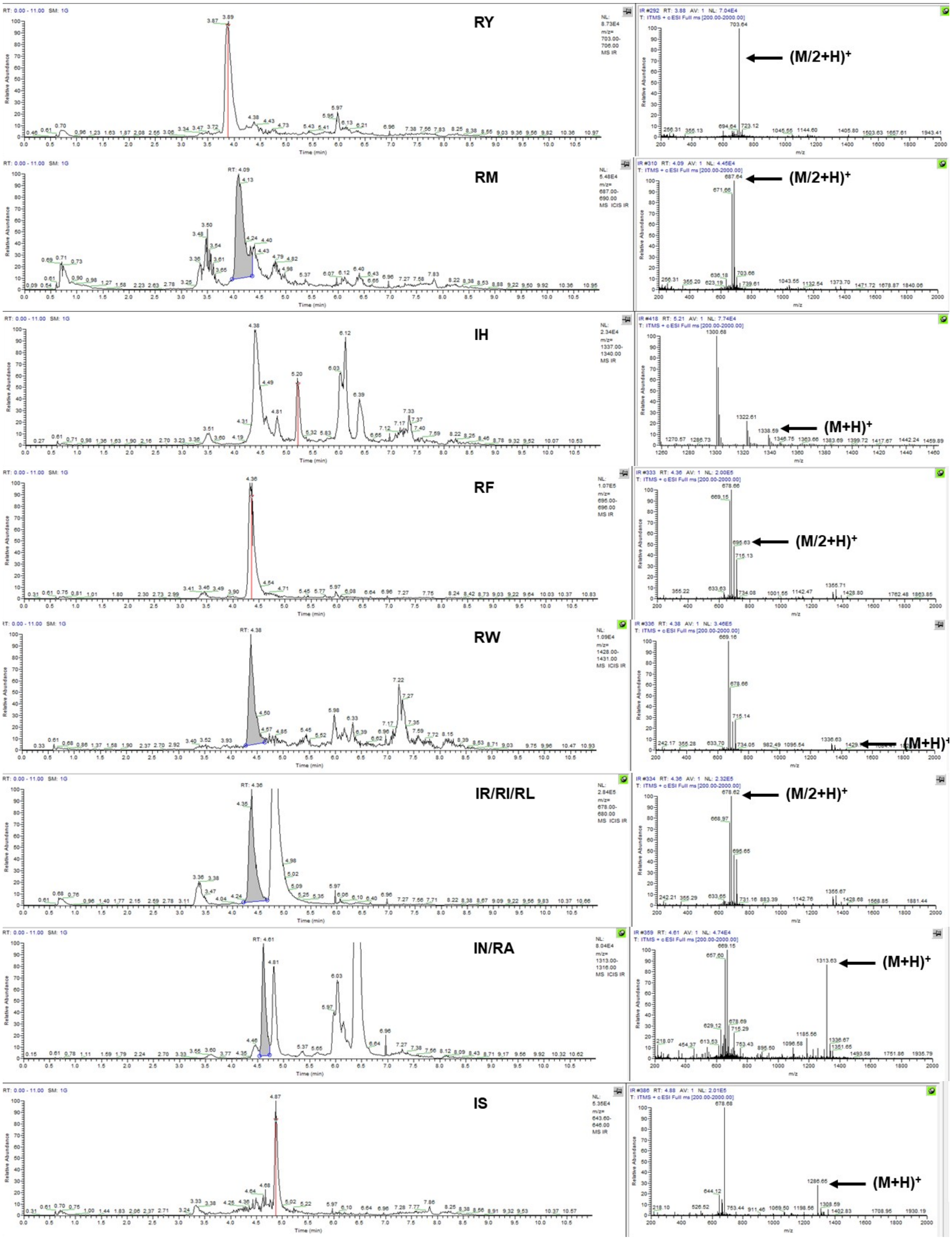
RY
(M/2+H)+
RM
(M/2+H)+
IH
(M+H)+
RF
(M/2+H)+
RW
(M+H)+
IR/RI/RL
(M/2+H)+
IN/RA
(M+H)+
IS
(M+H)+

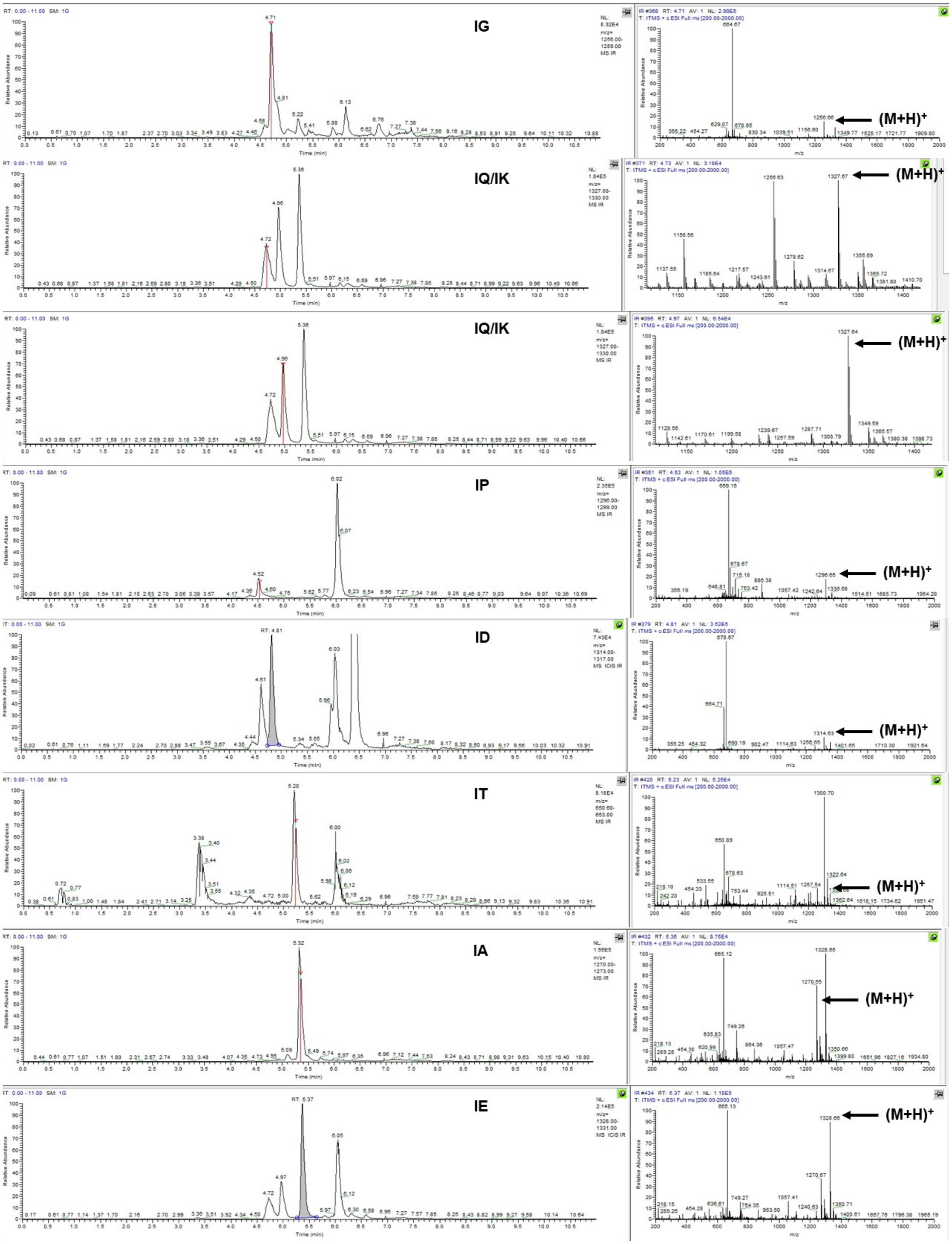
IG
(M+H)+
IQ/IK
(M+H)+
IQ/IK
(M+H)+
IP
(M+H)+
ID
(M+H)+
IT
(M+H)+
IA
(M+H)+
IE
(M+H)+

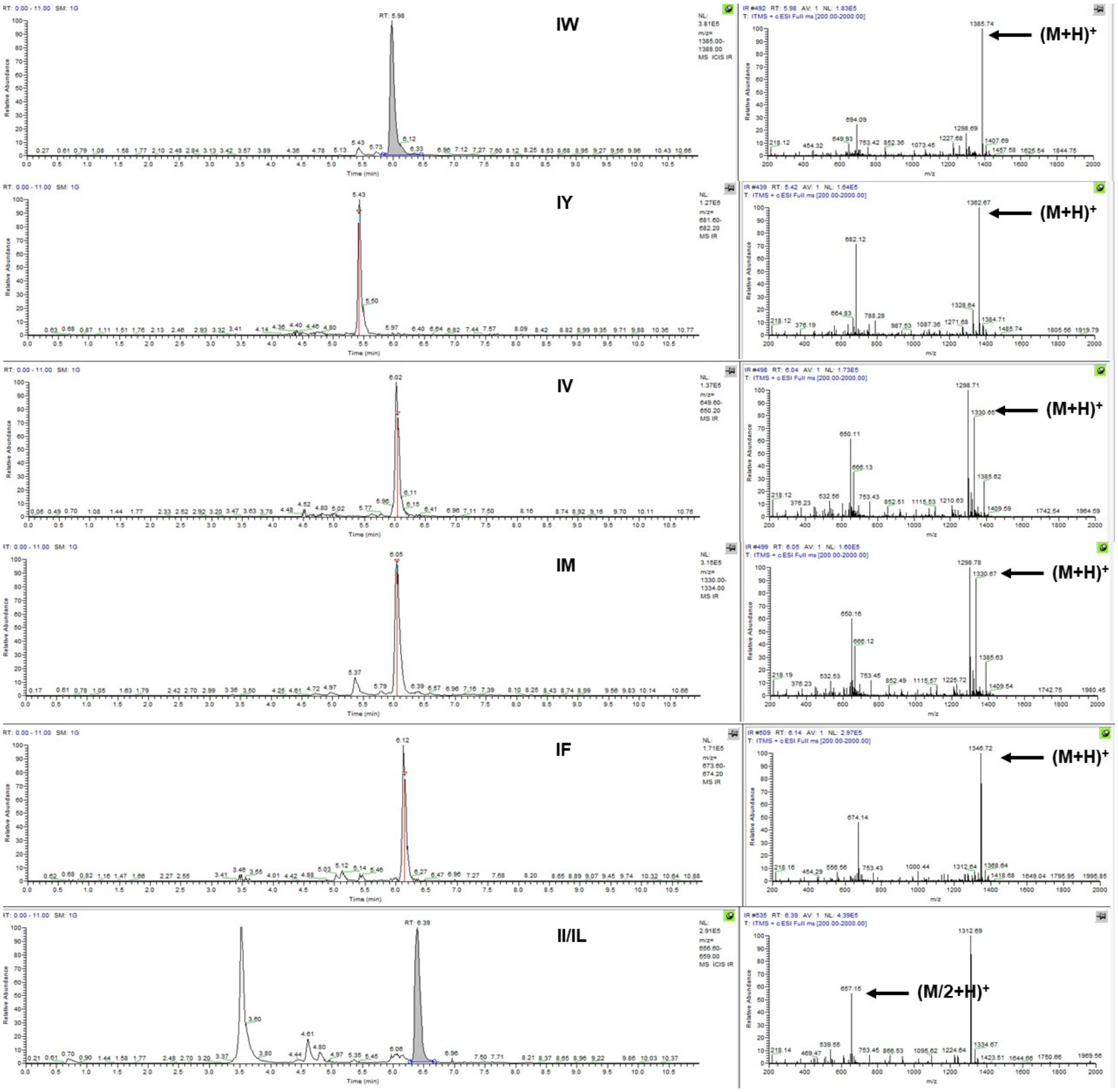
IW
(M+H)+
IY
(M+H)+
IV
(M+H)+
IM
(M+H)+
IF
(M+H)+
II/IL
(M/2+H)+

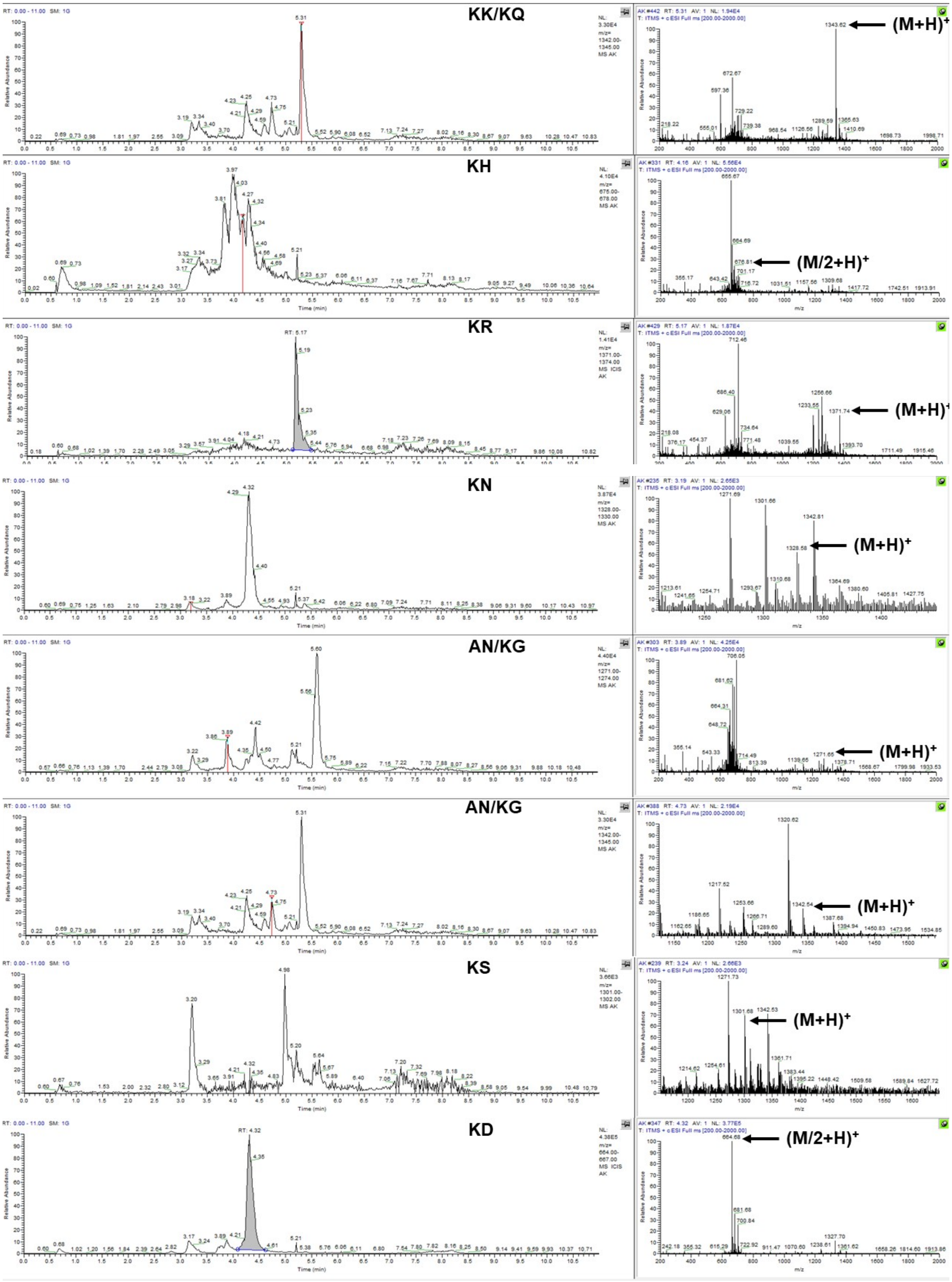
KK/KQ
(M+H)+
KH
(M/2+H)+
KR
(M+H)+
KN
(M+H)+
AN/KG
(M+H)+
AN/KG
(M+H)+
KS
(M+H)+
KD
(M/2+H)+

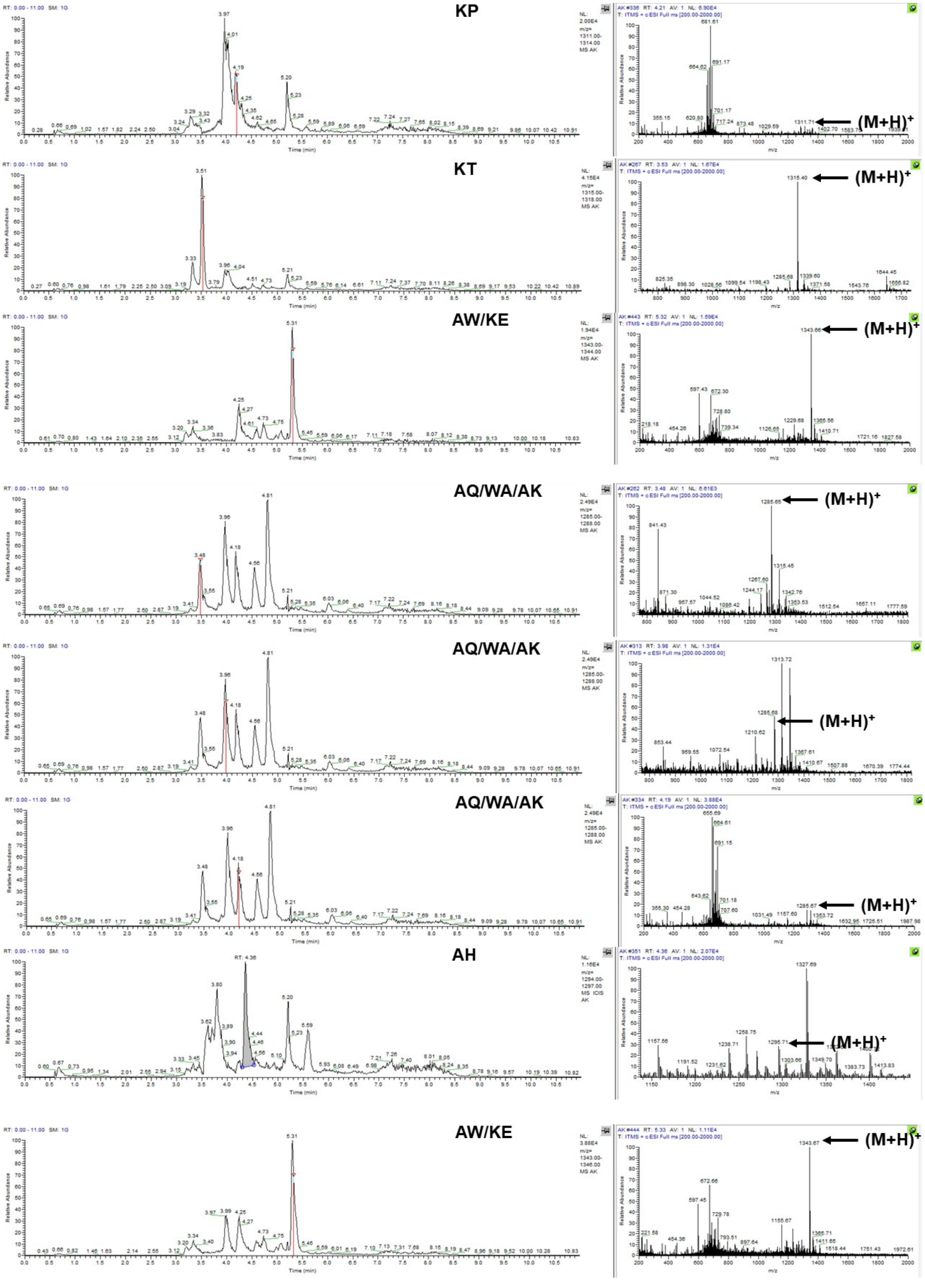
KP
(M+H)+
KT
(M+H)+
AW/KE
(M+H)+
AQ/WA/AK
(M+H)+
AQ/WA/AK
(M+H)+
AQ/WA/AK
(M+H)+
AH
(M+H)+
AW/KE
(M+H)+

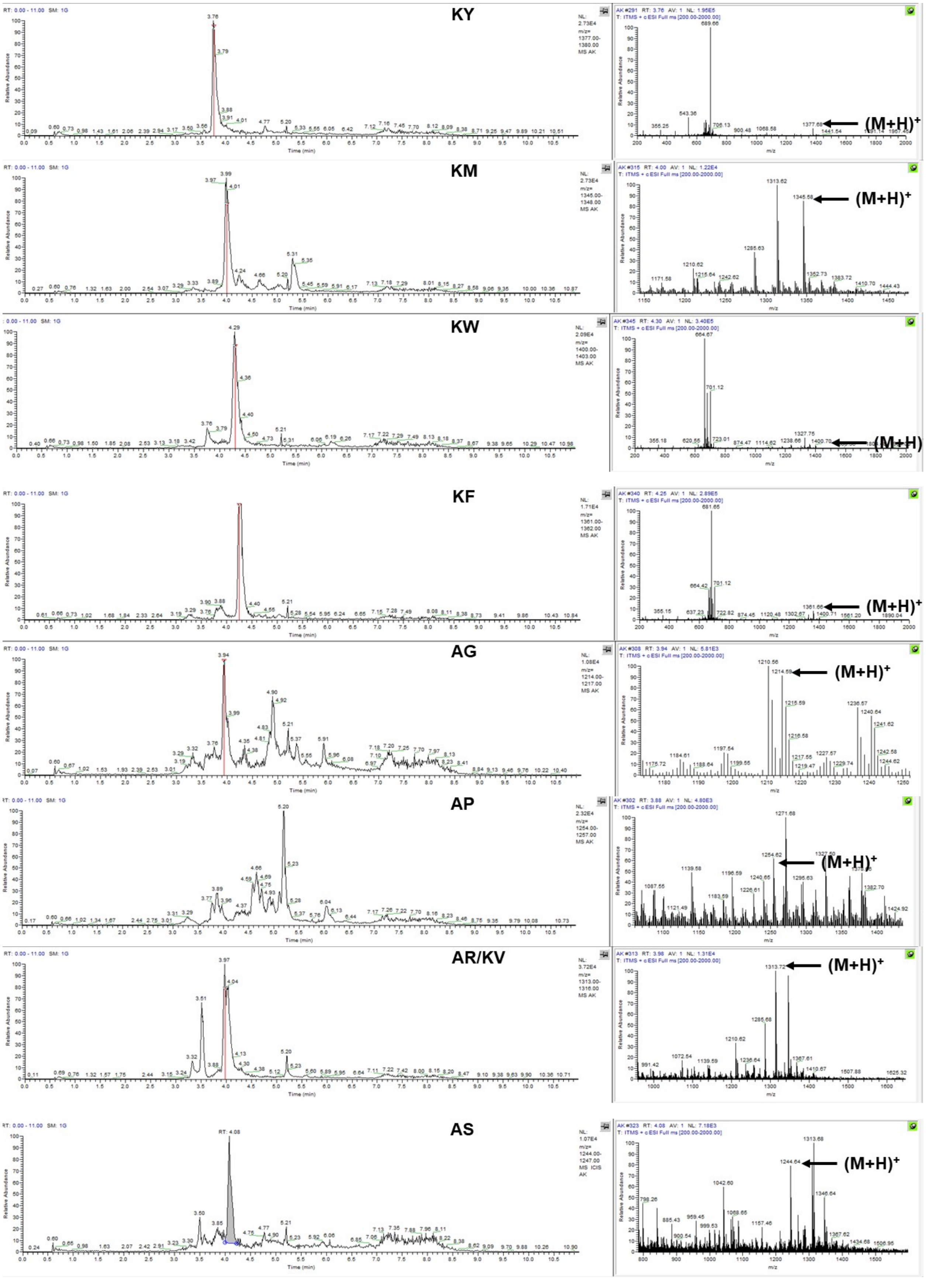
KY
(M+H)+
KM
(M+H)+
KW
(M+H)
KF
(M+H)+
AG
(M+H)+
AP
(M+H)+
AR/KV
(M+H)+
AS
(M+H)+

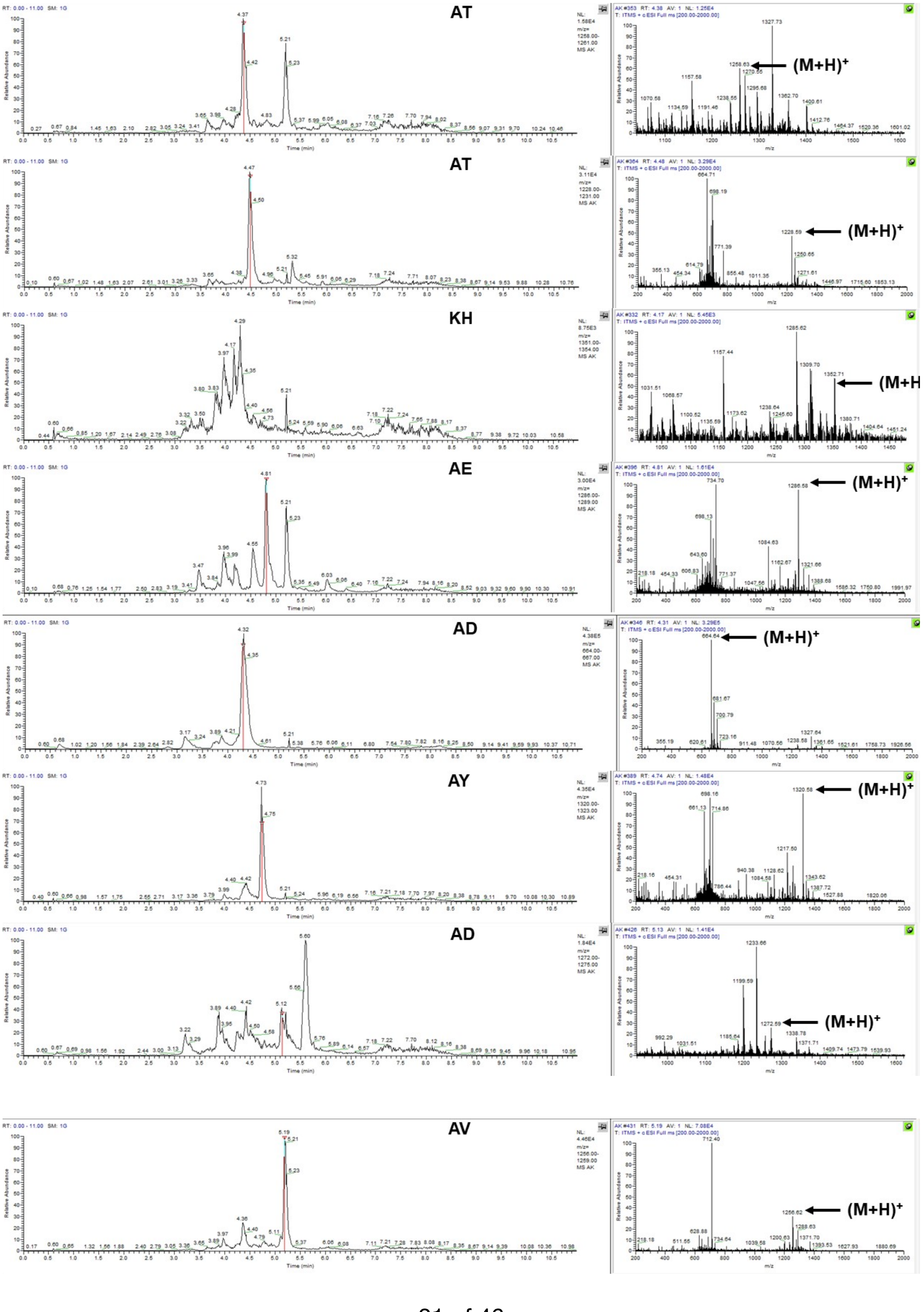
AT
(M+H)+
AT
(M+H)+
KH
(M+H
AE
(M+H)+
AD
(M+H)+
AY
(M+H)+
AD
(M+H)+
AV
(M+H)+

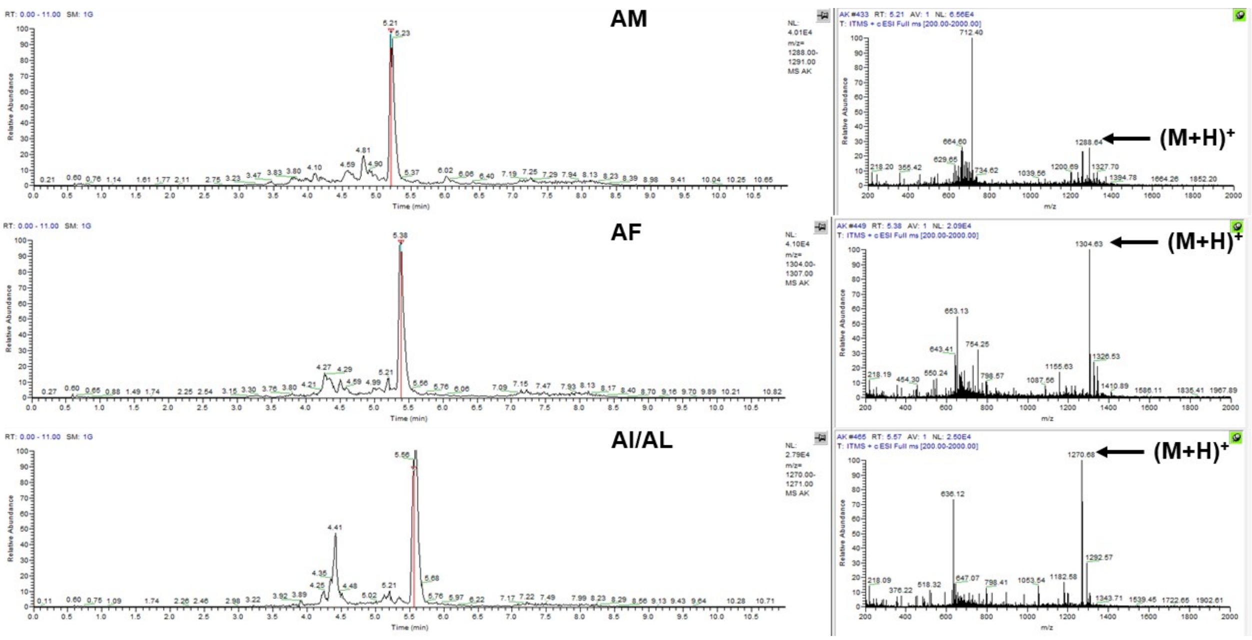
AM
(M+H)+
AF
(M+H)+
AI/AL
(M+H)+

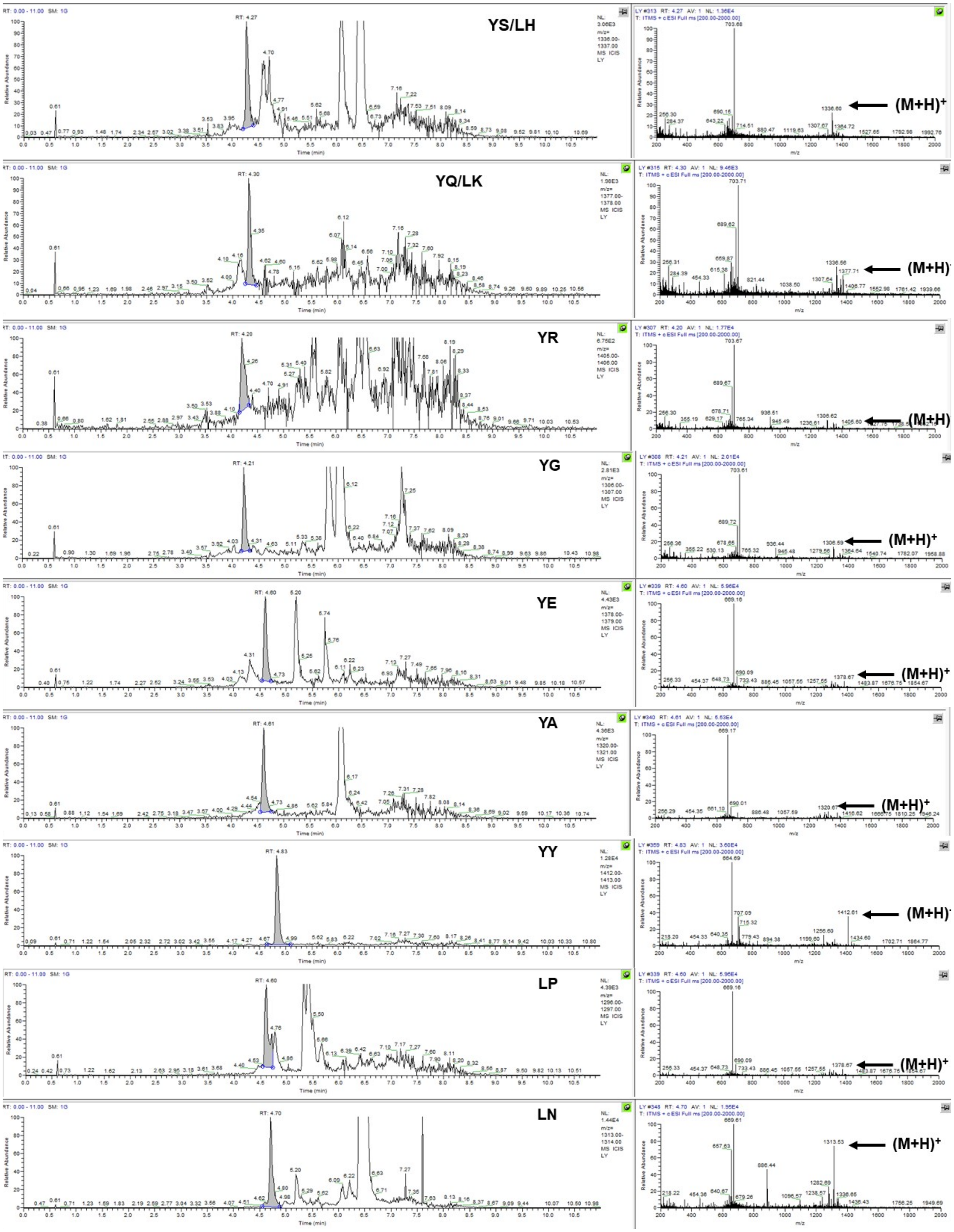
YS/LH
(M+H)+
YQ/LK
(M+H)+
YR
(M+H)+
YG
(M+H)+
YE
(M+H)+
YA
(M+H)+
YY
(M+H)+
LP
(M+H)+
LN
(M+H)+

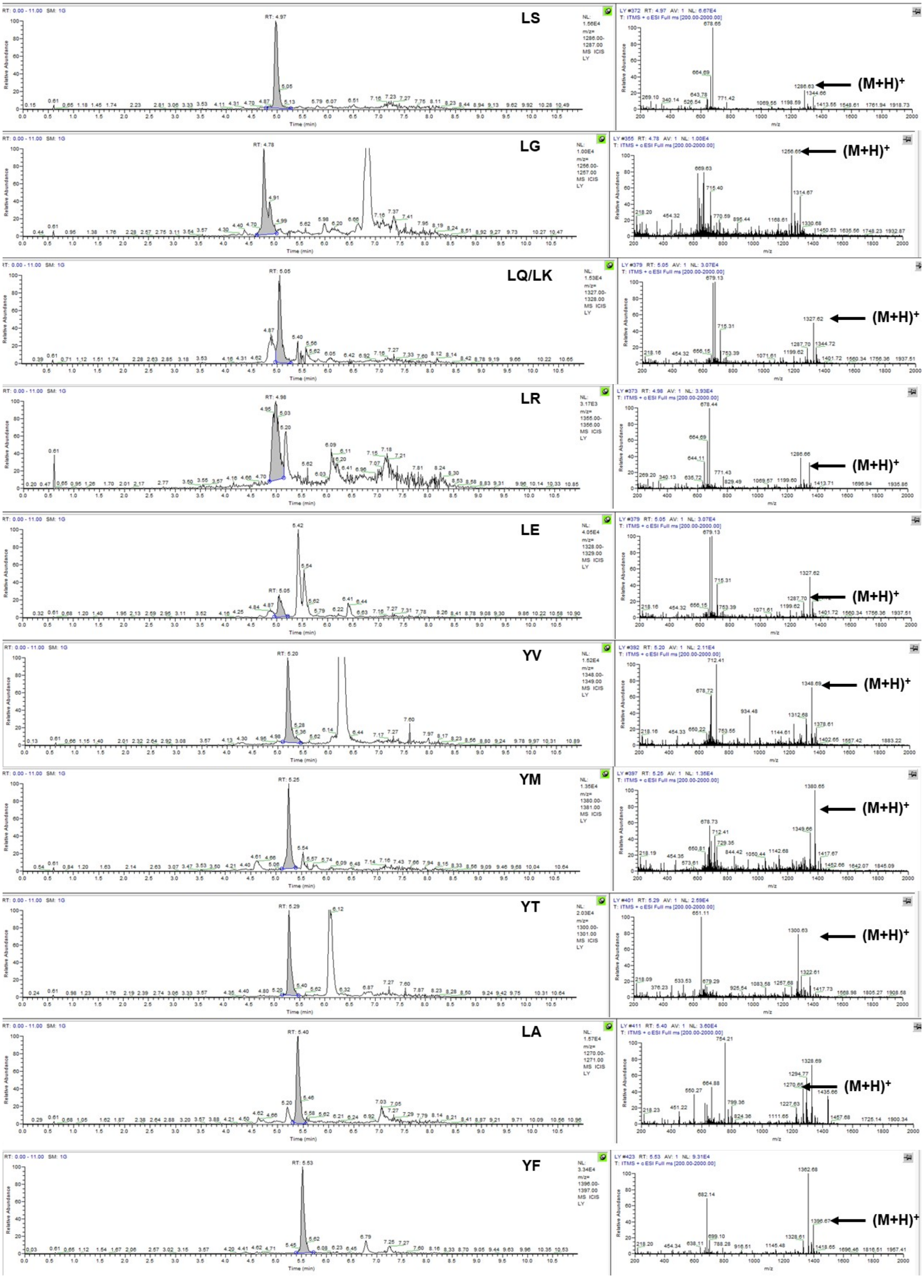
LS
(M+H)+
LG
(M+H)+
LQ/LK
(M+H)+
LR
(M+H)+
LE
(M+H)+
YV
(M+H)+
YM
(M+H)+
YT
(M+H)+
LA
(M+H)+
YF
(M+H)+

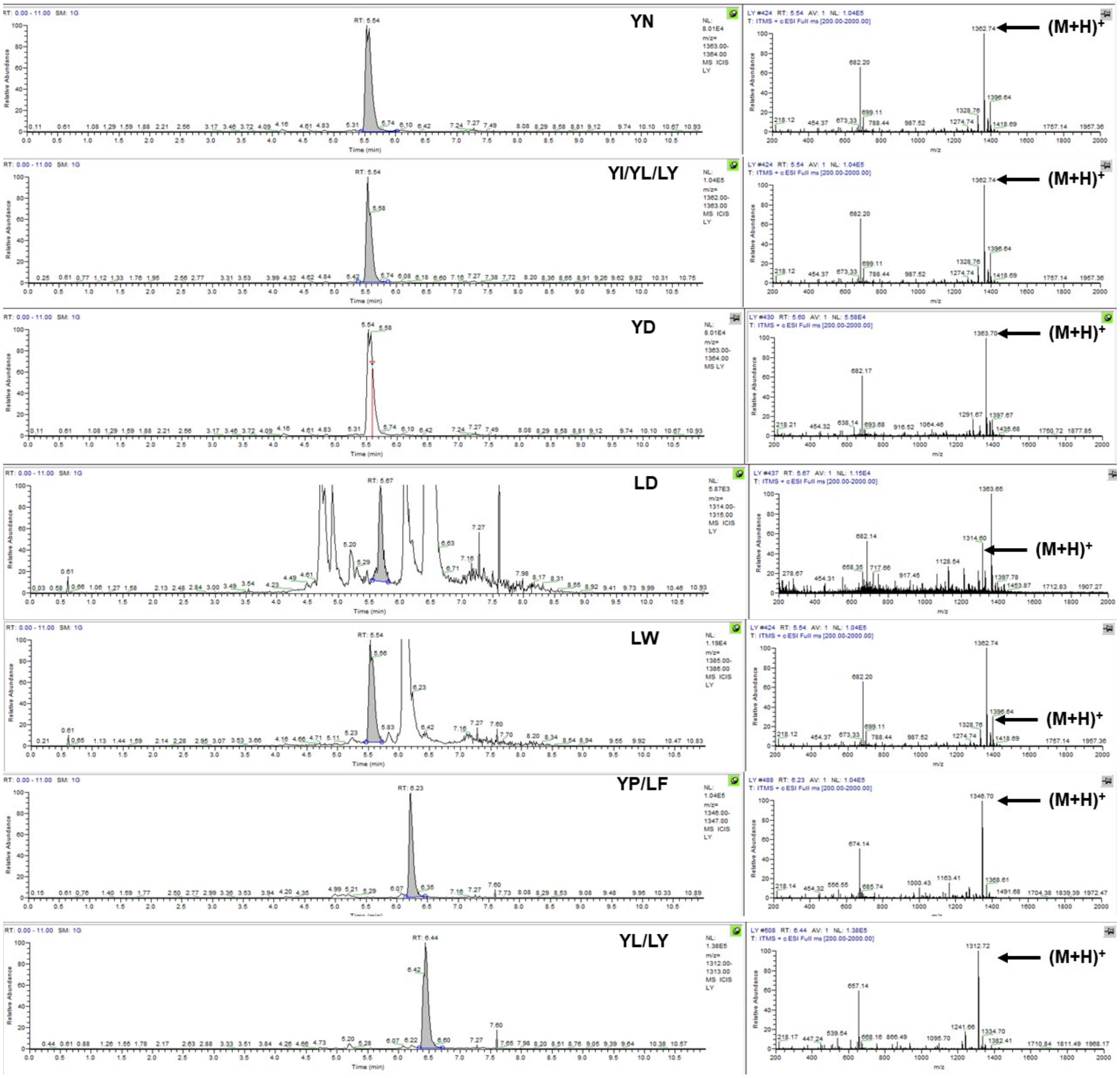
YN
(M+H)+
YI/YL/LY
(M+H)+
YD
(M+H)+
LD
(M+H)+
LW
(M+H)+
YP/LF
(M+H)+
YL/LY
(M+H)+

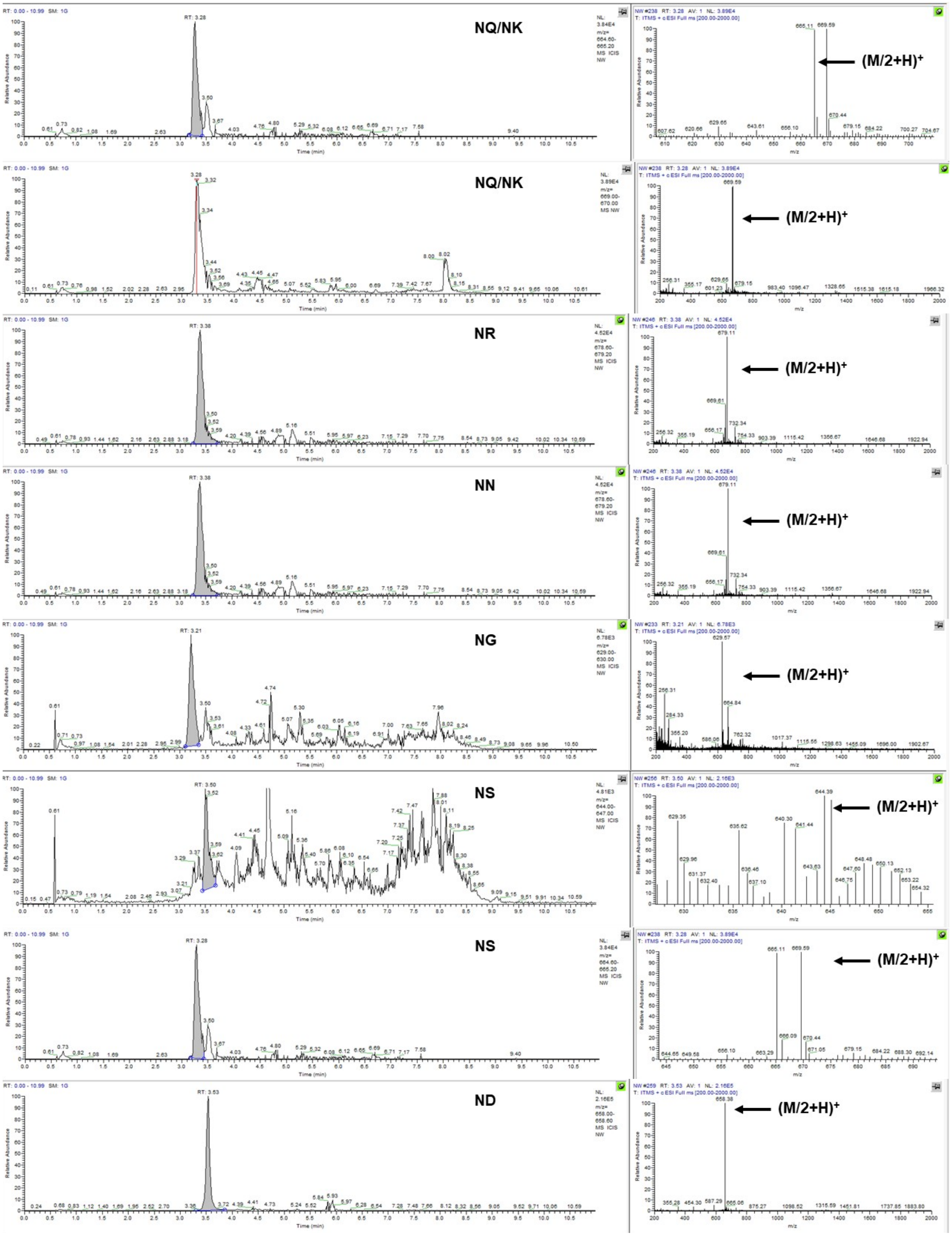
NQ/NK
(M/2+H)+
NQ/NK
(M/2+H)+
NR
(M/2+H)+
NN
(M/2+H)+
NG
(M/2+H)+
NS
(M/2+H)+
NS
(M/2+H)+
ND
(M/2+H)+

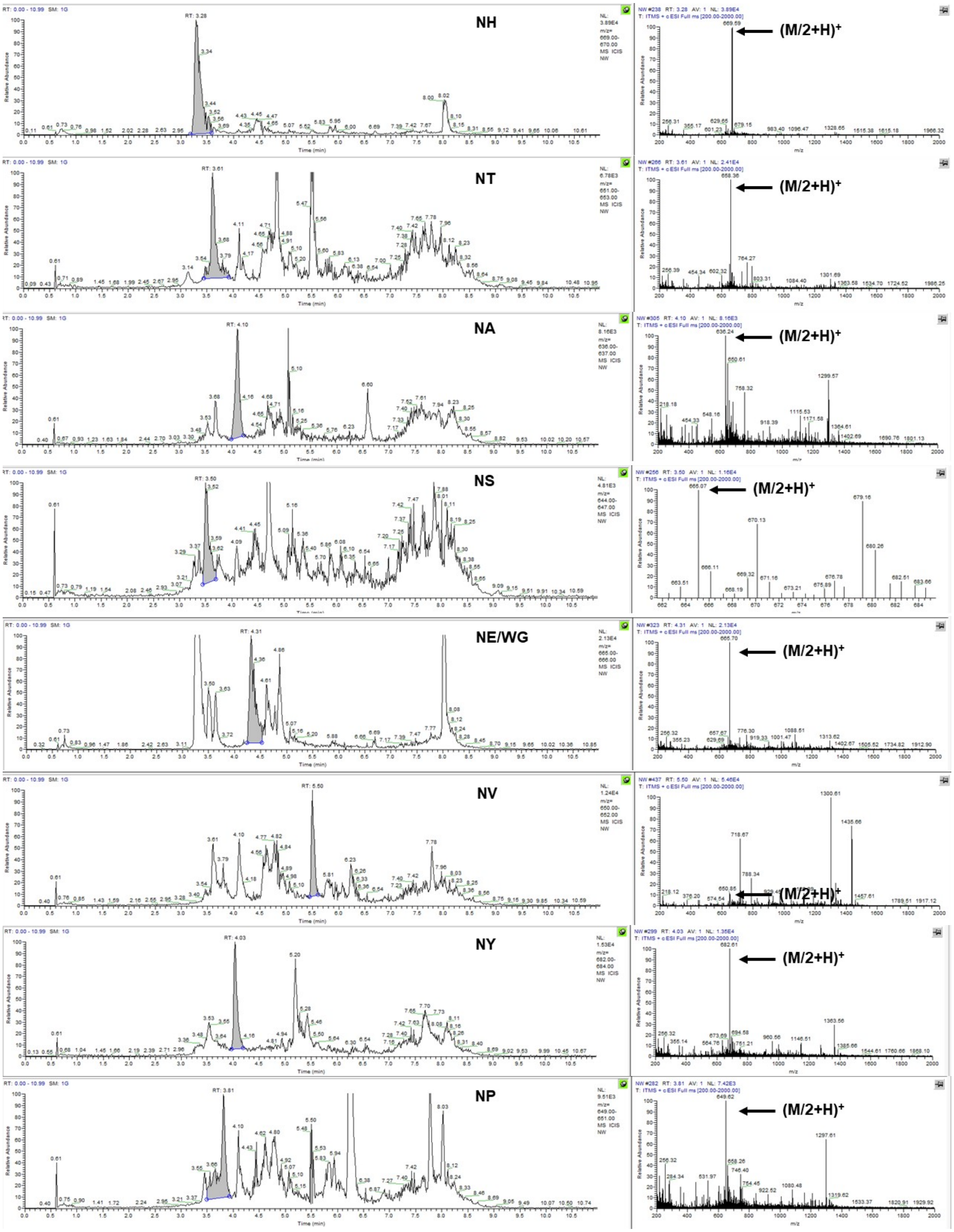
NH
(M/2+H)+
NT
(M/2+H)+
NA
(M/2+H)+
NS
(M/2+H)+
NE/WG
(M/2+H)+
NV
(M/2+H)+
NY
(M/2+H)+
NP
(M/2+H)+

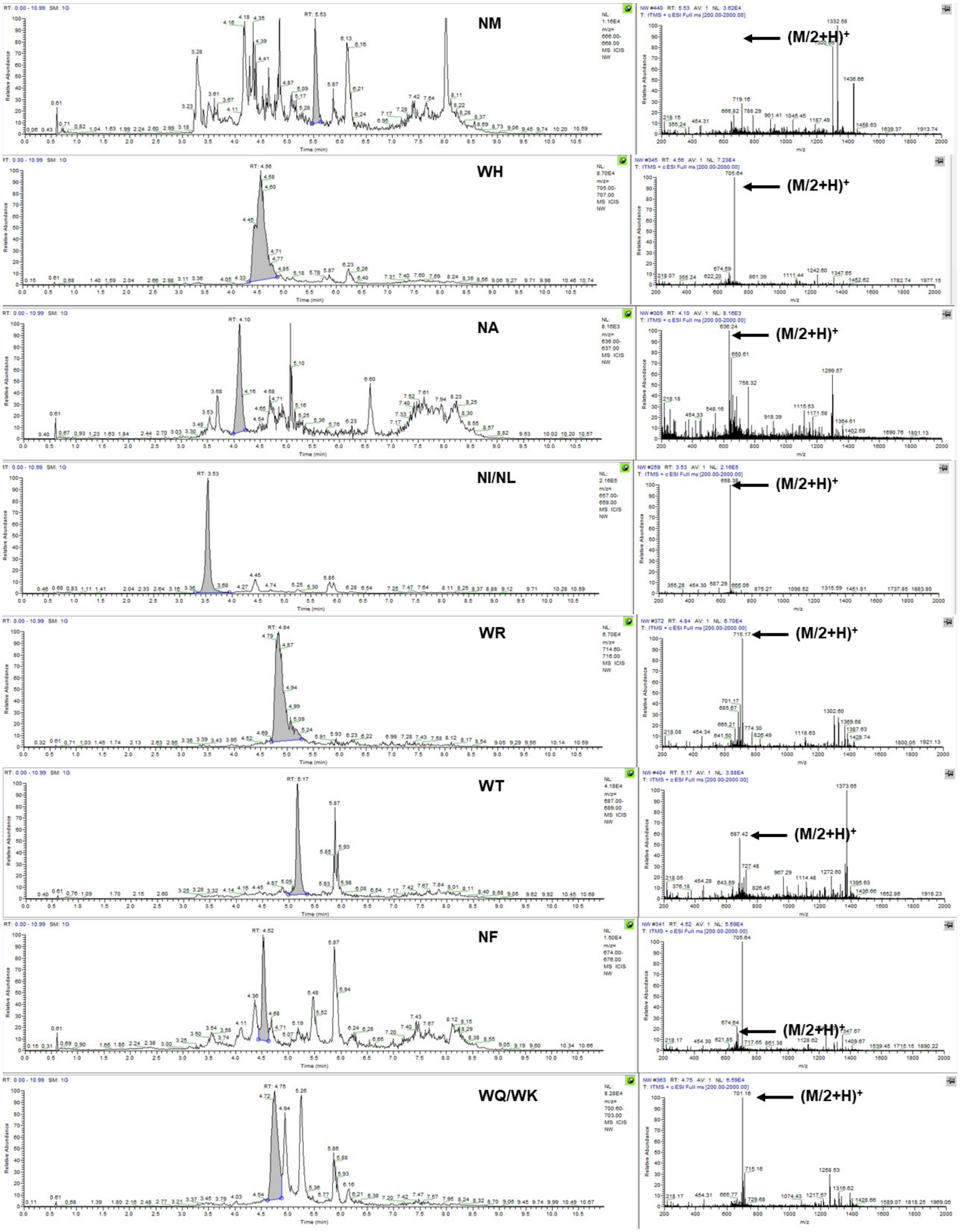
NM
(M/2+H)+
WH
(M/2+H)+
NA
(M/2+H)+
NI/NL
(M/2+H)+
WR
(M/2+H)+
WT
(M/2+H)+
NF
(M/2+H)+
WQ/WK
(M/2+H)+
Relative Abundance
Time (min)
m/z

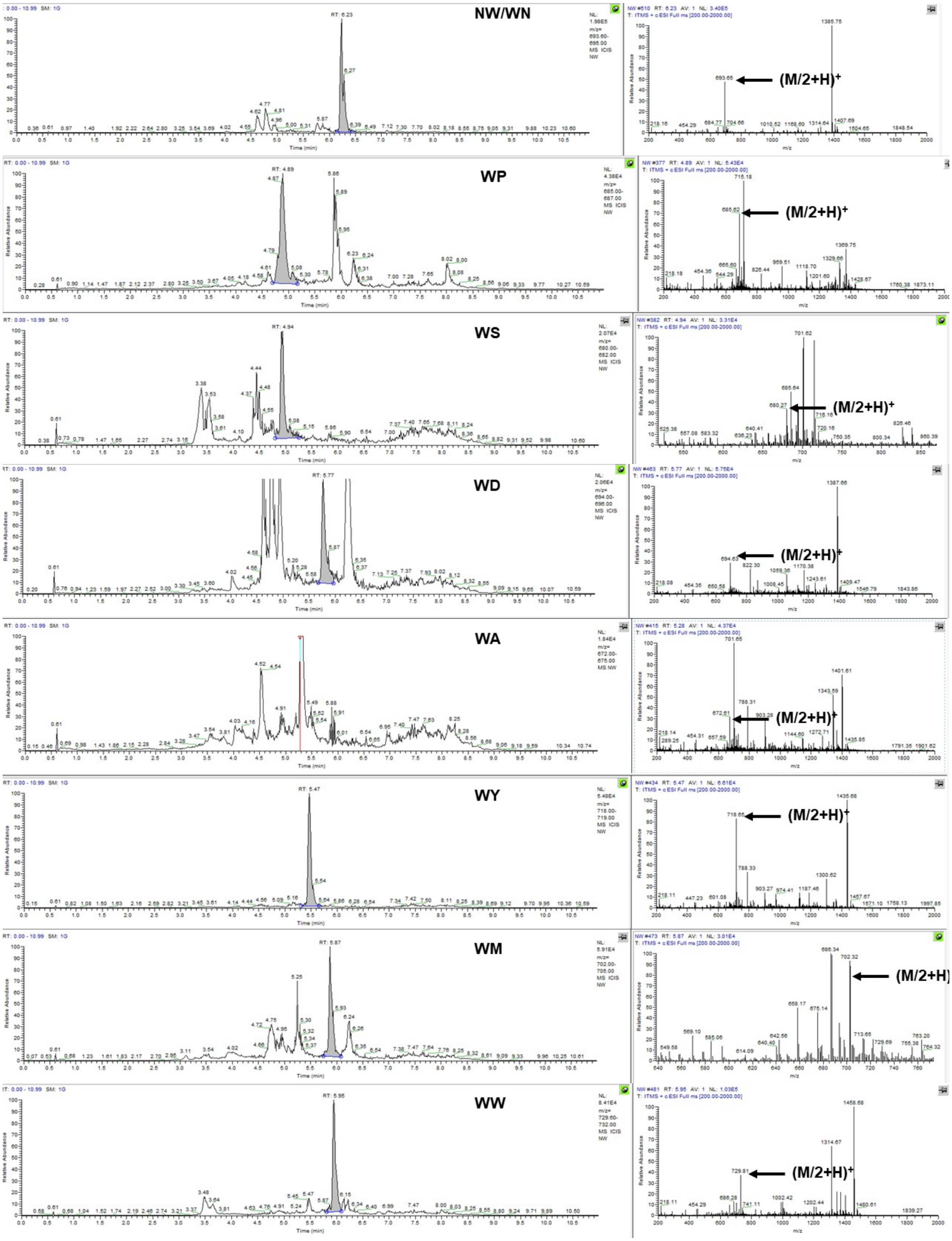
NW/WN
(M/2+H)+
WP
(M/2+H)+
WS
(M/2+H)+
WD
(M/2+H)+
WA
(M/2+H)+
WY
(M/2+H)+
WM
(M/2+H)
WW
(M/2+H)+

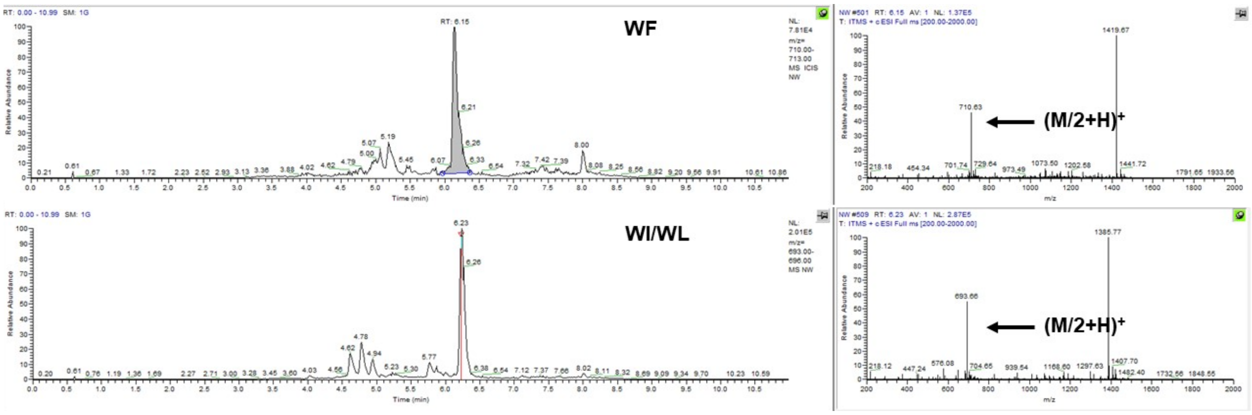
WF
WI/WL
(M/2+H)+
(M/2+H)+
Time (min)
m/z
Relative Abundance

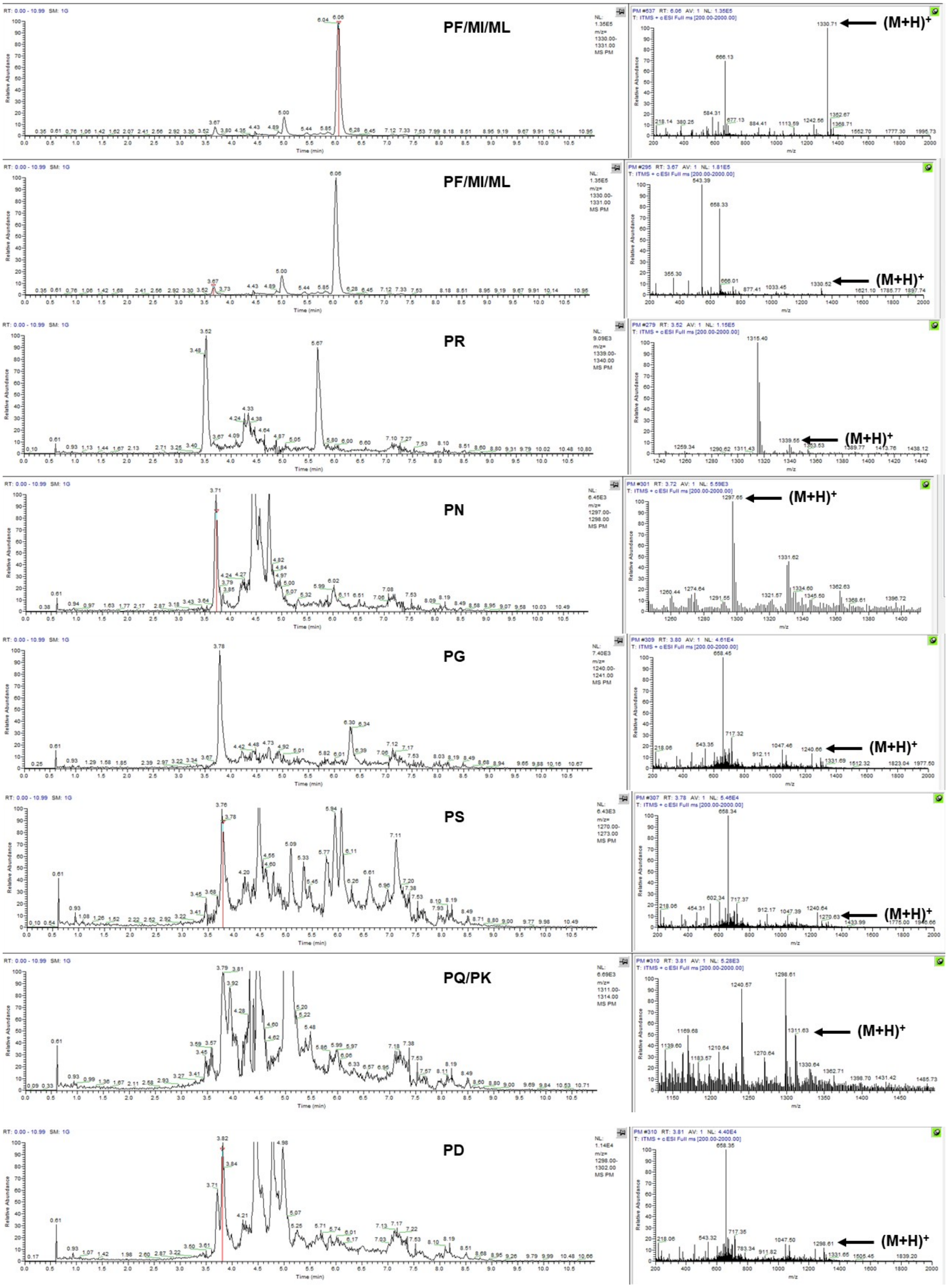
PF/MI/ML
(M+H)+
PF/MI/ML
(M+H)+
PR
(M+H)+
PN
(M+H)+
PG
(M+H)+
PS
(M+H)+
PQ/PK
(M+H)+
PD
(M+H)+

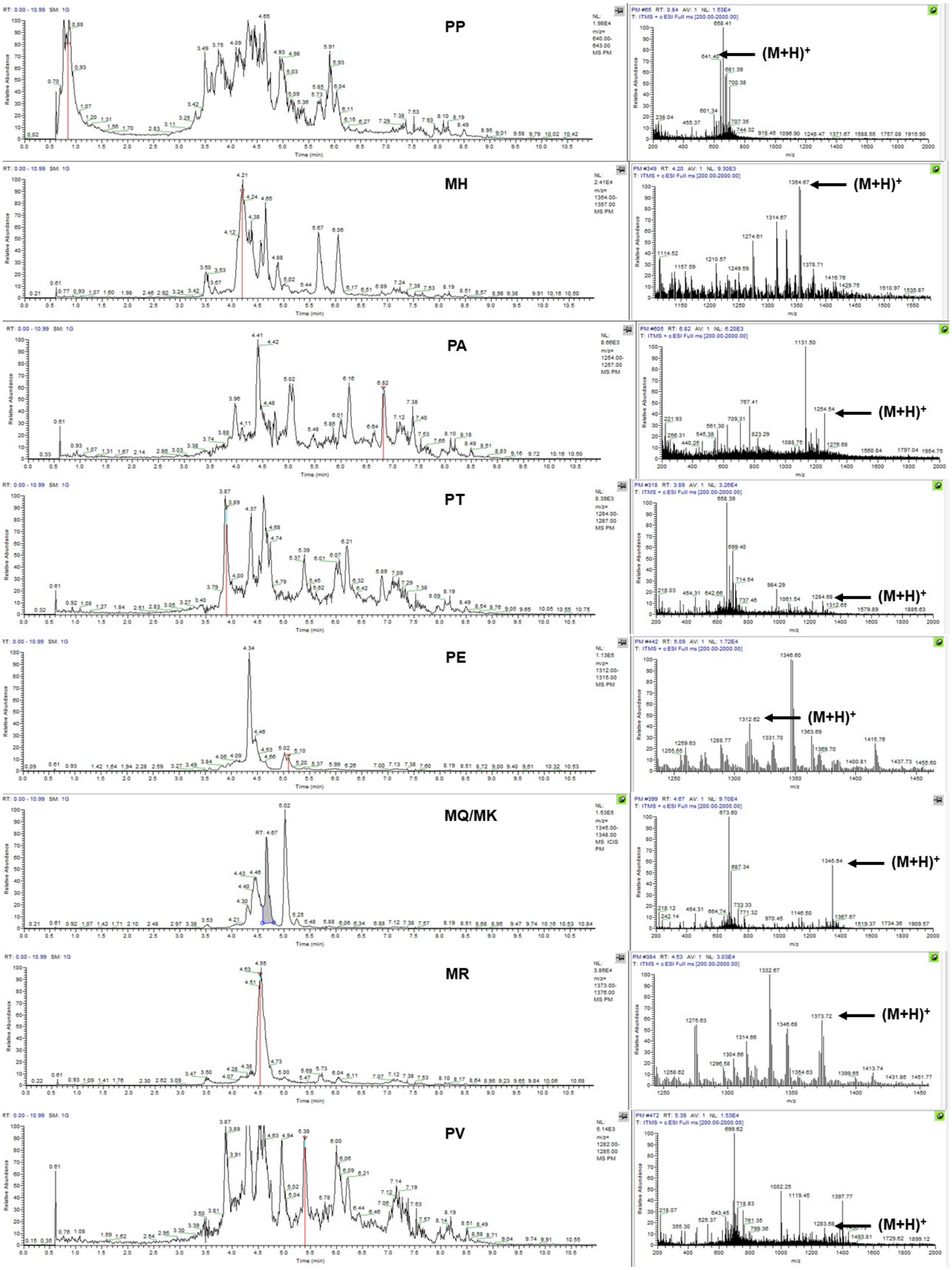
PP
(M+H)+
MH
(M+H)+
PA
(M+H)+
PT
(M+H)+
PE
(M+H)+
MQ/MK
(M+H)+
MR
(M+H)+
PV
(M+H)+

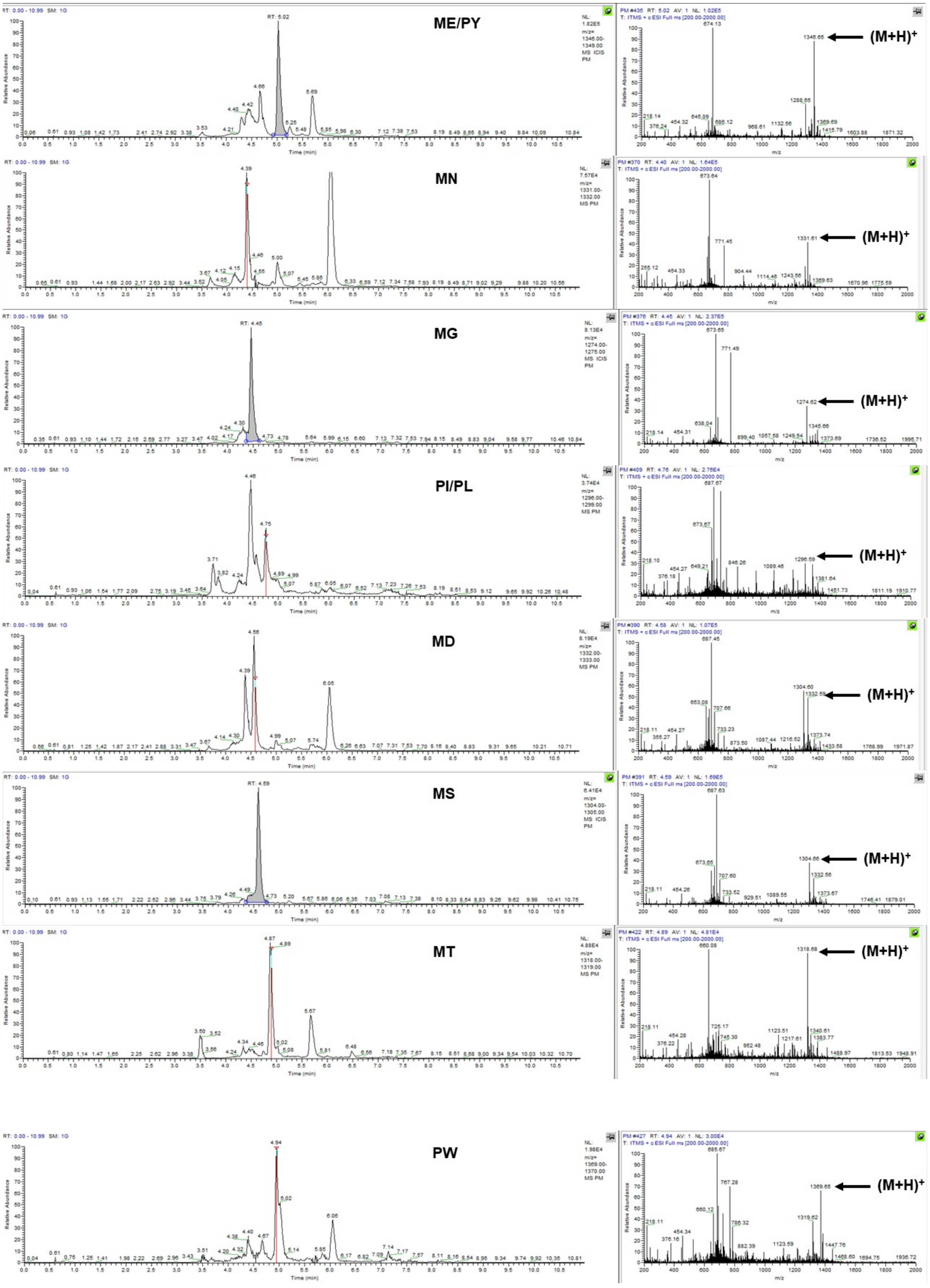
ME/PY
(M+H)+
MN
(M+H)+
MG
(M+H)+
PI/PL
(M+H)+
MD
(M+H)+
MS
(M+H)+
MT
(M+H)+
PW
(M+H)+

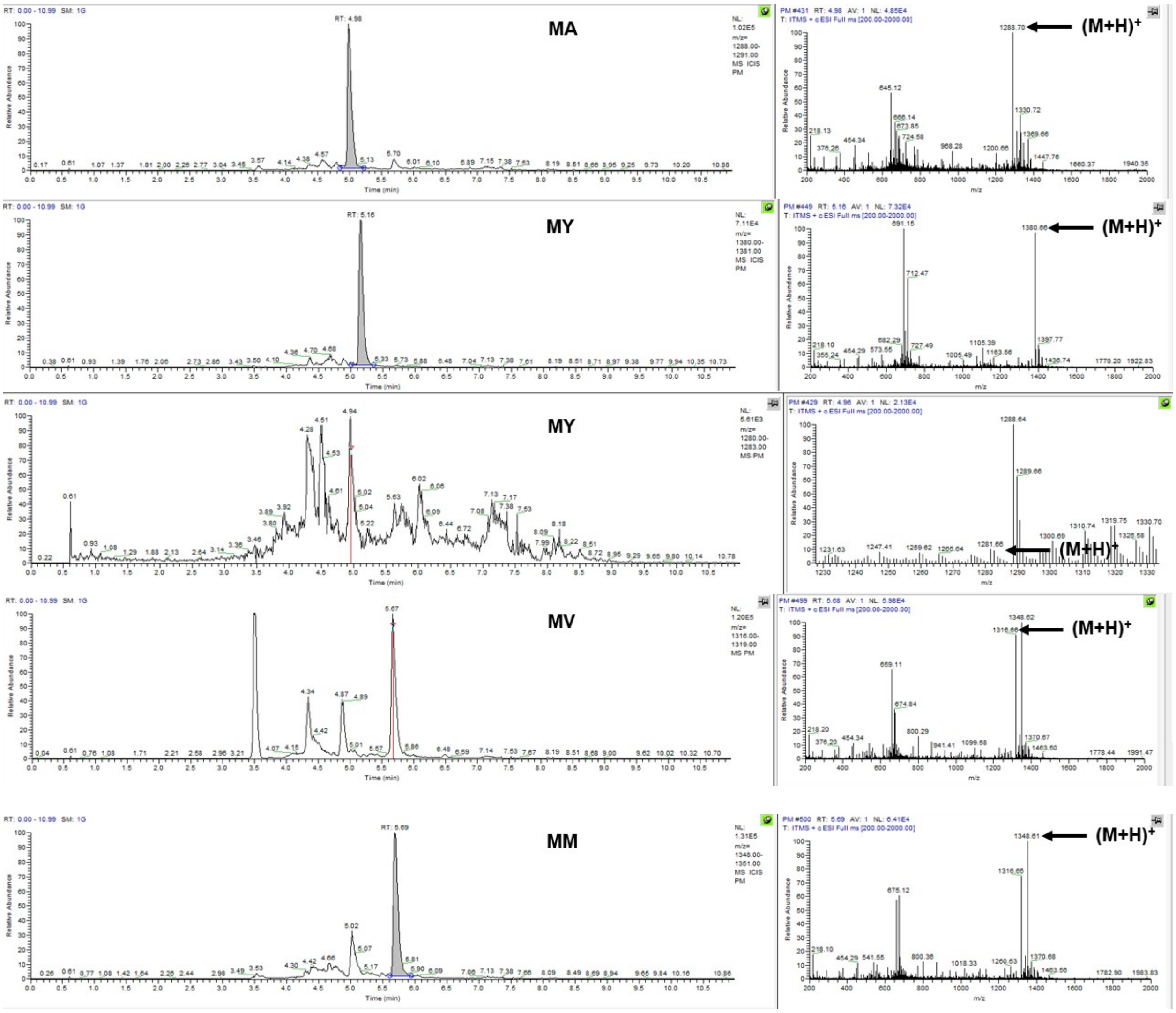
MA
(M+H)+
MY
(M+H)+
MY
(M+H)+
MV
(M+H)+
MM
(M+H)+

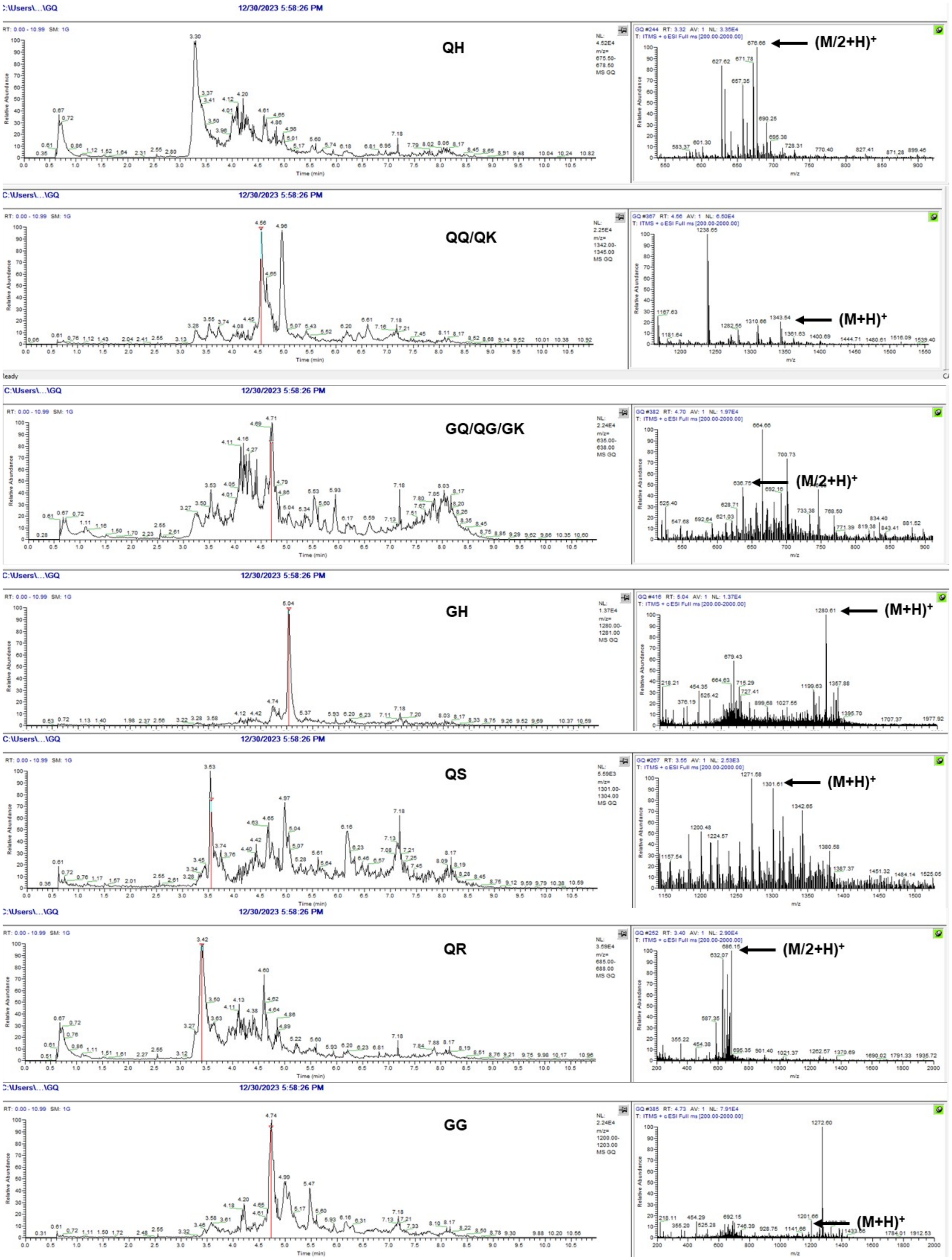
C:\Users\...\GQ
12/30/2023 5:58:26 PM
QH
(M/2+H)+
QQ/QK
(M+H)+
GQ/QG/GK
(M/2+H)+
GH
(M+H)+
QS
(M+H)+
QR
(M/2+H)+
GG
(M+H)+
Relative Abundance
Time (min)
m/z

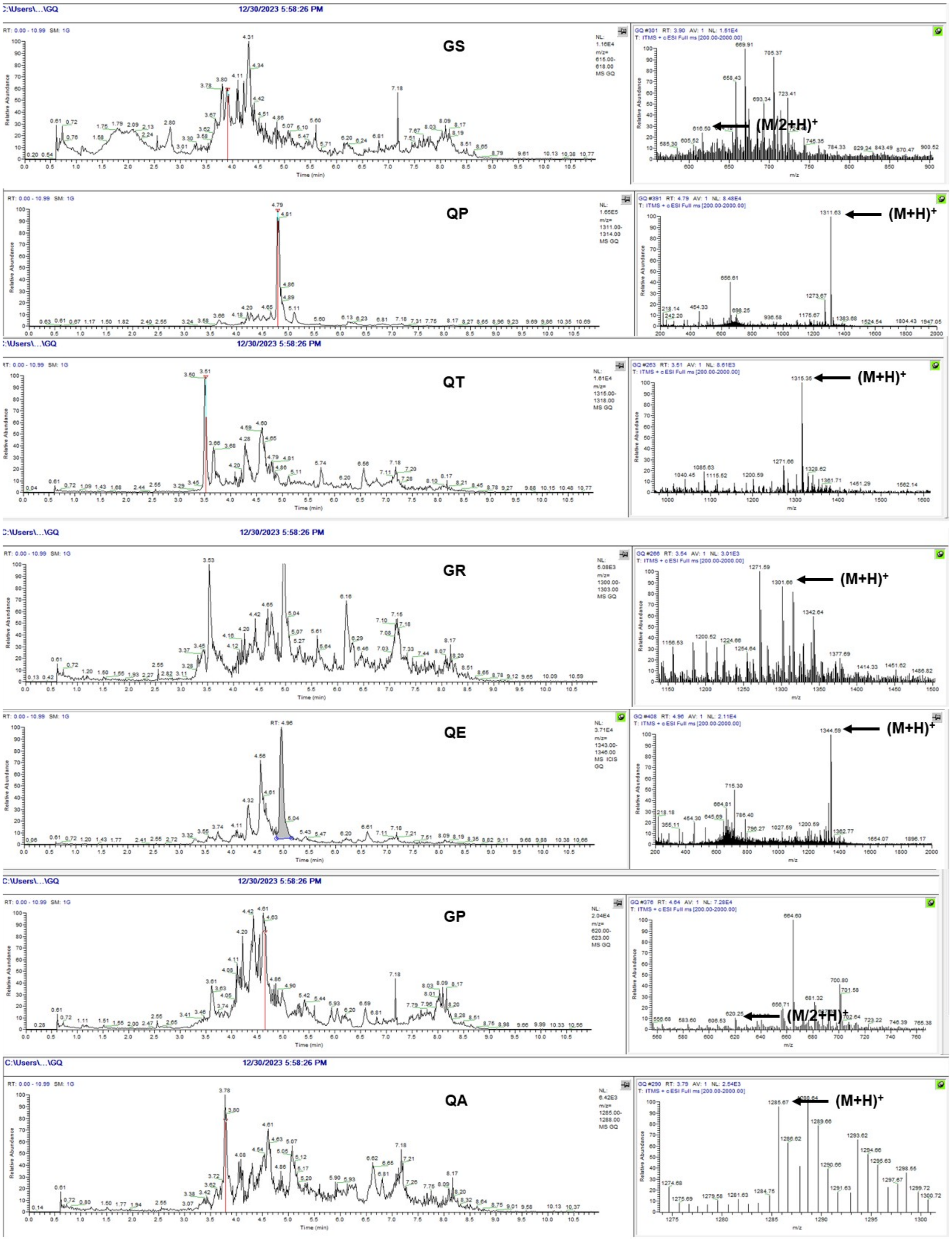
GS
(M/2+H)+
QP
(M+H)+
QT
(M+H)+
GR
(M+H)+
QE
(M+H)+
GP
(M/2+H)+
QA
(M+H)+

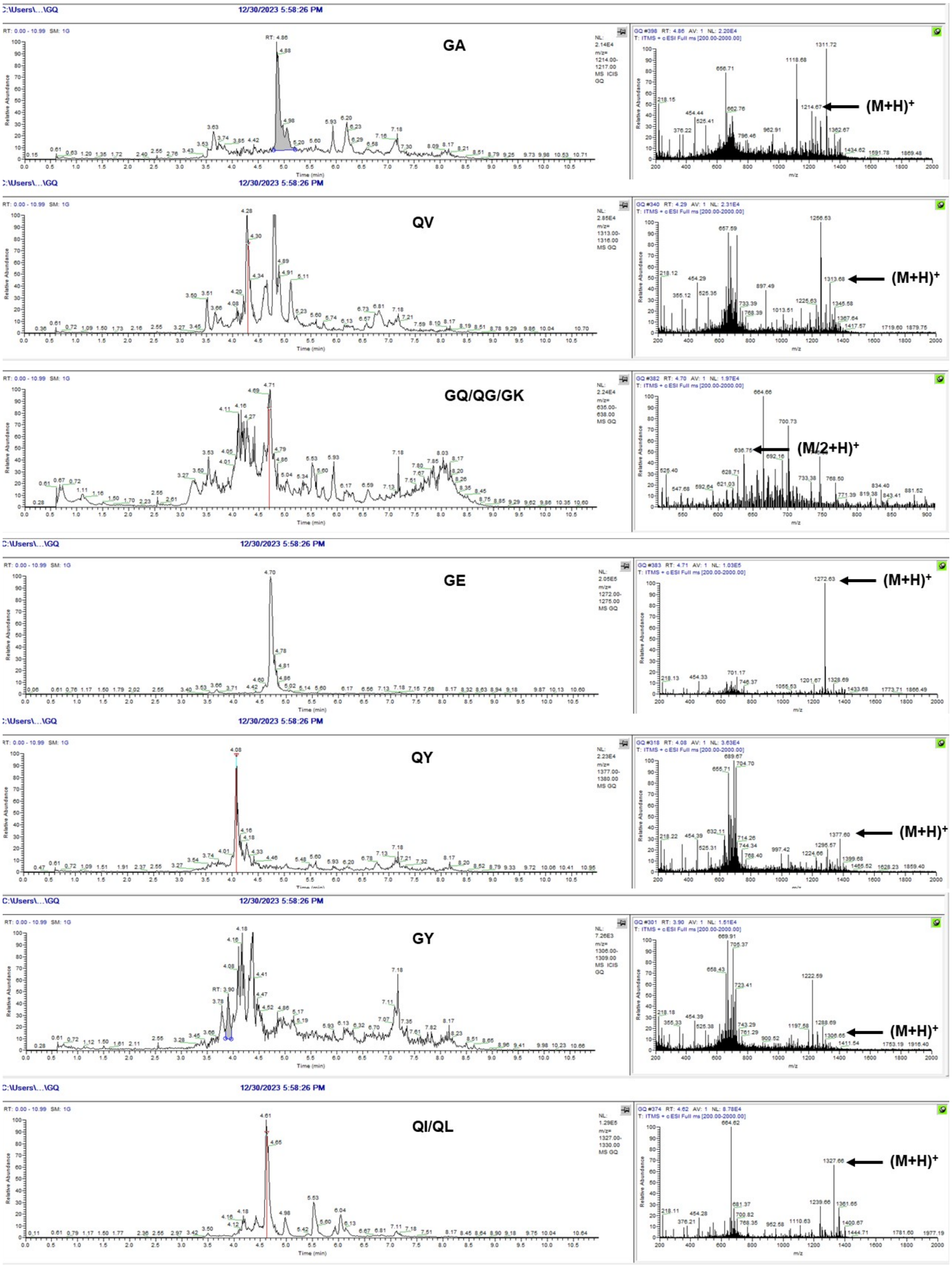
C:\Users\...\GQ
12/30/2023 5:58:26 PM
GA
(M+H)+
QV
(M+H)+
GQ/QG/GK
(M/2+H)+
GE
(M+H)+
QY
(M+H)+
GY
(M+H)+
QI/QL
(M+H)+
Time (min)
Relative Abundance
m/z

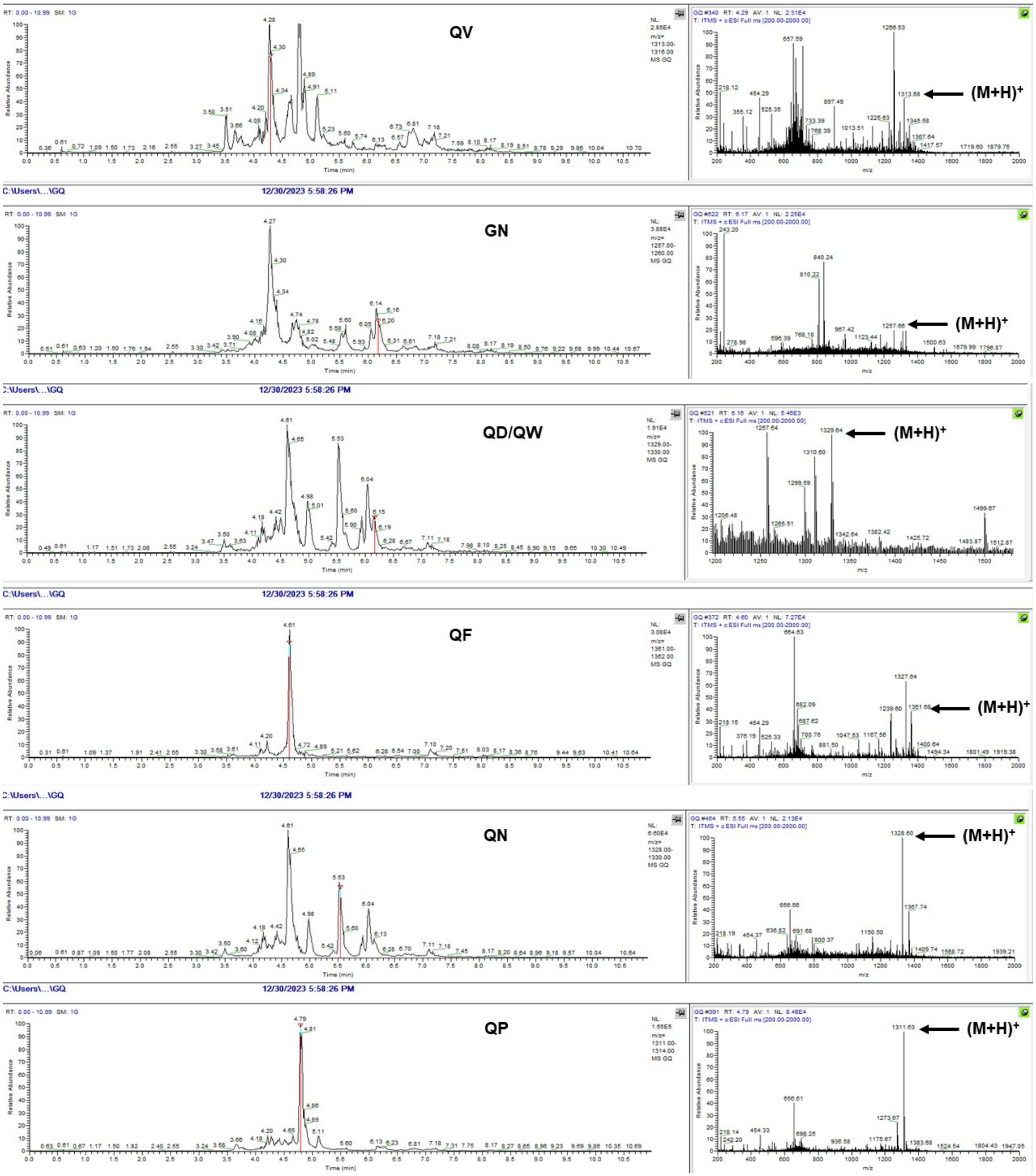
QV
(M+H)+
C:\Users\...\GQ
12/30/2023 5:58:26 PM
GN
(M+H)+
C:\Users\...\GQ
12/30/2023 5:58:26 PM
QD/QW
(M+H)+
C:\Users\...\GQ
12/30/2023 5:58:26 PM
QF
(M+H)+
C:\Users\...\GQ
12/30/2023 5:58:26 PM
QN
(M+H)+
C:\Users\...\GQ
12/30/2023 5:58:26 PM
QP
(M+H)+

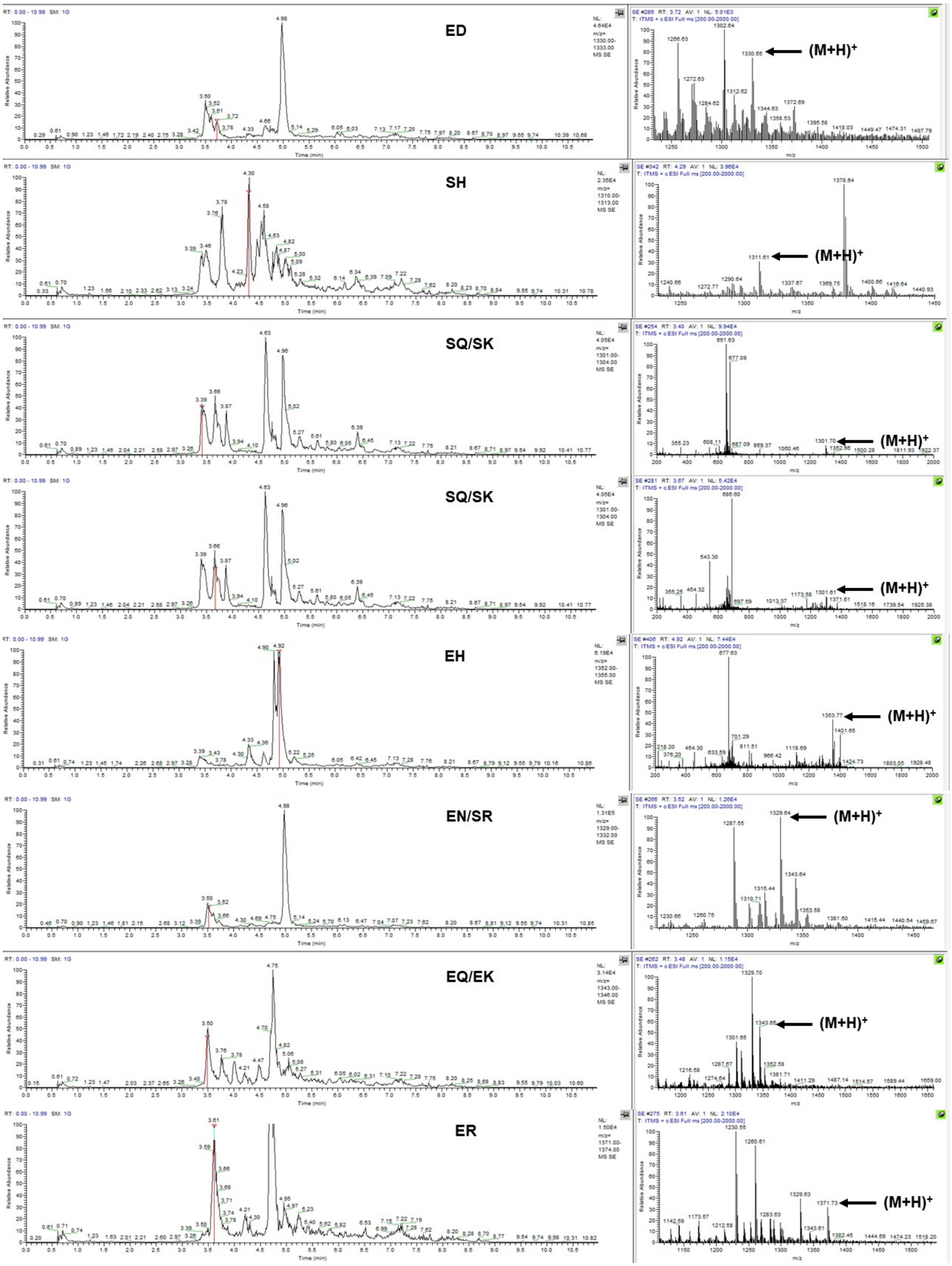
ED
(M+H)+
SH
(M+H)+
SQ/SK
(M+H)+
SQ/SK
(M+H)+
EH
(M+H)+
EN/SR
(M+H)+
EQ/EK
(M+H)+
ER
(M+H)+

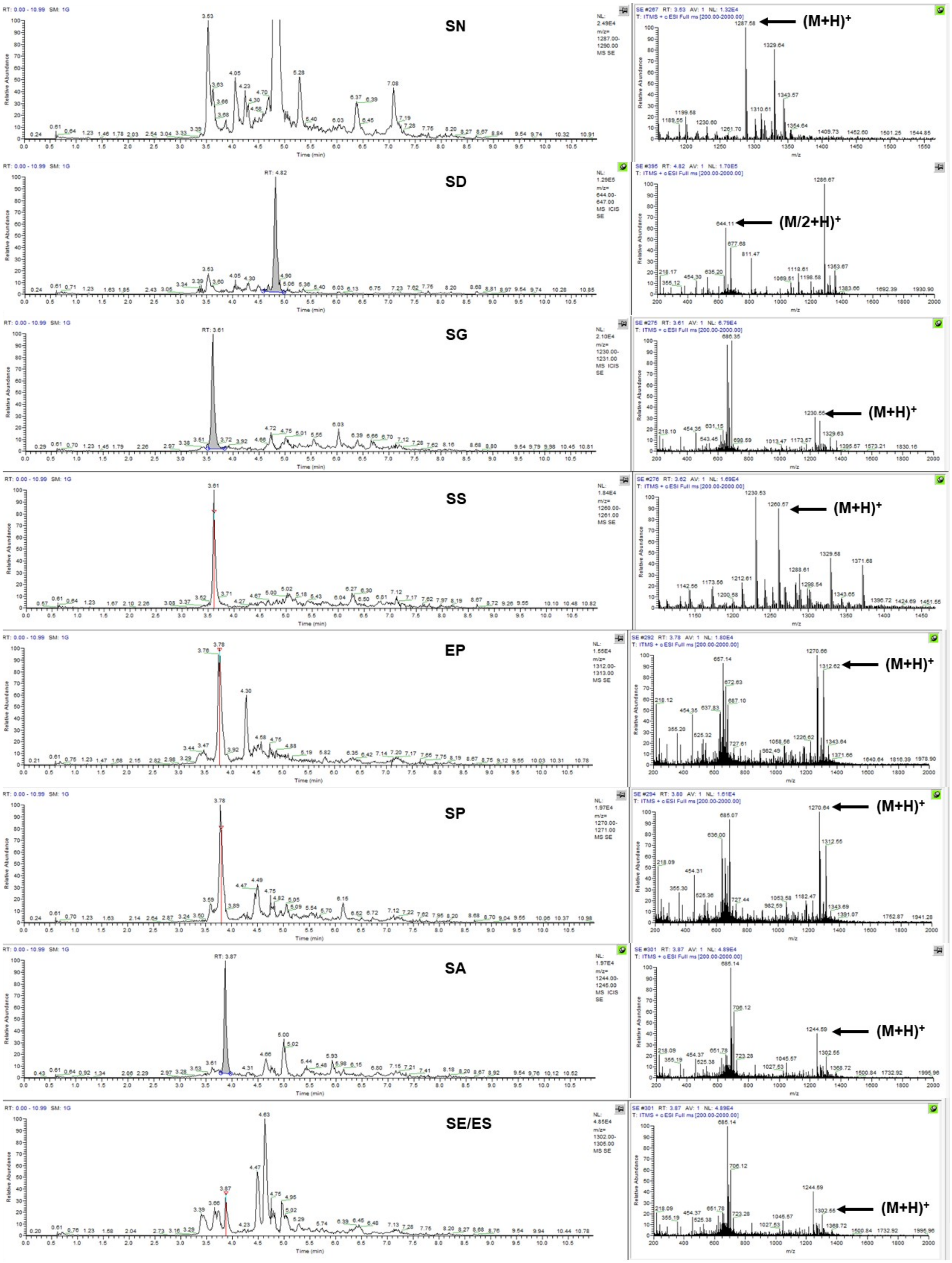
SN
(M+H)+
SD
(M/2+H)+
SG
(M+H)+
SS
(M+H)+
EP
(M+H)+
SP
(M+H)+
SA
(M+H)+
SE/ES
(M+H)+

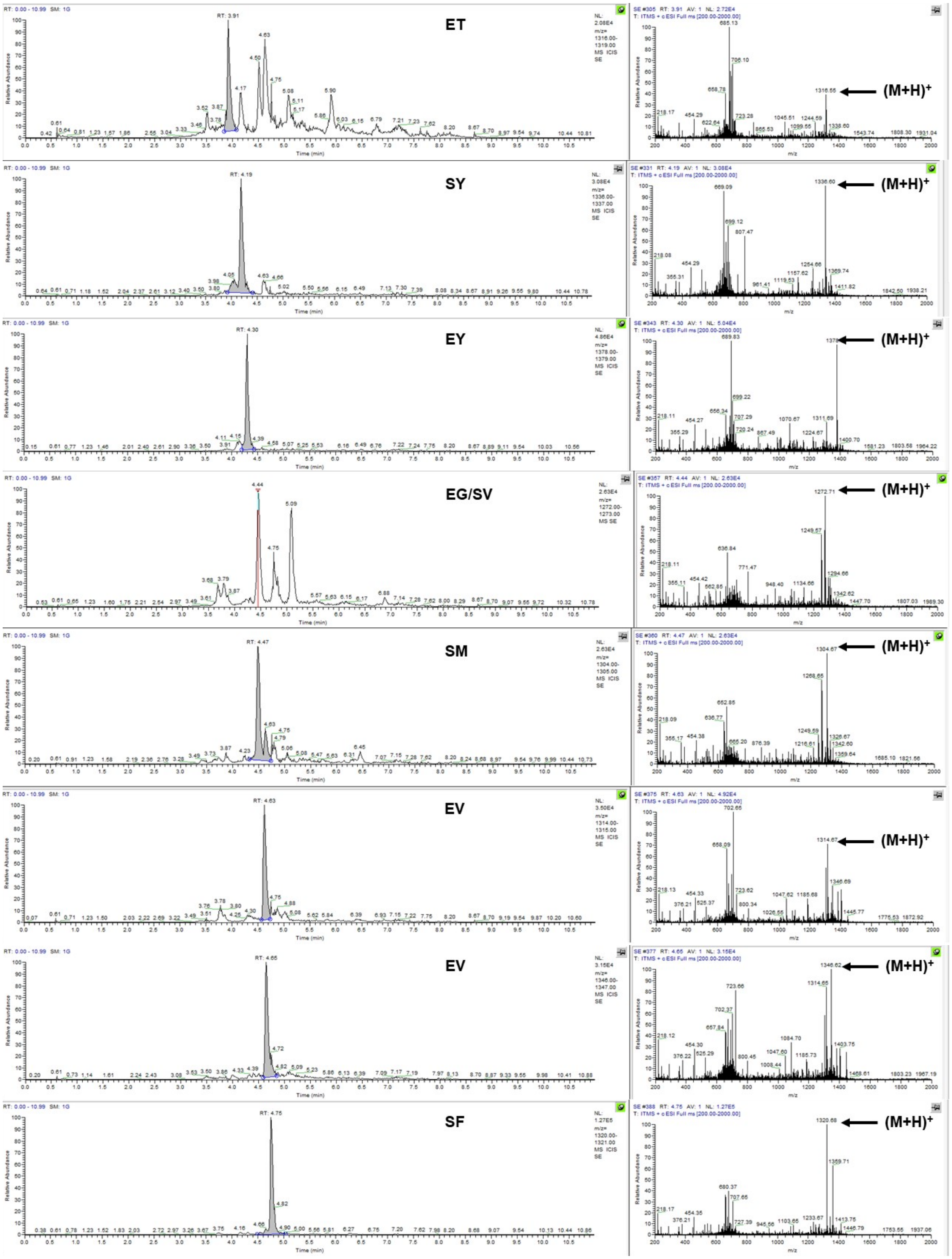
ET
(M+H)+
SY
(M+H)+
EY
(M+H)+
EG/SV
(M+H)+
SM
(M+H)+
EV
(M+H)+
EV
(M+H)+
SF
(M+H)+

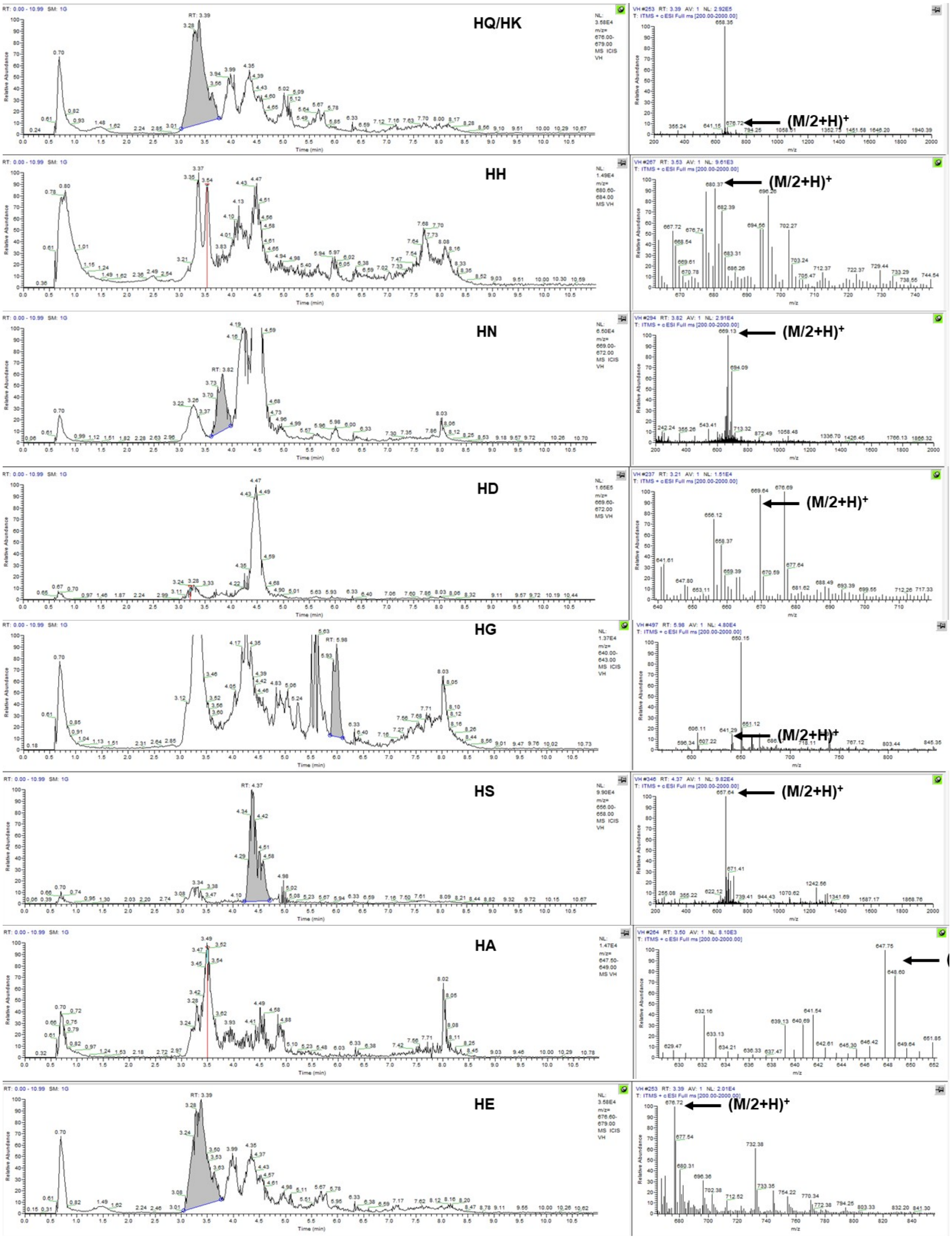
HQ/HK
(M/2+H)+
HH
(M/2+H)+
HN
(M/2+H)+
HD
(M/2+H)+
HG
(M/2+H)+
HS
(M/2+H)+
HA
HE
(M/2+H)+

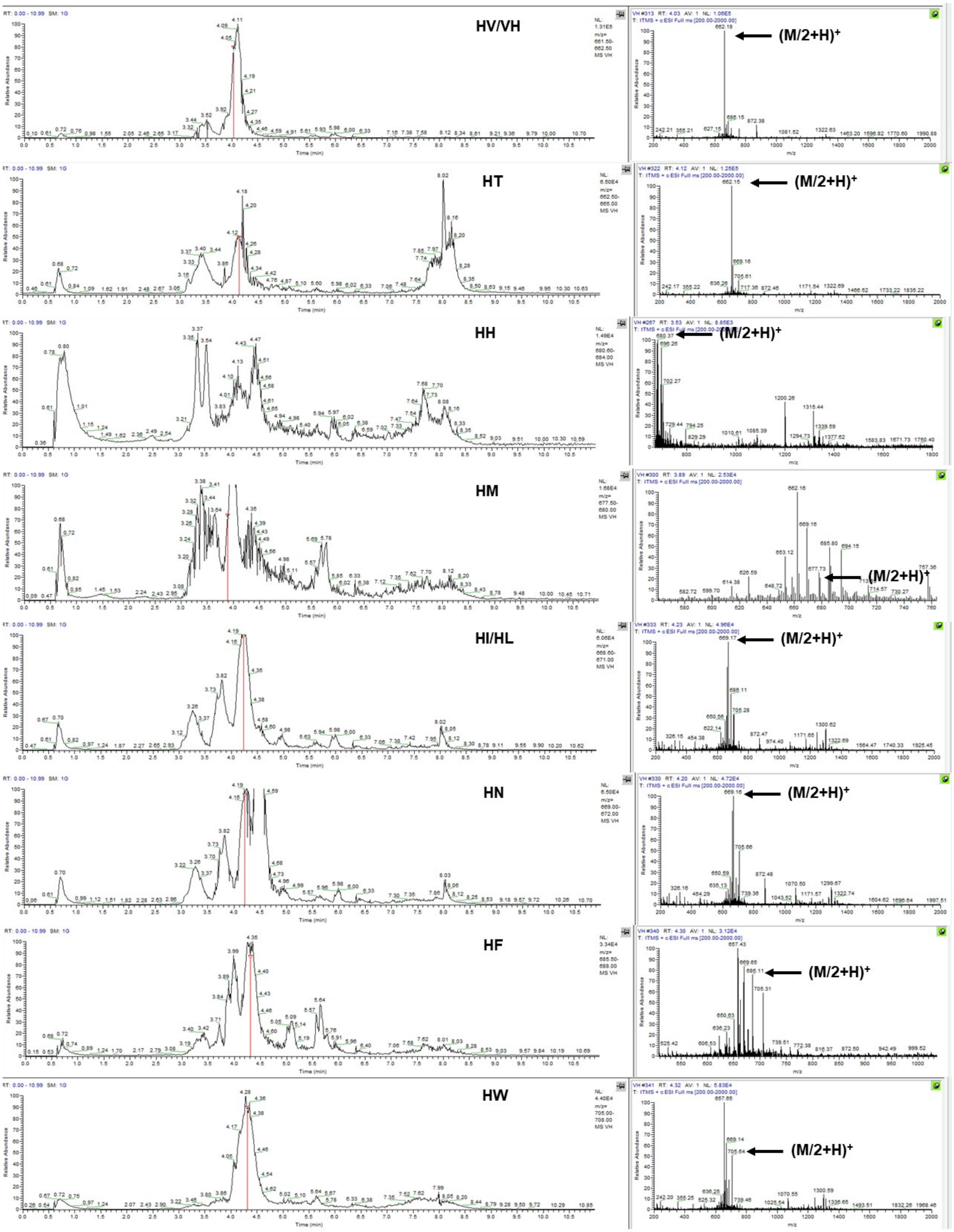
HV/VH
(M/2+H)+
HT
(M/2+H)+
HH
(M/2+H)+
HM
(M/2+H)+
HI/HL
(M/2+H)+
HN
(M/2+H)+
HF
(M/2+H)+
HW
(M/2+H)+

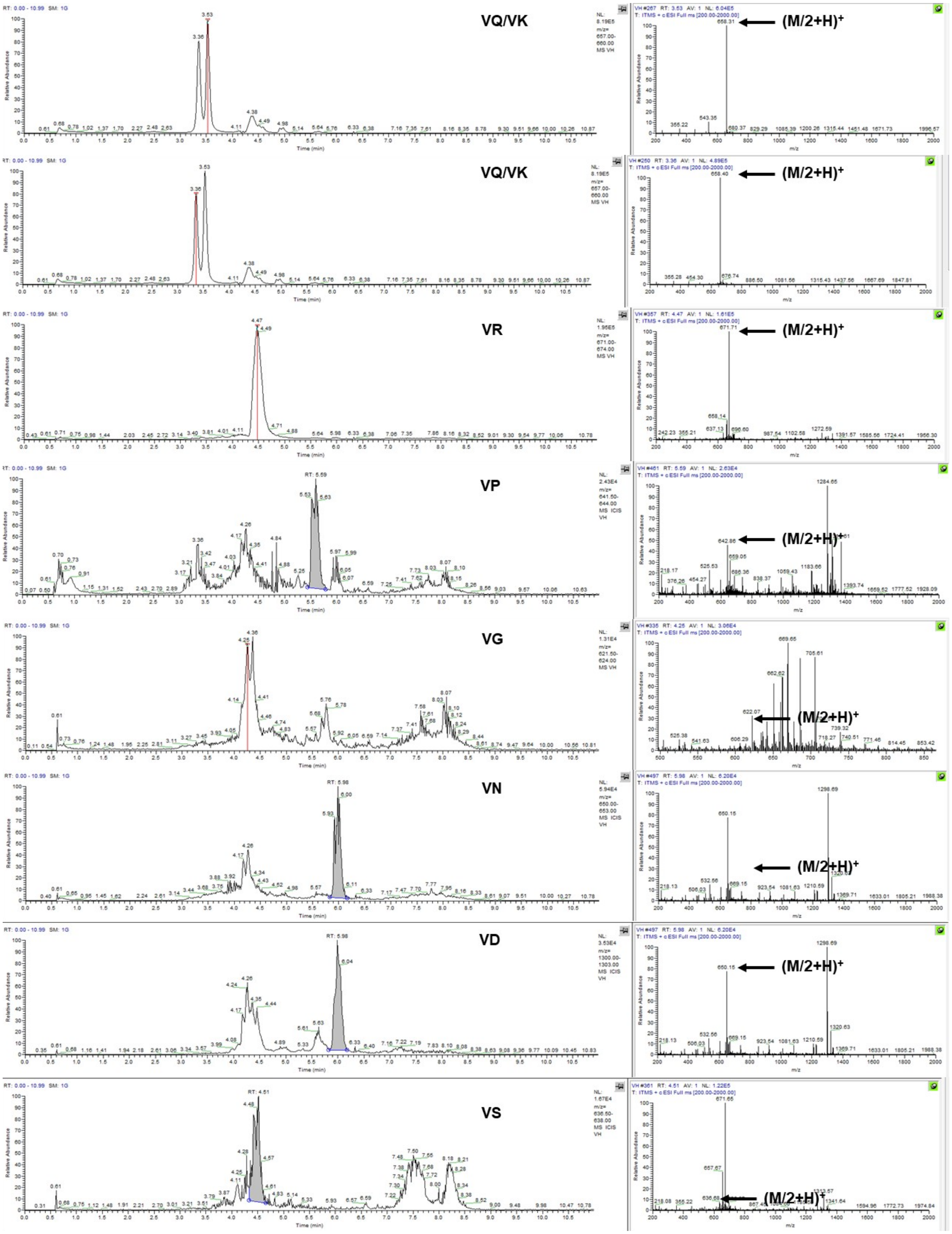
VQ/VK
(M/2+H)+
VQ/VK
(M/2+H)+
VR
(M/2+H)+
VP
(M/2+H)+
VG
(M/2+H)+
VN
(M/2+H)+
VD
(M/2+H)+
VS
(M/2+H)+

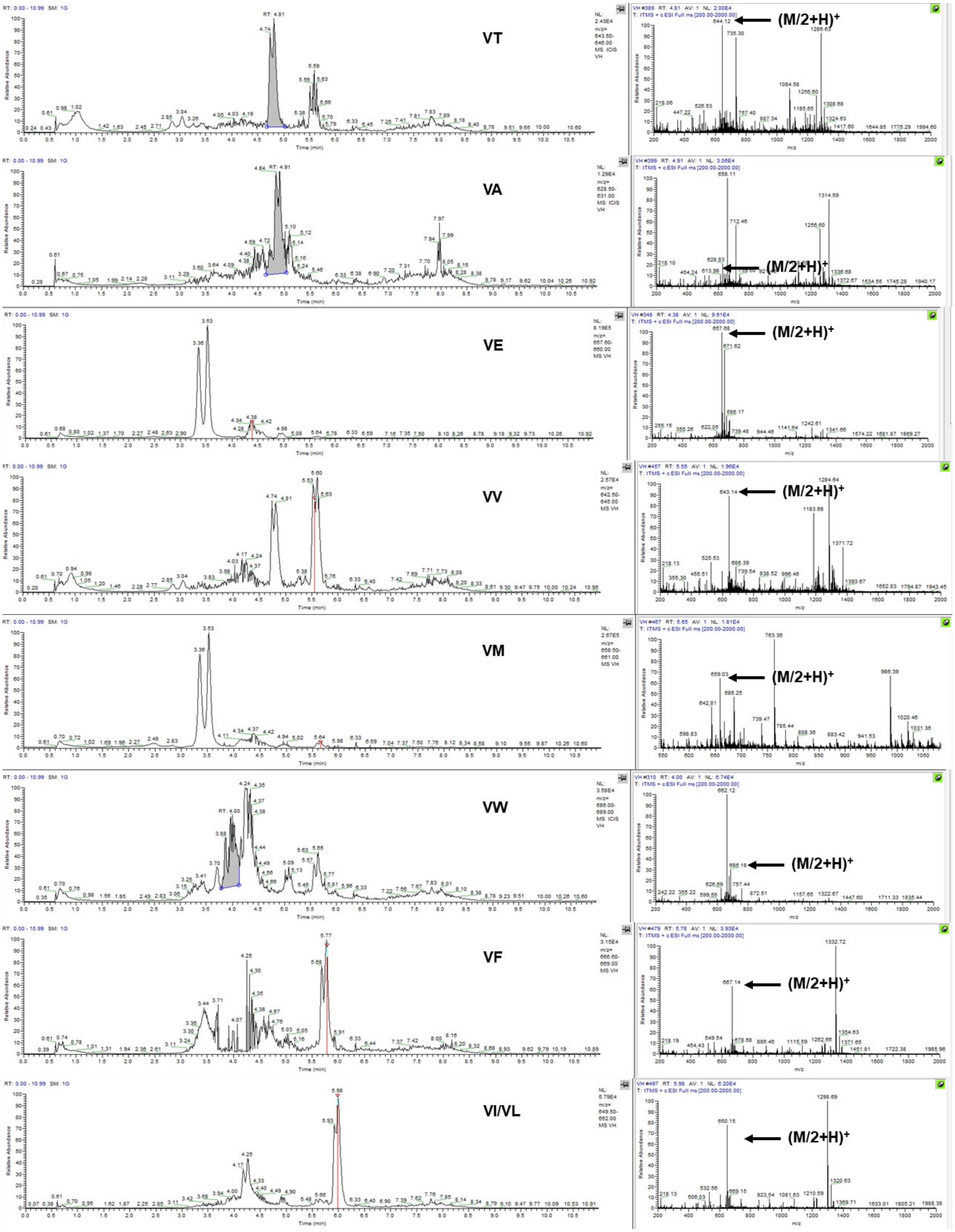


**Fig. S4.** Mass spec characterization of split-and-pool peptide libraries.
